# Intradental mechano-nociceptors serve as sentinels that prevent tooth damage

**DOI:** 10.1101/2024.05.11.593684

**Authors:** Elizabeth A. Ronan, Akash R. Gandhi, Karin H. Uchima Koecklin, Yujia Hu, Shuhao Wan, Brian S. C. Constantinescu, Mak E. Guenther, Maximilian Nagel, Ling-Yu Liu, Aditi Jha, Leen Dakhilalla, Kaitlyn J. Blumberg, Isaac T. Berthaume, Tomer Stern, Kevin P. Pipe, Bing Ye, Peng Li, Joshua J. Emrick

**Affiliations:** Department of Biologic and Materials Sciences & Prosthodontics, School of Dentistry, University of Michigan, Ann Arbor, MI, USA; Life Sciences Institute, University of Michigan, Ann Arbor, MI, USA; Department of Molecular and Integrative Physiology, University of Michigan Medical School, Ann Arbor, MI, USA; Department of Mechanical Engineering, University of Michigan, Ann Arbor, MI, USA; Sensory Cells and Circuits Section, National Center for Complementary and Integrative Health, Bethesda, MD, USA; Department of Cell and Developmental Biology, University of Michigan, Ann Arbor, MI, USA

## Abstract

Pain is the anticipated output of the trigeminal sensory neurons that innervate the tooth’s vital interior^1,2^; however, the contribution of intradental neurons to healthy tooth sensation has yet to be defined. Here, we employ in vivo Ca^2+^ imaging to identify and define a population of myelinated high-threshold mechano-nociceptors (intradental HTMRs) that detect superficial structural damage of the tooth, produce pain, and initiate a jaw opening reflex. Intradental HTMRs remain inactive when direct forces are applied to the intact tooth but become responsive to forces when the structural integrity of the tooth is compromised, and the dentin or pulp is exposed. Their terminals collectively innervate the inner dentin through overlapping receptive fields, allowing them to monitor the superficial structures of the tooth. Indeed, intradental HTMRs detect superficial enamel damage and encode its degree, and their responses persist in the absence of either PIEZO2 or Na_v_1.8^3,4^. As predicted, chemogenetic activation of intradental HTMRs results in a marked pain phenotype like that produced by systemic chemogenetic activation of nociceptors. Remarkably, optogenetic activation of intradental HTMRs triggers a rapid, jaw opening reflex via contraction of the digastric muscle. Taken together, our data indicate that intradental HTMRs serve as sentinels that guard against mechanical threats to the tooth; their activation not only triggers pain, but also results in physical tooth separation, which would prevent damage during mastication. Our work provides a new perspective of intradental neurons, highlighting their protective role, and illustrates the functional diversity of interoreceptors.

## INTRODUCTION

Mammals instinctively use their teeth for mechanical behaviors including chewing to initiate digestion and biting for prey capture and self-defense. The protection and maintenance of enamel and teeth are essential for mammals with teeth that do not continually regenerate. In humans, loss of teeth is associated with poorer diet and nutritional status^5^ as well as early mortality^6^. The somatosensory neurons within the trigeminal ganglion extend processes into the oral and craniofacial tissues, including the inner tooth. Broadly, activation of primary sensory cells and their downstream neuronal targets produces outputs including perceptions of touch, temperature, body position, and pain and sensorimotor reflexes^7,8^. While we assume that the activation of tooth innervation is the source of substantial pain^9^, whether tooth-innervating somatosensory neurons (hereafter intradental neurons) might contribute to the protection of tooth structure in health has not been explored.

The tooth comprises external mineralized layers (i.e., outermost enamel then dentin) and an internal soft tissue (i.e., dental pulp). Historically, intradental innervation is thought to arise from both myelinated (Aβ/Aδ) and unmyelinated (C-type) trigeminal sensory neurons. However, myelinated intradental neurons predominate based on immunohistochemical labeling^10^, and transcriptomic analyses indicating the majority of intradental neurons express *S100b*, a marker of myelinated somatosensory neurons^11,12^. While electrophysiological experiments have demonstrated that fibers with conduction velocities in the Aδ range spike in extreme conditions when the inner tooth has been exposed^13^, the molecular identity of intradental neurons and their capacity to respond in an intact tooth has not been explored. The combination of transcriptomic analyses with in vivo functional imaging provides a new ability to relate molecular identity with cell function^7,11,14–18^. Indeed, these tools have revealed the basis for touch sensation in the skin^12,19^, and provide great potential for defining the role of intradental neurons.

Here, we develop a strategy to identify intradental neurons in the trigeminal ganglion without damaging the tooth, then monitor their response profiles to mechanical stimulation. We determine most responding neurons are large diameter and co-express transcriptomic markers of myelinated mechano-nociceptors (HTMRs). These intradental neurons do not encode any direct forces applied to the intact outer tooth but are activated once the dentin has been exposed. Intradental terminal endings extend into the inner dentin providing an anatomical basis for their observed dentin-response profile. Importantly, we show that intradental neurons collectively respond to superficial damage to the tooth indepedent of *Piezo2* or *Scn10a*. We find that chemogenetic activation of intradental HTMRs elicits orofacial grimace and hunched posture that is indicative of pain. Additionally, we demonstrate optogenetic activation of intradental HTMRs evokes a rapid jaw opening response via digastric muscle contractions that withdraws opposing teeth from contact. This work not only provides a cellular target for addressing acute tooth pain, but also provides a new perspective on intradental neuron function as sentinels that monitor for mechanical threats and initiate a reflex to protect the teeth.

## RESULTS

### In vivo identification of large diameter intradental neurons through electrical stimulation of the intact tooth

To enable functional evaluation of intradental neurons, we adapted an in vivo trigeminal Ca^2+^ imaging technique in mice to visualize real-time activity of sensory neurons^7,20^ while stimulating individual molars (Figure 1A-F, S1A-C). Unbiased labeling of trigeminal sensory neurons was achieved using neonatal AAV-Cre injections in Ai95D (Rosa-LSL-GCaMP6f) mice as previously described^12,21^. We confirmed this approach broadly labeled TG neurons across all cell diameters (range = 7-52 µm, mean = 28 µm, Figure S1E) with a ∼50% transduction efficiency (Figure S1F). Past work in large mammals indicates that electrical stimulation applied to exposed dentin can elicit single unit responses within the alveolar nerve^22^. Furthermore, the electric pulp test is routinely used in clinical dentistry to assess tooth sensation in human patients. Thus, we reasoned that low voltage pulses delivered to the occlusal surface of individual intact molar teeth may identify associated intradental neurons. Electrical stimulation of molar 1 or molar 2 (2-4 V, 0.6 Hz, 200 ms, Figure 1A, see Methods) resulted in stereotyped phasic, transient Ca^2+^ responses from separate groups of trigeminal sensory neurons (Figure 1B-D) indicating discrete sensory inputs from each tooth. In each experimental iteration, we identified pulsing neurons (12-22 neurons, n=10, Figure 1E) with large diameters (mean = 39.4 ± 0.4 µm, n=162 neurons, Figure 1F) consistent with the upper range of somas of Aδ myelinated sensory neurons. These neurons also responded to another clinical assay used to test human tooth pulp sensitivity^23^ (Endo-Ice, Figure S1D). To confirm that electrical stimulation preferentially activates intradental neurons, we performed retrograde labeling (CTB-AF647) to ipsilateral mandibular molars 1 and 2 prior to Ca^2+^ imaging^24^. Indeed, the majority of responding neurons were labeled by CTB (46/51, n = 4 mice, Figure SG-I) indicating that electrical stimulation is a reliable method for identifying intradental neurons. Taken together, we have established an in vivo method to identify intradental neurons within an intact tooth that enables subsequent molecular and functional evaluation.

**Figure 1.**
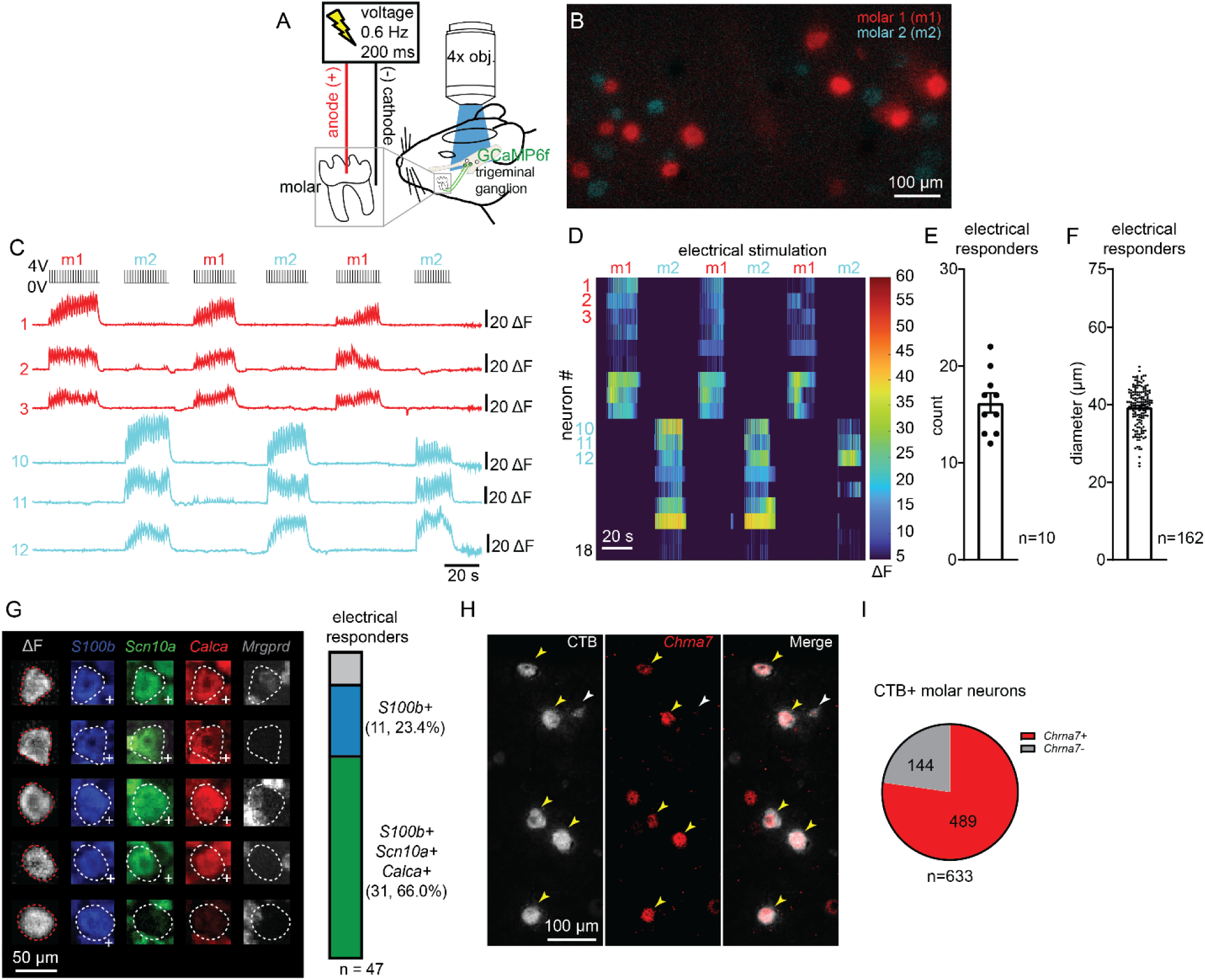
Functional imaging of intradental neurons reveals a large-diameter population of putative myelinated HTMRs. **(A)** Schematic of methodology used to record in vivo sensory neurons in the trigeminal ganglion (TG) in response to stimulation of mouse molar tooth in the oral cavity. Intradental neurons were identified via delivery of low voltage pulses (4V, 0.6 Hz, 200 ms) to individual mandibular molars. Data in (A-F) were obtained from adult Ai95(RCL-GCaMP6f)-D (Ai95D) mice injected postnatally (P0-P3) with AAV9-Cre. **(B-D)** Voltage stimulation enabled identification of intradental neurons innervating a single molar. (B) Imaging field showing merged single frames of Ca^2+^ response (ΔF). Neurons with responses entrained to voltage pulses to either molar 1 (m1, red) or molar 2 (m2, cyan). Scale bar, 100µm. (C) Example raw traces of GCaMP6f signal from identified m1 (red) and m2 (cyan) intradental neurons. Numbers correspond to neurons circled in (B). Bars to the right of traces indicate 20 ΔF. (D) Example heatmap from single TG showing in vivo GCaMP6f responses of 18 intradental neurons to voltage pulses. Neurons were grouped according to molar target of stimulation (m1 or m2). Stimuli used are indicated above the heatmap. Traces in (C) are designated neurons shown in (D). **(E)** Bar graph showing the mean number of neurons per trial that responded to ≥2 voltage pulses, designating them as intradental. Plotted individual data points represent the number of responding neurons in separate imaging trials. Error bars indicate the SEM. n = 10 mice. **(F)** Bar graph showing the mean diameter of neurons that responded to ≥2 voltage pulses, designating them as intradental. Plotted individual data points represent the number of responding neurons in separate imaging trials. Bar shows the mean value, and error bars indicate the SEM. n = 162 neurons from 10 mice. **(G)** Alignment of in vivo functional imaging responses with multiplexed whole-mount in situ hybridization (ISH) enables molecular classification of intradental neurons. Data for all experiments were obtained from Ai95(RCL-GCaMP6f)-D (Ai95D) mice injected postnatally (P0-P3) with AAV9-hSyn1-mCherry-2A-iCre, enabling mCherry expression to serve as guideposts to align functional imaging with ISH images. (Left) Multiplexed whole-mount ISH staining following alignment to in vivo GCaMP6f fluorescence. Probes: s100 calcium-binding protein B (*S100b*, blue), alpha-Nav1.8 (*Scn10a*, green), calcitonin gene-related peptide (CGRP; *Calca*, red), mas-related G protein-coupled receptor member D (*Mrgprd*, white). Plus sign in bottom right of each subpanel indicates positive staining. (Right) Vertical bar (parts of a whole) indicates number and percentage of neurons expressing selected genes in intradental neurons. *S100b, Scn10a,* and *Calca* mark the majority of functionally-responding intradental neurons designating these as large diameter, myelinated nociceptors indicative of HTMRs (66.0%, 31/47 responding neurons). Most other neurons were *S100b*-positive labeled cells indicating large-diameter myelinated cells (23.4%, 11/47 responding neurons). n = 5 mice, comprising 47 intradental neurons. (**H-I**) Representative image from retrograde-labeling of intradental neurons using CTB-AF647 tracer. Retrograde tracer CTB-AF647 was applied to small cavitations in maxillary and mandibular molars (m1/m2) 16 hr prior to tissue harvest. The expression of the candidate marker gene *Chrna7* was examined by ISH. (H) Left panel: CTB labeling; Middle panel: *Chrna7* expression; Right panel: Merge. Scale bar: 100 µm. Yellow arrow heads indicate retrograde-labeled intradental neurons that were co-positive for *Chrna7*. White arrowhead indicates cells that are only positive for CTB. (I) Pie chart comparing the expression of *Chrna7* of CTB-labeled intradental neurons. n=4 mice, comprising 633 retrograde-labeled intradental neurons.

### Intradental neurons express markers of putative HTMRs

We next sought to determine the molecular identity of responding intradental neurons. To this end, we performed Ca^2+^ imaging followed by post-hoc in situ hybridization of whole mount dissected trigeminal ganglion^12^. We reasoned that an initial panel of probes (*S100b*, *Scn10a*, *Calca*, and *Mrgprd*)^12,24^ would allow us to differentiate whether intradental neurons represent one or more broad mechanoreceptor subtypes. We again relied on neonatal AAV injection to induce simultaneous GCaMP6f and mCherry expression in a range of trigeminal sensory neurons (Figure S1E-F). mCherry fluorescence and ISH provided reliable guideposts (Figure S2A-C) to align in situ hybridization staining (Figure S2D-H) with fluorescence responses from Ca^2+^ imaging. These alignments revealed that some responding intradental neurons expressed *S100b* alone (11/47) but most expressed *S100b*, *Scn10a*, *and Calca* (31/47) which was consistent with our prior in vitro characterization indicating the majority represent nociceptive large diameter, myelinated HTMRs^12^. Conversely, intradental neurons lacked expression of *Mrgprd* (1/47), a marker of unmyelinated C-type HTMRs^12^(Figure 1G). To further confirm our molecular characterization, Ca^2+^ imaging of *S100b*+ trigeminal sensory neurons (*S100b*-Cre; Ai95D) enabled identification of intradental neurons (Figure S3A-C) and indicated that our electrical stimulation paradigm is capable of capturing >75% of all intradental neurons captured by retrograde labeling with CTB (range = 70%-87%, 83/111, n = 4, Figure S3D-F). Together these results demonstrate that intradental neurons primarily represent myelinated nociceptive HTMRs.

Recent studies revealed that HTMRs in the dorsal root ganglia (DRG) can be further differentiated by the expression of select transcriptional markers^25^, which may apply to myelinated HTMRs of the TG^26^. Specifically, *S100b*+/*Scn10a*+ myelinated HTMRs represent two groups (Figure S4A, B) that are distinguished by the expression of *Bmpr1b, Chrna7,* or *Smr2* (Figure S4C-E). To evaluate if intradental neurons represent one (or more) of these molecularly-defined subclasses of HTMRs, we first marked intradental neurons using retrograde tracing (CTB-AF647) from the ipsilateral molar teeth, then performed ISH to determine co-expression of *Bmpr1b, Chrna7,* or *Smr2*. As predicted, we found that *Bmpr1b, Chrna7,* and *Smr2* are expressed within subsets of TG sensory neurons (Figures 1H and S4F, H). Importantly, we determined that *Chrna7* (489/633, ∼77%, n = 4 mice, Figure 1I) was found in the majority of CTB+ intradental neurons. Few CTB+ neurons expressed either *Smr2* or *Bmpr1b* (*Smr2*: 5/229, ∼2%, n = 3 mice; *Bmpr1b*: 8/102, ∼8%, n = 3 mice; Figure SG, I). Taken together these data indicate that intradental neurons are a subset of myelinated HTMRs marked by *S100b*, *Scn10a*, and *Chrna7* expression.

### Intradental neurons respond to direct force after enamel removal

As intradental neurons express transcriptional markers of HTMRs, we next set out to determine their responses to mechanical forces directed to the intact tooth. We first tested whether intradental neurons respond to a range of innocuous to noxious forces applied to the enamel surface (0.4 - 20 g). Interestingly, we found that none of the intradental neurons were activated by direct mechanical forces despite being viable based on responsiveness to EndoIce (Figure 2A-C). As exposed dentin is associated with sensitivity in humans in the clinical setting^27^, we reasoned that intradental neurons might detect these mechanical forces in exposed teeth. To test this, we evaluated responses to application of force before and after exposing the dental pulp. As expected, we found that intradental neurons were not activated by the mechanical force on the intact tooth (Figure 2D). In contrast, in the same tooth immediately after the dental pulp was exposed, a portion of intradental neurons now responded to mechanical stimulation (9/54, 17% of electrical responders, Figure 2D, F). Furthermore, in teeth where only the dentin was exposed, direct mechanical force now recruited responses from the majority of intradental neurons (32/41, 78% of electrical responders, Figure 2E, F). Notably, responses were not due to immune cell infiltration at this immediate stage as show by no significant change in pan-immune (CD45+) staining of the pulp after dentin or pulp exposure compared to controls (Figure SA-M). From these experiments, we conclude that intradental neurons are HTMRs (intradental HTMRs) that do not contribute to any direct forces applied to the intact tooth but respond to mechanical force after the removal of enamel.

**Figure 2.**
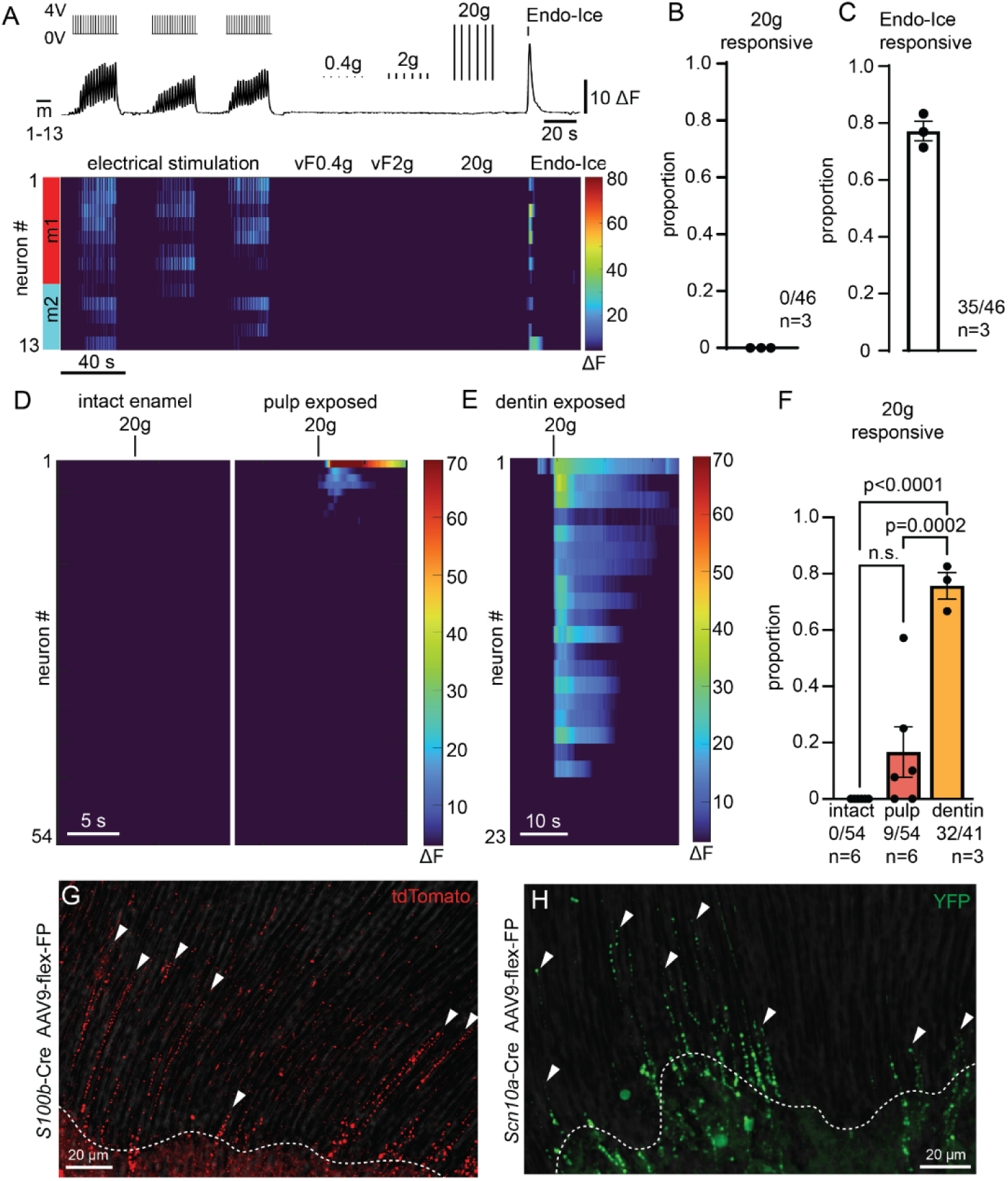
Intradental neurons are HTMRs that encode direct force after enamel removal and innervate the dental pulp and inner dentin. **(A-C)** Intradental neurons do not respond to direct forces applied to the intact tooth. (A) Example trace (top) and associated heatmap (bottom) showing absence of intradental neuron response to direct forces applied to the intact tooth, although they maintained responsiveness to Endo-Ice. Stimuli used are indicated above the traces. Bar to the right of trace indicates 10 ΔF. (B) Bar graph showing proportion of intradental neurons that respond to direct force. Plotted individual data points represent the proportion of force responsive/intradental neurons per trial. Bar shows the mean and error bars indicate the SEM. n = 3 mice, comprising 46 cells. (C) Bar graph showing proportion of intradental neurons that respond to Endo-Ice. Plotted individual data points represent the proportion of Endo-Ice responsive/intradental neurons per trial. Bar shows the mean and error bars indicate the SEM. n = 3 mice, comprising 46 cells. Data for A-C were obtained from Ai95(RCL-GCaMP6f)-D (Ai95D) mice injected postnatally (P0) with AAV9-Cre. **(D-F)** Intradental neurons respond to direct forces following structural damage of the tooth. (D) Heatmap showing response profiles of 54 intradental neurons from 3 mice when direct force is applied to the tooth while intact (left) and following pulp exposure via removal of overlying enamel and dentin (right). (E) Example heatmap showing the response of intradental neurons to direct force applied to exposed dentin. (F) Bar graph showing proportion of direct force response/intradental neurons before and after two forms of tooth structural damage. Plotted individual data points represent the proportion of force responsive/intradental neurons per trial. Bar shows the mean and error bars indicate the SEM. n = 6 mice for intact enamel, n = 6 mice for pulp exposed, n = 3 mice for dentin exposed. p-values from one-way ANOVA with Tukey correction indicated with the graph. Data for D-F were obtained from Ai95(RCL-GCaMP6f)-D (Ai95D) mice injected postnatally (P0-P3) with AAV9-Cre. **(G)** Representative image of immunostained neuronal terminals merged with brightfield image of dentin tubules in demineralized molars from *S100b*-Cre mice receiving TG injection of AAV9-flex-tdTomato to induce sparse labeling of trigeminal sensory neurons. Enamel, a fully mineralized structure, has been removed completely by demineralization. This experiment was repeated in n = 5 mice. Scale bar: 20 µm. **(H)** Representative image of immunostained neuronal terminals merged with brightfield image of dentin tubules in demineralized molars from *Scn10a*-Cre mice injected using a Cre-dependent AAV to induce sparce labeling of terminals (see methods). Enamel, a fully mineralized structure, has been removed completely by demineralization. This experiment was repeated in n = 4 mice with additional littermate and no primary controls. Scale bar: 20 µm.

### Intradental HTMRs terminate in the tooth pulp and inner dentin

Our experiments testing direct force on tooth structures suggest the presence of terminal endings of the intradental neurons within the dentin, as mechanical stimulation of exposed dentin recruits their activation in the TG. We applied genetic strategies relying on *S100b* and *Scn10a* to drive fluorescent labeling of intradental HTMRs to visualize their terminal endings. While *S100b* delineates myelinated somatosensory neurons, it is also broadly expressed in the nervous system as well as non-neuronal peripheral cells. To restrict fluorescent labeling to *S100b*+ somatosensory neurons we crossed *S100b*-Cre into a Cre-dependent neuronal specific reporter strain (*Snap*25-LSL-2A-eGFP). We found dense GFP+ innervation of the coronal pulp as well as parallel GFP+ terminals penetrating to the inner 3^rd^ of the dentin (Figure S6A-I). As *Scn10a* expression is highly restricted to sensory neurons, we repeated immunohistochemical experiments in teeth obtained from *Scn10a*-Cre; Ai32 mice (*Scn10a*-Cre; LSL-ChR2-YFP). Here, we observed remarkably similar YFP+ innervation patterns within the pulp and inner dentin (Figure S6J-O).

While these studies indicate that dentin is uniformly innervated by endings stemming from intradental neurons, the endings of individual neurons could not yet be discerned. We reasoned that sparse labeling of terminal endings^28^ (Figure S7A-D) would reveal terminal patterns within the dentin. Here, fluorescently-labeled fibers (*S100b*-tdT and *Scn10a*-GFP) were localized to only a portion of the overall dentin typically within a single pulp horn (Figure 2G, H) and exhibited tuft-like patterning penetrating into non-contiguous dentinal tubules. These findings indicate that single neurons target several tubules and multiple neurons innervate a local region of the inner dentin. With our sparse labeling, we also noted that terminal morphology appeared as a series of puncta running in parallel along the length of dentin tubules (Figure 2G, H). Taken together, we establish that intradental HTMRs, as defined by expression of *S100b* and *Scn10a*, contribute intermingled terminals to create dense innervation of the inner dentin.

### Intradental HTMRs function to detect tooth enamel damage

Intradental HTMRs respond to mechanical stimulation following dentin exposure, providing input *after* the inner layers of the tooth have been exposed. We next sought to explore if these neurons also activate *as* molar teeth are damaged. To this end, we monitored responses of trigeminal sensory neurons while simultaneously damaging the enamel surface using a dental bur. Indeed, cutting of the surface enamel evoked responses from most intradental HTMRs (64/70, 91%, n = 4, Figure S8A-C). Conversely, vibration (50 - 200 Hz) applied to the tooth evoked almost no activity (2/52, 3%, Figure S8D-G) indicating these cells respond to the damage component of cutting. Taken together, these experiments indicate that intradental HTMRs encode damaging mechanical stimulation of individual teeth.

### Intradental HTMRs detect superficial frictional damage of the tooth surface and encode damage severity

The cutting of the exterior tooth produced overt damage of the enamel layer (Figure S9A) and triggered a response from intradental HTMRs. Does the response profile of these neurons provide information about the degree of tooth damage? To this end we sought to introduce a stimulus that creates lesser damage by removing the carbide flutes from the dental bur. This “fluteless” bur produces friction, but not perceptible damage (Figure S9A). Remarkably, friction on the surface of the tooth triggered the activation of almost half of the intradental HTMRs identified by electrical stimulation (98/205, 47%, n = 7 mice, Figure 3A-C). Subsequently, the cutting stimulus evoked responses in these same initial 98 neurons in addition to recruiting responses from 58 additional intradental HTMRs (156/205, 76%, n = 7 mice, Figure 3A-C). Interestingly when examining neurons that responded to both stimuli, friction triggered smaller magnitude responses compared to cutting, suggesting that intensity of activation increases with degree of tooth damage (Figure 3D). Post hoc in situ again confirmed that intradental HTMRs that respond to tooth cutting largely represent *Scn10a*+ neurons (22/27, ∼82%, n = 4 mice, Figure 3E). We found that cutting and friction produced equivalent, minimal warming of the inner tooth (<3⁰C, Figure S9B-D) that is unlikely to trigger any response and cannot account for our observed differences between response magnitude. These results demonstrate that intradental HTMRs detect and encode the degree of superficial frictional damage.

**Figure 3.**
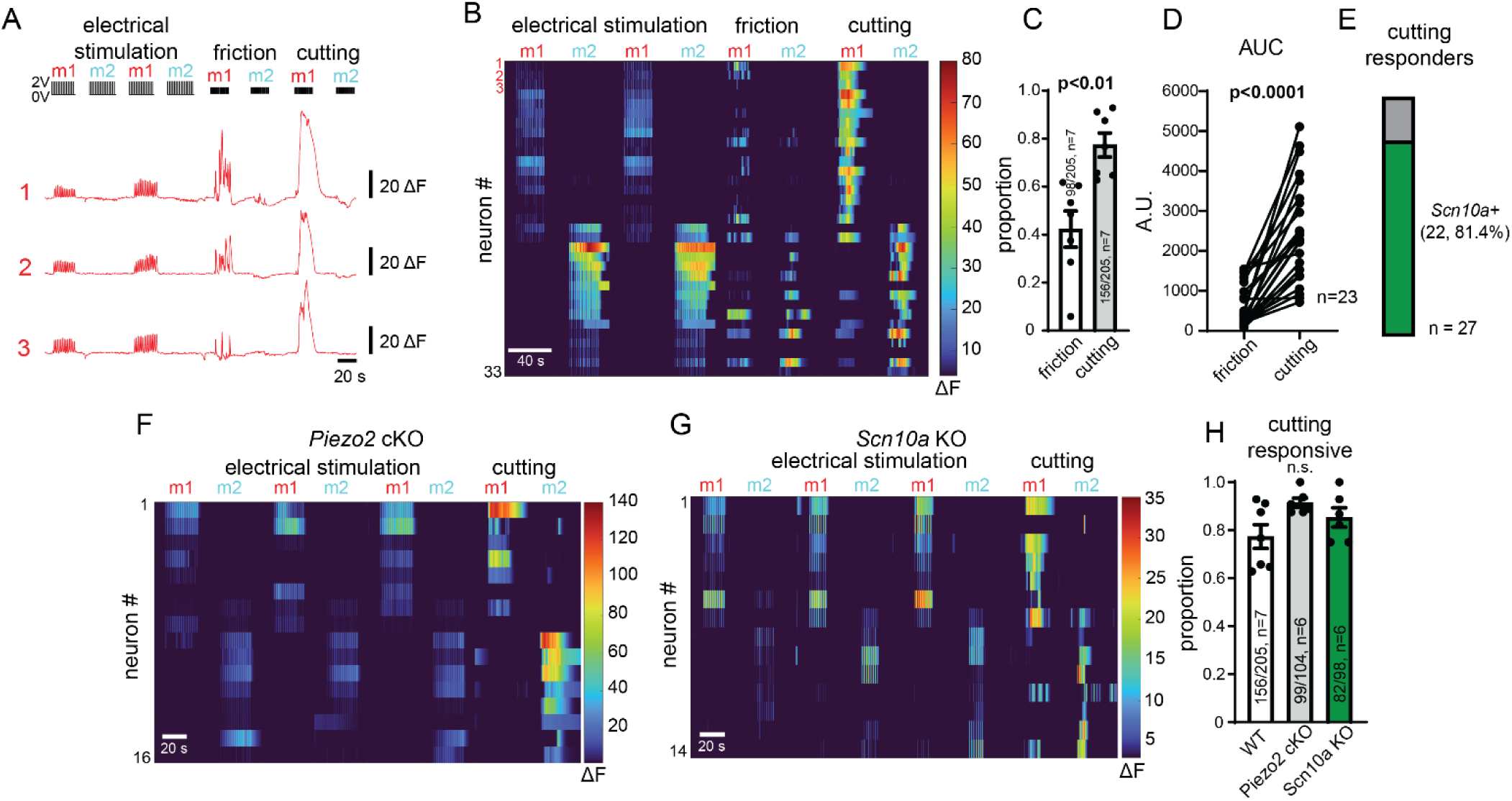
Intradental HTMRs detect enamel damage, independent of *Piezo2* or *Scn10a* (Nav1.8). **(A-E)** Intradental HTMRs sense cutting and superficial frictional damage of tooth enamel. (A) Example traces showing intradental neuron response to enamel cutting and friction applied to individual molars. Stimuli used are indicated above the traces. Bars to the right of traces indicate 20 ΔF. (B) Example heatmap containing traces shown in (A). (C) Bar graph showing proportion of friction and cutting responsive intradental neurons identified by electrical stimulation. Plotted individual data points represent the proportion for each trial. Bar shows the mean and error bars indicate the SEM. n = 7 mice, comprising 205 total cells, p<0.01, Unpaired t-test. (D) Line graph depicting magnitude of Ca^2+^ responses of intradental neurons to friction versus cutting. Individual points represent calculated area under the curve (AUC) during designated stimulation for each cell. n = 23 cells, p<0.0001, Paired t-test. (E) Vertical bar (parts of a whole) indicates number and percentage of cutting responsive neurons that express *Scn10a* (81.4%, 22/27 cutting responsive neurons, n = 4 mice). Results in E are cutting responses from primary data also used in Figure 1G. **(F)** Example heatmap showing stereotypical intradental HTMR responses to cutting persist in *Piezo2*-cKO mice. **(G)** Example heatmap showing stereotypical intradental HTMR responses to cutting persist in *Scn10a*-KO mice. **(H)** Bar graph depicting the proportion of intradental HTMRs that respond to enamel cutting in wildtype, *Piezo2*-cKO, and *Scn10a*-KO mice. Plotted individual data points represent the calculated proportion of enamel cutting responsive/intradental neurons. Bar shows the mean, and error bars indicate the SEM. Intradental neurons were imaged from n = 6 or 7 adult mice per genotype. No significant differences were observed with p>0.05 using One-way ANOVA with Tukey’s correction.

### Damage responses of intradental HTMRs are independent of *Piezo2* or *Scn10a* (Na_v_1.8)

Our previous work indicated that intradental neurons express the mechanosensitive ion channel *Piezo2*^24^ that underlies mechanosensation in the skin^12,21^. Is Piezo2 required for intradental HTMR activation by enamel cutting? To investigate this, we examined neuron responses after AAV-mediated conditional knock out of *Piezo2* (Piezo2-cKO) which circumvents lethality of global *Piezo2* deletion^12,21^. We first validated loss of *Piezo2* in sensory neurons using in situ hybridization against *Piezo2* and in vivo calcium imaging. Indeed, we found that neurons that were transduced with AAV-Cre (i.e., GCaMP6f+ neurons) had marked loss of cytoplasmic *Piezo2* using ISH (Figure S10A, B). Furthermore, as previously demonstrated^12,17^, we found that *Piezo2*-dependent innocuous cheek brush largely failed to evoke responses from GCaMP6f+ neurons in calcium imaging in Piezo2-cKO mice (Figure S10C). Next, we examined responses of intradental neurons to cutting of the molar tooth in the context of Piezo2-cKO. Despite loss of *Piezo2*, we observed no significant change in the proportion of intradental HTMRs responding to cutting (Figure 3F, H). These findings indicate that neuronal detection of enamel damage persists despite the absence of *Piezo2*.

Intradental HTMRs are marked by *Scn10a* expression, encoding the pore-forming alpha subunit of Na_v_1.8 that contributes to noxious mechanical pain^4^. We next sought to determine whether *Scn10a* is required for intradental HTMR sensory transduction in response to enamel damage. To this end, we examined neuronal responses in the well-established knock in/knock out Scn10a-Cre driver strain^29^ after validating genotypes by PCR. As expected, *Scn10a*-Cre enabled capture of intradental HTMRs with similar proportion of cutting responders (Figure 3C, S8A-C, S10D-F). However, in homozygotes where *Scn10a* translation is lost from Cre expression^30^, we found no reduction in the proportion of intradental HTMRs responding to enamel cutting (Figure 3G, H). These experiments indicate that the *Scn10a* gene product (Na_v_1.8) is dispensible for the transduction of damage responses. Taken together, these experiments suggest an unknown high-threshold mechanical receptor may underlie damage responses of intradental HTMRs.

### Chemogenetic activation of intradental HTMRs elicits a marked pain phenotype

Activation of sensory neurons within the tooth via exposed dentin is known to elicit pain in humans^31^. Considering our transcriptional analysis and in vivo imaging support that intradental neurons represent nociceptive HTMRs, we next sought to confirm that their activation produces pain in freely-moving animals. To this end, we crossed the *Scn10a*-Cre driver into the CAG-LSL-Gq-DREADD (hM3Dq) to enable selective activation of *Scn10a*+ sensory neurons using clozapine-N-oxide (CNO). We reasoned that this chemogenetic strategy (*Scn10a*-DREADDq) would permit both broad activation of *Scn10a*+ sensory neurons following i.p. administration of CNO as well as targeted activation^32^ of intradental neurons after application of CNO directly to the tooth (Figure 4A). First, we captured behaviors of *Scn10a*-DREADDq and controls lacking DREADDq at baseline and following administration of CNO (0.1 mg/kg, i.p.). *Scn10a*+ sensory neurons primarily represent nociceptors^11^ and, as expected, their widespread activation in *Scn10a*-DREADDq mice elicited marked postural and orofacial expressions consistent with hunching and grimace that are associated with pain^33^. These elements were largely absent in baseline or controls even when animals were not moving (i.e., resting). Conversely, behaviors such as locomotion, rearing, and grooming were common in baseline and control videos. Examples of pain, resting, locomotion, rearing, and grooming were extracted from these initial recordings (Figure 4B) and used to train the AI-driven behavioral analysis tool LabGym^34,35^ to enable automated scoring of these behaviors in full-length videos. Quantification confirmed that after 0.1 mg/kg CNO i.p., *Scn10a*-DREADDq mice predominately exhibited pain (Figure 4C, D, G), while control mice demonstrated resting, locomotion, and grooming that was comparable to uninjected mice at baseline (Figure S11A-E). We next sought to evaluate pain following application of CNO to the tooth. While we expect that CNO should remain localized to the site of application, we reduced the dosage of CNO 10-fold to further reduce possible systemic effects of CNO. Remarkably, after 0.01 mg/kg CNO was applied to the tooth (see Methods for details), we found that *Scn10a*-DREADDq mice, but not control mice, exhibited a duration of pain comparable to those receiving 0.1 mg/kg CNO i.p. (Figure 4E, G, see Table 1 for statistics). *Scn10a*-DREADDq mice receiving 0.01 mg/kg CNO i.p. exhibited minimal pain (Figure 4F-G, see Table 1 for statistics) indicating that pain from CNO applied to the tooth was not due to off-target effects. Furthermore, mice in control groups demonstrated significant increases in locomotion and resting and trended toward more grooming compared to *Scn10a*-DREADDq mice receiving tooth CNO (Figure S11A-E, see Table 2-6 for statistics). Together these data support that intradental HTMRs are nociceptors that produce pain upon activation.

**Figure 4.**
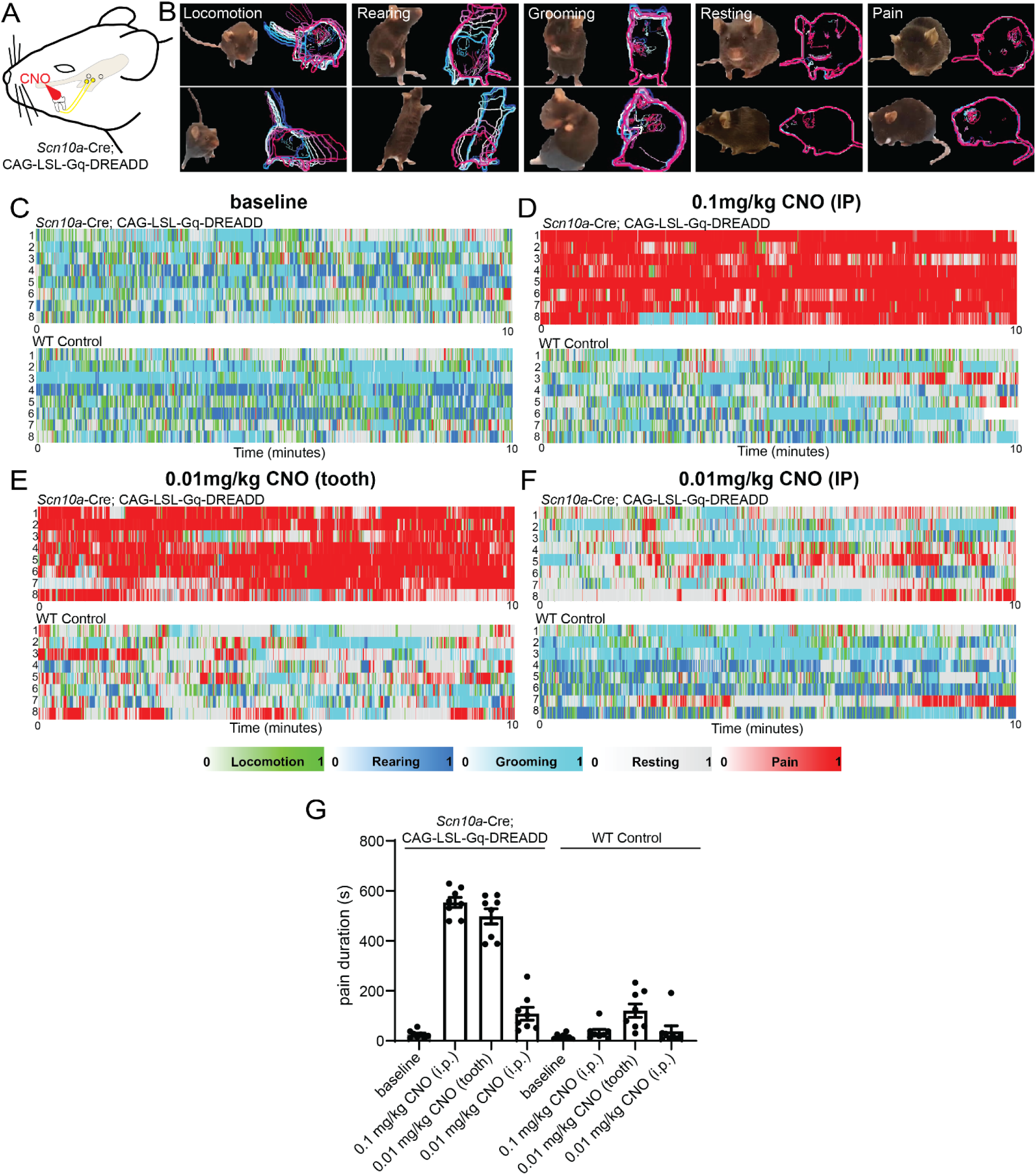
Chemogenetic activation of intradental HTMRs elicits a marked pain phenotype. (**A**) Schematic demonstrating the experimental approach. The excitatory Gq receptor hM3Dq (CAG-LSL-Gq-DREADD) was driven by *Scn10a*-Cre. To selectively activate intradental neurons, CNO was directly applied to the tooth (see Methods). (**B**) Representative examples showing a single video frame with the background removed (left) and corresponding motion pattern images generated by LabGym (right), illustrating different behavioral categories detected by the software. An example from the front (top) or side (bottom) is provided for each behavior. Each behavior was defined within a 10-frame time window extracted from videos with an extracted frame rate at 15 fps. During time window, the video clip and the associated pattern image form a data pair used by LabGym for training and analysis. In the pattern images, color curves represent the contours of the whole body and key body parts at successive time points, capturing the dynamics of the behavior. (**C-F**) Raster plots showing the behavior categorizations over time for *Scn10a-*Cre; CAG-LSL-Gq-Dreadd (top) and wildtype controls (bottom) during (C) baseline, (D) 0.1mg/kg CNO i.p., (E) 0.01 mg/kg CNO applied to unilateral m1 and m2 mandibular molars, and (F) 0.01mg/kg CNO i.p. Color bars indicate behavioral categories, with color intensity reflecting the probability of each behavior. The x-axis denotes time (seconds) and the y-axis denotes individual animals, with each row corresponding to a single animal. n = 8 animals per genotype. (**G**) Bar graph depicting pain duration corresponding to panels C-F. Plotted individual data points represent the cumulative pain duration for individual animals. Bar shows the mean, and error bars indicate the SEM (see Table 1 for statistics).

### Optogenetic activation of intradental HTMRs induces digastric muscle contraction and initiates a protective jaw opening reflex

Whereas acute pain has clear utility in certain forms of injury, tooth pain has little benefit toward mammalian fitness. Indeed, the stereotyped pain behavior evoked by chemogenetic activation (Figure 4B, E, G) may paradoxically reduce the ability of the animals to respond to environmental danger. In addition to producing pain, somatosensory HTMRs are also known to initiate reflexes to protect structures from damage^7^. Considering teeth are at risk of damage from masticatory forces and/or hard food substrates during regular function, we next sought to determine whether activation of the intradental HTMRs might drive protective responses that contribute to fitness. Interestingly, low intensity electrical stimulation of teeth in larger mammals has been shown to elicit electromyogram (EMG) activity in some muscles of mastication^36^. Indeed, we first confirmed that electrical stimulation of the inferior alveolar nerve (IAN) elicited EMG activity in the anterior digastric muscle (Figure S12A) which contributes to jaw opening (i.e., mandibular deflection). We next sought to determine if selective optogenetic activation of intradental neurons was capable of inducing EMG activity in the anterior digastric muscle. To this end, we relied on AAV-Cre transduction of intradental neurons via cavitations in the molar teeth to initiate the expression of channelrhodopsin-2 (ChR2) in a Cre-dependent effector strain (Ai32, Figure 5A, S12B). ISH confirmed that AAV transduction targeted intradental neurons since ChR2-YFP+ sensory neurons largely expressed both *S100b* and *Scn10a*+ (103/123 neurons, n = 4 mice, Figure S12C, D). Activation of intradental neurons expressing ChR2+ using blue light stimulation (15 ms pulse) of the trigeminal ganglion produced an immediate, transient waveform in the ipsilateral anterior digastric EMG (Figure 5B-C, S6D, F-H; 0.83 +/- 0.13 mV amplitude, n = 5 mice). Alternatively, we found no EMG activity in response to blue light stimulation when intradental neurons were negative for ChR2. The ipsilateral masseter muscle that contributes to closing the jaw also failed to demonstrate EMG activity after light stimulation in either condition (Figure 5B, S12E). Prolonged blue light stimulation (100 ms pulse) elicited similar waveforms in ipsilateral digastric EMG (Figure S12E-H; p>0.05) likely reflecting equivalent ChR2 inactivation. Changes in EMG activity were never observed at light offset (Figure 5B, S12 D, E). Taken together, these data indicate that somatic optogenetic activation of intradental neurons leads to electrical activity of the anterior digastric muscle. The anterior digastric muscles function to open the jaw by depressing the mandible^37^.

**Figure 5.**
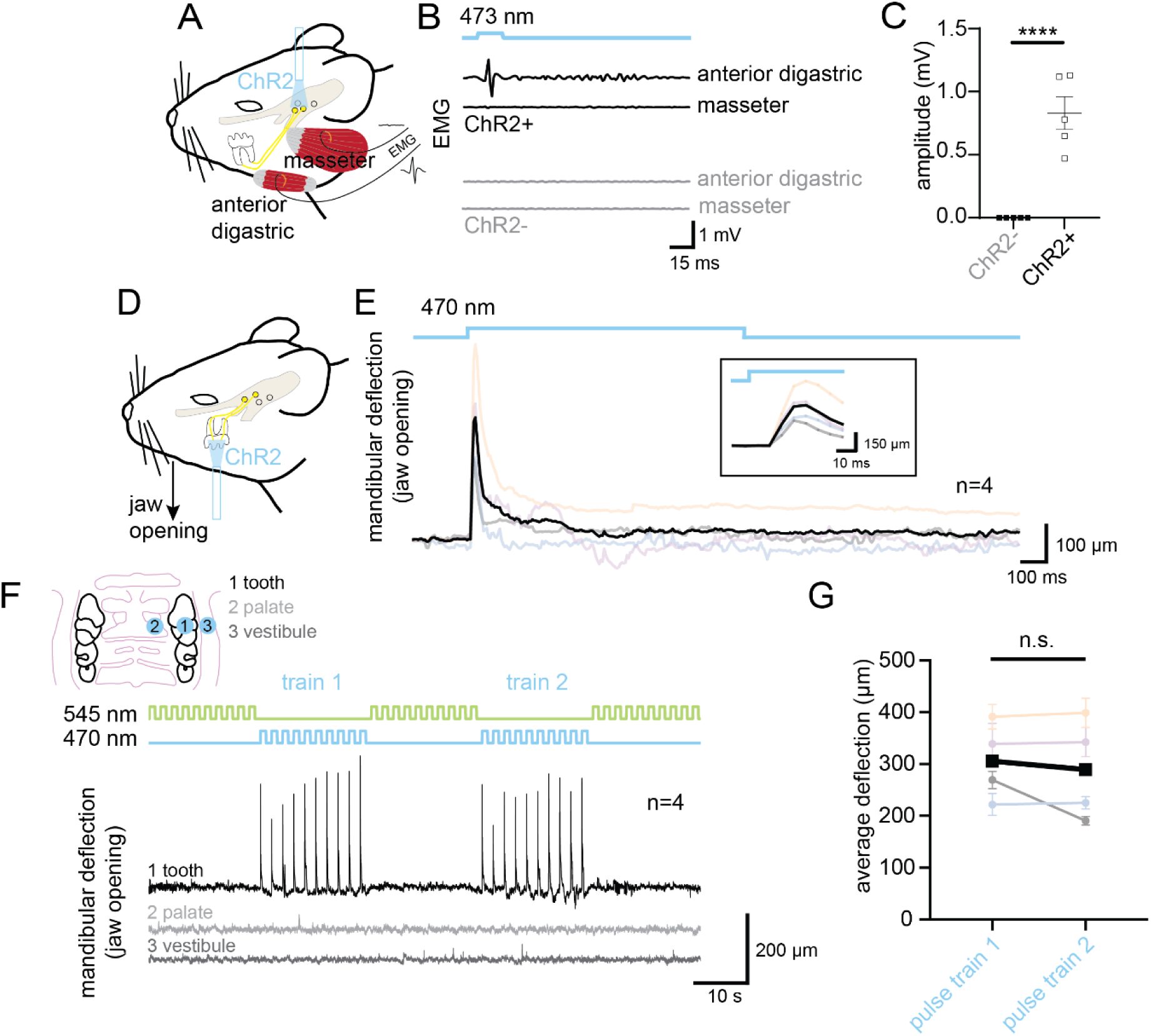
Optogenetic activation of intradental HTMRs induces digastric muscle activity and initiates a jaw opening reflex. **(A-C)** Optogenetic activation of intradental neurons induces activity in the anterior digastric muscle EMG. (A) Schematic demonstrating the experimental approach to selectively activate intradental neurons while recording EMG from the anterior digastric and masseter muscles. Channelrhodopsin-2 was selectively expressed in right mandibular intradental neurons innervating M1 and M2 molars via tooth injection of AAV6/2-hEF1a-iCre into Ai32(RCL-ChR2(H134R)/EYFP) mice. An optical fiber was implanted over the right trigeminal ganglia to selectively activate intradental neurons while EMG signals of the ipsilateral anterior digastric or masseter muscles were recorded. (B) Example traces demonstrating EMG activity elicited by optogenetic activation of intradental neurons (15 ms, 473 nm). EMG activity is observed in the ipsilateral anterior digastric muscle, but not masseter muscle. Mice lacking channelrhodopsin-2 expression in intradental neurons show no changes in EMG activity for either the anterior digastric or the masseter muscles in response to blue light stimulation; n = 5 mice for all conditions. (C) Graph depicting the amplitude of activity of digastric muscle EMG in response to blue light stimulation (15 ms, 473 nm). Plotted individual data points represent the peak-to-peak amplitude of the elicited EMG trace. Error bar indicates SD. n = 5 mice per condition. Significant differences were observed between experimental (ChR2+) and control animals (ChR2-) with p<0.0001 using Unpaired t-Test. **(D-G)** Optogenetic activation of intradental neuron terminals in the intact tooth results in mandibular deflection (i.e. jaw opening). (D) Schematic demonstrating the experimental approach. Channelrhodopsin-2 expression (Ai32) was driven by *Scn10a-*Cre. An optical fiber (500 µm) was positioned perpendicular to the surface of the first maxillary molar tooth or adjacent oral structures (hard palate or oral vestibule). Deflection was determined by monitoring the position of maxillary and mandibular incisors (see Methods). (E) Traces of average measured mandibular deflection in response to a single blue-light pulses (470 nm, 1s) directed to the molar tooth. Black trace represents the average (n = 4 mice). Multicolored traces represent average measurements from each mouse (n = 10 traces/mouse). Inset shows brief timescale of trace from (E) demonstrating kinetics of mandibular deflection in relation to light onset. (F) Average trace of measured mandibular deflection in response to trains of blue or green light pulses (470nm or 545nm, 10 pulses, 0.5 Hz, 1s) directed at oral structures. Sequential blue light pulses directed at the molar tooth (black trace), but not adjacent structures (light and dark grey traces) induced mandibular deflection after each blue light onset. No jaw movements were observed in response to green light stimulation. (G) Line graph depicting average mandibular deflection in response to sequential blue light pulse train stimulation of molar. Average deflection from each mouse is plotted as multicolored individual points and represents mean peak amplitude and error bars show SEM of mandibular deflection during stimulation. n = 10 measurements per mouse per train related to traces shown in (F). Black points represent the average of n = 4 mice. p>0.05 using Unpaired t-Test.

To directly determine if collective activation of intradental HTMRs initiates overt jaw movement, we next employed a genetic strategy to drive ChR2 expression in these cells using *Scn10a*-Cre. For these experiments, we selectively activated intradental HTMRs by directing our optical fiber onto a single intact molar tooth (Figure 5D). Here, we found that pulses of blue light (470 nm, 1s) elicited a rapid, brief mandibular deflection (10 ms onset, reversing after 30 ms, Figure 5E). Paralleling EMG activity, light offset had no effect on jaw position. We determined that repeated blue light pulses (10 pulses, 0.5 Hz, 1s) induced a series of mandibular deflections entrained to light onset (Figure 5F) that did not diminish between pulse trains (Figure 5F-G, S12I), average deflection 305 +/- 37 vs 289 +/- 49 µm, p>0.05). Conversely, green light (545 nm) failed to produce any jaw movements despite identical stimulating frequency and duration (10 pulses, 0.5 Hz, 1s). Importantly, blue light stimulation of molar-adjacent tissues (i.e., hard palate or oral vestibule) also failed to produce observable jaw movements (Figure 5F). Notably, despite the short latency of jaw movement following blue light stimulation, we found an absence of direct *Scn10a*+ projections to the motor nucleus (5N) controlling the anterior digastric (5ADi) (*Scn10a*-Cre; LSL-ChR2-YFP, Figure S13A, B). Taken together, these experiments reveal that optogenetic activation of intradental HTMRs indirectly drives a rapid jaw opening reflex.

## DISCUSSION

Mammalian teeth are specialized internal organs that endure a range of mechanosensory stimuli during biting and mastication. While intradental neurons have been known to evoke pain through their activation, a biological role had not been defined for this interoceptive innervation using modern approaches. Here, we establish an in vivo functional imaging approach to identify intradental neurons in an intact tooth, define their responses to mechanical damage of the tooth, and determine that their targeted activation leads to pain and jaw opening. Our experiments reveal a population of intradental HTMRs that 1) detect forces applied to the exposed dentin as well as damage of the superficial enamel, 2) do not contribute to the detection of innocuous forces encountered by the tooth, 3) target the inner dentin with intermingled terminals providing anatomical evidence supporting their coordinated response to stimuli, 4) rely on a Piezo2- and Scn10a-independent mechanism to transduce damage responses, 5) drive pain in a free-moving animal and 6) lead to digastric muscle contraction and a jaw opening reflex.

Protective reflexes have been described for the knee (patellar reflex), cornea (eye-blink reflex) and the jaw (jaw-jerk reflex) that occur within 100 ms latencies. Here, we describe an extremely rapid reflex (5-15 ms) that is initiated following the activation of terminals of intradental HTMRs within the dentin and indirectly triggers digastric muscle contraction. We predict that the intradental HTMRs respond when the jaw is closing and one or more teeth make unexpected contact with hard foods (i.e., cartilage or bone) or opposing teeth. In this scenario, the activation of intradental HTMRs would result in the contraction of the digastric which would ostensibly slow, stop, or reverse the closing mandibular movement. Based on our data, activation of intradental HTMRs within a single tooth during mastication would be sufficient to drive a reflex response. Future work will be required to determine the brainstem circuit that underlies the jaw-opening reflex.

Recent work indicates that *Calca*+ circumferential-HTMRs (circ-HTMRs) innervating hair follicles in skin mediate a nocifensive response^7^. Intradental HTMRs are most similar to the CGRP-ζ based on their expression of *Chrna7*^26^. However, intradental neurons are likely to be distinct from CGRP-ζ neurons found within the DRG considering their lack *Smr2* and their unique terminal morphology^19^. Thus, we propose that intradental HTMRs represent a new subclass of HTMRs that innervates and protects other vulnerable tissue structures by engaging withdrawal reflexes.

Intradental HTMRs efficiently respond to damage of the tooth indicating they may be tuned for its detection; however, the precise component of the damage stimulus that triggers intradental HTMR responses is not known. The hydrodynamic theory posits that stimuli of varying etiology (e.g., thermal, mechanical, or chemical) converge on microfluidic fluid flow through dentinal tubules which mechanically-activates receptors and triggers sensation^38^. We cannot rule out that superficial damage of the tooth may contact a portion of dentin on the mouse molar to initiate fluid flow. Our observations that intradental HTMRs respond to cold and direct force following dentin exposure further support this possibility. While ‘HTMRs’ reflects harmonized somatosensory nomenclature^26^, designating them as ‘myelinated mechano-nociceptors’ would be more accurate if miniscule fluid movements and associated low forces underlie the activation of intradental neurons. Follow-up studies are necessary to provide important distinction. For instance, comparing micro-structural changes in the tooth caused by vibration vs. cutting could reveal the activating component and whether there is accompanying fluid flow. Alternatively, performing in vivo imaging in conjunction with precise thermal manipulation of the teeth may enable differentiation of molecular vs. physical gating and uncoupling the dependence of thermal responses on fluid flow.

Intradental neurons have been previously suggested to function as LTMRs^39^. On the contrary, our data directly show that intradental neurons fail to respond to direct forces or vibration applied to the intact tooth. Instead, it is highly probable that light forces applied to the teeth are transduced by LTMRs innervating surrounding periodontal (i.e., tooth surrounding) tissues. Future studies investigating the molecular identity and innervation targets within surrounding periodontium tissues will shed insight into the mechanisms for how direct mechanical forces applied to the tooth are detected.

We determined that damage responses by intradental neurons are not abolished by their loss of the canonical mechanoreceptor Piezo2 or the nociceptive transmitter Na_v_1.8. We cannot rule out that these membrane channels may contribute to tooth sensation; however, our data suggest that blocking either ion channel in intradental neurons is unlikely to prevent their activation while the teeth are structurally damaged. This is in contrast to a recent clinical trial that supports the effectiveness of inhibition of Na_v_1.8 to diminish post-surgical inflammatory pain^40^. While we acknowledge that these data represent negative findings, if substantiated, they hint at the existence of unidentified molecular sensors for mechanical stimulation of the tooth. Single-cell sequencing of intradental neurons^41^ toward screening and identification of novel mechanoreceptors and nociceptive transducers will provide additional targets for next-generation dental anesthetics that can be validated with our approach.

Our work provides fundamental insight into the basis and role of tooth sensation, but some open questions remain. While some evidence suggests that the tooth pulp receives innervation from C-fiber nociceptors that contribute to forms of tooth-related pain, our work to-date suggests these are unlikely to contribute to the protective reflex response. Our findings do not exclude the possibility that C-fibers may be present deep in the tooth pulp where they could contribute to differential aspects of tooth physiology in the context of inflammation. Furthermore, our study did not explore the contribution of resident pulp cells (i.e. odontoblasts) to dental sensation. While odontoblasts have been proposed as a putative sensory cell based on in vitro assays^42^, these cells lack both vesicles and microscopic proximity to nerve terminals that would enable traditional synaptic transmission. Indeed, neuronal HTMR counterparts in the epidermis skin terminate as free nerve endings^43^ and act as the primary sensory cell. Future experiments will be necessary to determine if activation of odontoblasts elicits responses from intradental neurons. Clinically, we predict intradental HTMRs may underlie acute pain during biting as part of cracked tooth syndrome^44^. Further work will be needed to determine if responses of intradental HTMRs are altered by the inflammatory milieu and can explain other forms of dental pain. For instance, intradental HTMRs may exhibit lower threshold responses to mechanical stimulation in the context of tooth infection, and begin to contribute to the exquisite pain associated with toothache.

Our study indicates that intradental HTMRs respond to damage, initiate an important protective reflex, and drive overt pain behaviors. Chemogenetic activation of intradental HTMRs represents a new model for orofacial pain that can be leveraged to evaluate analgesics for their ability to attenuate pain as well as reveal additional orofacial features that correlate with the experience of pain. Further experiments would examine if the jaw-opening reflex and pain from the activation of intradental neurons are necessarily concurrent or may be uncoupled in freely-behaving animals.

## ACKNOWLEDGEMENTS

This work is supported by K22DE029779 (J.J.E.), T32DE007057 (E.A.R.), and T32DC00011 (E.A.R.). We thank Dr. Nicholas Ryba for his feedback throughout this project and valuable comments on this manuscript. We thank Dr. Nima Ghitani for his technical assistance with in vivo imaging. We thank Dr. Alexander Chesler, Dr. Lauren Surface, and members of the Emrick lab for helpful comments and suggestions. We are grateful to donors of knockin mice for *S100b*-Cre knockin mice (Ryba); *Piezo2*-cKO knockin mice (Chesler); and *Snap25*-LSL-2A-EGFP-D knockin mice (Dr. Claire LePichon).

## AUTHOR CONTRIBUTIONS

All authors gave final approval and agreed to be accountable for all aspects of the work. J.J.E. and E.A.R. conceptualized and directed the study. E.A.R., A.R.G., S.W., and J.J.E. performed Ca^2+^ imaging experiments with acquisition help from M.N., and assistance in analysis from T.S.. E.A.R., M.E.G., I.B., A.R.G., B.S.C.C, A.J., K.J.B., and J.J.E. performed survival surgeries and ISH and histological studies. E.A.R. performed chemogenetic behavior experiments with assistance from Y.H., L.D. and S.W.. Y.H., E.A.R., and L.D. analyzed chemogenetic behavior data using LabGym. K.H.U.K. performed optogenetic stimulation and EMG measurements and analyzed the data. E.A.R., B.S.C.C., and J.J.E. performed optogenetic stimulation and mandibular deflection measurements. L.Y. evaluated brainstem projections of nociceptive neurons. E.A.R., A.R.G., S.W., Y.H., B.Y., K.P.P., P.L. and J.J.E. interpreted the data. J.J.E. and E.A.R. wrote the manuscript and generated figures with input from all other authors.

## REFERENCES

1. Henry, M. A. & Hargreaves, K. M. Peripheral Mechanisms of Odontogenic Pain. Dental Clinics of North America 51, 19–44 (2007).

2. Renton, T. Dental (Odontogenic) Pain. Rev Pain 5, 2–7 (2011).

3. Murthy, S. E. et al. The mechanosensitive ion channel Piezo2 mediates sensitivity to mechanical pain in mice. Sci Transl Med 10, eaat9897 (2018).

4. Akopian, A. N. et al. The tetrodotoxin-resistant sodium channel SNS has a specialized function in pain pathways. Nat Neurosci 2, 541–548 (1999).

5. Xu, K.-H. et al. Association of Tooth Loss and Diet Quality with Acceleration of Aging: Evidence from NHANES. Am J Med 136, 773–779.e4 (2023).

6. Koka, S. & Gupta, A. Association between missing tooth count and mortality: A systematic review. J Prosthodont Res 62, 134–151 (2018).

7. Ghitani, N. et al. Specialized Mechanosensory Nociceptors Mediating Rapid Responses to Hair Pull. Neuron 95, 944–954.e4 (2017).

8. Barik, A., Thompson, J. H., Seltzer, M., Ghitani, N. & Chesler, A. T. A Brainstem-Spinal Circuit Controlling Nocifensive Behavior. Neuron 100, 1491–1503.e3 (2018).

9. Slade, G. D. Epidemiology of dental pain and dental caries among children and adolescents. Community Dent Health 18, 219–227 (2001).

10. Byers, M. R. & Cornel, L. M. Multiple complex somatosensory systems in mature rat molars defined by immunohistochemistry. Archives of Oral Biology 85, 84–97 (2018).

11. Nguyen, M. Q., Wu, Y., Bonilla, L. S., von Buchholtz, L. J. & Ryba, N. J. P. Diversity amongst trigeminal neurons revealed by high throughput single cell sequencing. PLoS ONE 12, e0185543 (2017).

12. von Buchholtz, L. J. et al. Decoding Cellular Mechanisms for Mechanosensory Discrimination. Neuron 109, 285–298.e5 (2021).

13. Närhi, M. V., Hirvonen, T. J. & Hakumäki, M. O. Activation of intradental nerves in the dog to some stimuli applied to the dentine. Arch Oral Biol 27, 1053–1058 (1982).

14. von Buchholtz, L. J., Lam, R. M., Emrick, J. J., Chesler, A. T. & Ryba, N. J. P. Assigning transcriptomic class in the trigeminal ganglion using multiplex in situ hybridization and machine learning. Pain 161, 2212–2224 (2020).

15. Sharma, N. et al. The emergence of transcriptional identity in somatosensory neurons. Nature 577, 392–398 (2020).

16. Usoskin, D. et al. Unbiased classification of sensory neuron types by large-scale single-cell RNA sequencing. Nat Neurosci 18, 145–153 (2015).

17. Servin-Vences, M. R. et al. PIEZO2 in somatosensory neurons controls gastrointestinal transit. Cell 186, 3386–3399.e15 (2023).

18. Lam, R. M. et al. PIEZO2 and perineal mechanosensation are essential for sexual function. Science 381, 906–910 (2023).

19. Qi, L. et al. A DRG genetic toolkit reveals molecular, morphological, and functional diversity of somatosensory neuron subtypes. 2023.04.22.537932 Preprint at 10.1101/2023.04.22.537932 (2023).

20. Yarmolinsky, D. A. et al. Coding and Plasticity in the Mammalian Thermosensory System. Neuron 92, 1079–1092 (2016).

21. Szczot, M. et al. PIEZO2 mediates injury-induced tactile pain in mice and humans. Sci Transl Med 10, eaat9892 (2018).

22. Virtanen, A., Närhi, M., Huopaniemi, T. & Hirvonen, T. Thresholds of intradental A-and C-nerve fibres in the cat to electrical current pulses of different duration. Acta Physiologica Scandinavica 119, 393–398 (1983).

23. Chen, E. & Abbott, P. V. Dental Pulp Testing: A Review. Int J Dent 2009, 365785 (2009).

24. Emrick, J. J., von Buchholtz, L. J. & Ryba, N. J. P. Transcriptomic Classification of Neurons Innervating Teeth. J Dent Res 99, 1478–1485 (2020).

25. Qi, L. et al. A mouse DRG genetic toolkit reveals morphological and physiological diversity of somatosensory neuron subtypes. Cell 187, 1508–1526.e16 (2024).

26. Bhuiyan, S. A. et al. Harmonized cross-species cell atlases of trigeminal and dorsal root ganglia. Sci Adv 10, eadj9173 (2024).

27. Wanachantararak, S., Ajcharanukul, O., Vongsavan, N. & Matthews, B. Effect of cavity depth on dentine sensitivity in man. Arch Oral Biol 66, 120–128 (2016).

28. Byers, M. R. & Calkins, D. F. Trigeminal sensory nerve patterns in dentine and their responses to attrition in rat molars. Arch Oral Biol 129, 105197 (2021).

29. Stirling, C. L. et al. Nociceptor-specific gene deletion using heterozygous NaV1.8-Cre recombinase mice. PAIN 113, 27 (2005).

30. Nassar, M. A. et al. Nociceptor-specific gene deletion reveals a major role for Nav1.7 (PN1) in acute and inflammatory pain. Proc Natl Acad Sci U S A 101, 12706–12711 (2004).

31. Närhi, M. The neurophysiology of the teeth. Dent Clin North Am 34, 439–448 (1990).

32. Guo, C. et al. Pain and itch coding mechanisms of polymodal sensory neurons. Cell Rep 42, 113316 (2023).

33. Rossi, H. L. et al. Evoked and spontaneous pain assessment during tooth pulp injury. Sci Rep 10, 2759 (2020).

34. Hu, Y. et al. LabGym: Quantification of user-defined animal behaviors using learning-based holistic assessment. Cell Rep Methods 3, 100415 (2023).

35. Goss, K. et al. Quantifying social roles in multi-animal videos using subject-aware deep-learning. 2024.07.07.602350 Preprint at 10.1101/2024.07.07.602350 (2024).

36. Närhi, M., Virtanen, A., Hirvonen, T. & Huopaniemi, T. Comparison of electrical thresholds of intradental nerves and jaw-opening reflex in the cat. Acta Physiol Scand 119, 399–403 (1983).

37. Perry, S. K. & Emrick, J. Trigeminal somatosensation in the temporomandibular joint and associated disorders. Front. Pain Res. 5, (2024).

38. Brannstrom, M. The hydrodynamic theory of dentinal pain: sensation in preparations, caries, and the dentinal crack syndrome. J Endod 12, 453–457 (1986).

39. Dong, W. K., Chudler, E. H. & Martin, R. F. Physiological properties of intradental mechanoreceptors. Brain Research 334, 389–395 (1985).

40. Jones, J. et al. Selective Inhibition of NaV1.8 with VX-548 for Acute Pain. N Engl J Med 389, 393–405 (2023).

41. Lee, P. R. et al. Transcriptional profiling of dental sensory and proprioceptive trigeminal neurons using single-cell RNA sequencing. Int J Oral Sci 15, 1–14 (2023).

42. Bleicher, F. Odontoblast physiology. Experimental Cell Research 325, 65–71 (2014).

43. Abraira, V. E. & Ginty, D. D. The sensory neurons of touch. Neuron 79, 618–639 (2013).

44. Lynch, C. D. & McConnell, R. J. The cracked tooth syndrome. J Can Dent Assoc 68, 470– 475 (2002).

