## Supplementary Materials for "Intradental mechano-nociceptors serve as sentinels that prevent tooth damage"

### METHODS

### RESOURCE AVAILABILITY

#### Lead contact

#### Materials availability

This study did not generate new unique reagents.

#### Data and code availability

All data reported in this paper is available from the lead contact upon request.

All custom code will be deposited on Github and made publicly available at date of publication.

Any additional information necessary to reanalyze the data reported in this paper is available from the lead contact upon request.

### EXPERIMENTAL MODEL AND SUBJECT DETAILS

#### Animal assurance statement

All animal experiments were performed in accordance with protocols approved by the University of Michigan Institutional Animal Care and Use Committee following NIH guidelines. Experiments were performed with male and female mice. Number of mice used are indicated in the figure legends for each experiment. Mice were group housed at room temperature with ad libitum access to standard lab mouse pellet food and water on a 12 h light/12 h dark cycle. Mouse lines were purchased from Jackson or received from listed contributors, and crossed with reporter lines without additional backcrossing.

**Mouse lines**

The mouse lines used in the present study included Ai95(RCL-GCaMP6f)-D (C57BL/6J) (#028865, JAX)[1], Piezo2$^{lox/lox}$; Tac1-tagRFP-2a-TVA line[2], *Snap25-LSL-2A-EGFP-D* (#021879, JAX)[1], Na$_V$1.8-Cre (#036564, JAX)[3], CAG-LSL-Gq-DREADD (#026220, JAX)[4], C57BL/6J (#000664, JAX) and Ai65(RCF-tdT)[5], Ai32(RCL-ChR2(H134R)/EYFP) (#024109, JAX)[6]. *S100b*-Cre is an unpublished mouse line from Dr. Nicholas Ryba that was validated in this study. For the main body of this study, Ai95D transgenic mice were bred and GCaMP6f expression was induced via postnatal AAV-*Cre* injection.

### METHOD DETAILS

#### AAV Viral Delivery

To broadly induce GCaMP6f expression in trigeminal ganglia (TG) neurons in Ai95D and *Piezo2*-cKO transgenic mice, postnatal day 1-3 mouse pups were injected intraperitoneally with AAV9-Cre. Prior to injection, mouse pups were transiently anesthetized by placing pups on a plastic petri dish on ice. A Hamilton syringe (Hamilton Company, Reno, USA) was used to inject $10^{12}$ viral genomes in a 10μL volume of ssAAV-9/2-hEF1a-iCre-WPRE-bGHp(A) (Physical titer: 8.1 x 10E12 vg/ml, catalog #v225-9, University of Zurich Viral Vector Facility VVF, Zurich, CH) diluted in sterile saline. To induce GCaMP6f expression while simultaneously co-expressing mCherry in the TG of Ai95D mice for post hoc ISH following imaging, Ai95D were postnatally injected via the same protocol with ssAAV-9/2-hSyn1-chI-mCherry_2A_iCre-

WPRE-SV40p(A) (Physical titer: 5.6 x 10E12 vg/ml, catalog #v147-9, University of Zurich Viral Vector Facility VVF, Zurich, CH).

Sparse labeling of intradental neurons in Cre lines was achieved via two AAV injection approaches. A cohort of adult (6-8 week old) *S100b*-Cre mice were injected with AAV9-CAG-FLEX-tdT (Physical titer: 1-9 x 10E13 vg/ml, Vigene) bilaterally to the trigeminal ganglion by passing through the medial wall of the orbit. Virus was diluted 1:1 in 1X sterile PBS and 1.3 µL injections were delivered to each TG. Mice were anesthetized using the SomnoSuite® Low-Flow Anesthesia System (5% induction, 2.5% maintenance, Kent Scientific). Two weeks following injections, mice were euthanized, perfused, and TG and jaws were dissected and processed as described below. Sparse labeling of *S100b*+ neurons relied on variable viral transduction efficiency. For sparse labeling of intradental neurons in *Scn10a*-Cre mice, we performed neonatal AAV injections using a Brainbow viral transduction approach. Using the protocol described above for neonatal injections, pups were injected with a 10µL volume of a 1:5 dilution of pAAV-EF1α-Brainbow-invert tagBFP-eYFP-wPRE (Physical titer: 2.2 X 10E13 vg/mL, catalog #V120025, Addgene) and pAAV-Ef1a-Brainbow/mCherry/mTFP-WPRE (Physical titer: 2.2 X 10E13 vg/mL, catalog #V160749, Addgene) diluted in sterile saline. At 8 weeks of age, animals were euthanized, perfused, and TG and jaws were dissected and processed as described below. It is worth noting we observed robust expression of all four Brainbow fluorophores (BFP/YFP/mCherry/TFP) in TG; however, labeling of intradental terminal endings was limited to YFP detected using IHC. We suspect this may be due to deficiencies in terminal ending trafficking of the three other fluorophores.

To selectively induce channelrhodopsin-2/EYFP fusion protein in intradental neurons, adult (8 weeks old) Ai32(RCL-ChR2(H134R)/EYFP) mice were anesthetized and access to the maxillary or mandibular molars was achieved using the surgical protocol described for retrograde labeling below. Shallow occlusal cavitations were prepared on the right maxillary and mandibular M1 and M2 molars. Hydrophobic bonding agent was applied to surrounding tissues to prevent nonspecific transduction. A micropipette tip was used to precisely inject $10^{12}$ viral genomes to the exposed cavitations using a total volume of 2 µL of 1:1 5% silk fibroin (Advanced BioMatrix, Cat. No. 5154-20ML): ssAAV-6(F129L)/2-hEF1a-iCre-WPRE-bGHp(A) (Physical titer: 7.2 x 10E12 vg/ml, catalog #225-6(F129L)/2, University of Zurich Viral Vector Facility VVF, Zurich, CH). The total volume was delivered using 3 applications over 10 min allowing each application to gently dry then the preparation was covered with a dental filling (Flow-It ALC B2; Pentron) then light cured. Mice were allowed to recover for at least 3 weeks to ensure adequate viral-induced expression prior to optical fiber implantation (described below).

**Retrograde labeling of intradental neurons**

Retrograde labeling of intradental trigeminal neurons via cholera toxin B-subunit (CTB) reconstituted in Milli-Q water was performed as described previously[7]. Briefly, adult mice (over 6 weeks of age) were deeply anesthetized via isoflurane administration (4% for induction and 1.5-2% for maintenance using a SomnoSuite® Low-Flow Anesthesia System (Kent Scientific) administered through a secured nose cone. Ophthalmic ointment (Fisher Scientific, Catalog #NC0490117) was applied to the eyes to prevent drying. Body temperature was maintained using a hand warmer. Access to the mandibular molars was achieved via a custom device designed to separate the maxillary and mandibular incisors to open the mouth vertically, followed by oral insertion of surgical retractors horizontally to expose the molars. Shallow occlusal cavitations were prepared with a ¼ round carbide dental bur attached to a micromotor drill (variable RPM). Hydrophobic bonding agent was applied to surrounding tissues to prevent nonspecific labeling. CTB volume (up to 1 μL) was delivered using 3 applications over 15 min allowing each application to gently dry then the preparation was covered with a dental filling (Flow-It ALC B2; Pentron) then light cured. Given our previous report that 16 hours is sufficient to induce robust CTB labeling of intradental neurons, mice were allowed to recover for this amount of time prior to either harvesting TGs for ISH or conducting in vivo imaging experiments. For in vivo imaging, tooth fillings were removed immediately prior to the experiment.

**Immunohistochemistry**

Animals were perfused using 30 mL of 4ºC 1X PBS followed by 4% PFA/1XPBS. TG were dissected and fixed in 4% PFA for 2 hours then incubated overnight in 30% sucrose/1XPBS at 4ºC until they had sunk, after which they were placed in OCT, frozen with dry ice, and stored in -80ºC until sectioning. For evaluation of immune cell in response to dentin or pulp exposure surgeries were performed to expose the dentin or pulp (as described below in **Tooth Stimulation**) using the same time course as with calcium imaging experiments before perfusions. Mandibles and maxillae were dissected and post-fixed overnight in 1X PBS, 4% PFA. Samples were then demineralized for 7 days in 0.1 M EDTA at 42ºC shaking (350RPM) with buffer exchange every 2 days. After demineralization, the jaws were cryoprotected in 30% sucrose/1XPBS for 1-2 days until they had sunk, after which they were placed in OCT, frozen with dry ice, and stored in -80ºC until sectioning. OCT embedded tissues were sectioned (Leica CM1950) at 20 μm (TG) or 30 μm (hemisected whole skulls) and thaw captured onto Gelatin Subbed Slides (Southern Biotech, #SLD01-BX). Slides were stored in -80ºC until immunohistochemistry was performed. Slides were brought to room temperature then a hydrophobic barrier was drawn around the tissues using an ImmEdge Pen (Vectorlabs). Slides were post-fixed using 1X PBS/4% PFA for 10 minutes then washed with 1X PBS three times. Sections were blocked with 10% Normal Goat Serum in 0.1%Triton-X-100/1X PBS for 1 hour at RT, then incubated in primary antibody 4ºC overnight. After washing 3 times in 1X PBS, sections were then incubated in secondary antibody for 2 hr at RT in dark. Finally, sections were washed 3 times in 0.1%Triton-X-100/1X PBS then mounted in Prolong Diamond with or without DAPI (Life Technologies) coverslipped, then imaged using a confocal microscope (Olympus FV3000, Evident Scientific, Inc.).

| **Antibodies** | **Source** | **Identifier** |
|---|---|---|
| Rabbit anti-RFP (1:500) | Rockland | 600-401-379 |
| Chicken anti-GFP (1:1000) | Life Technologies | PA1-9553 |
| Sheep anti-GFP/eYFP (1:500) | Bio-Rad | 4747-1051 |
| Rat anti-CD45 (1:500) | BioLegend | 103101 |
| Goat anti-rabbit (Alexa Fluor 568) | Life Technologies | A11011 |
| Goat anti-chicken (Alexa Fluor 647) | Life Technologies | A21449 |
| Donkey anti-sheep (Alexa Fluor 488) | Life Technologies | A11015 |
| Donkey anti-rat (Alexa Fluor 647) | Invitrogen | A48272 |

***In situ* hybridization (ISH) of TG in tissue sections**

Mouse TG were freshly dissected, embedded in OCT, and flash frozen on dry ice before cryosectioning (Leica CM1950) at a thickness of 20 μm onto Superfrost Plus Slides (Fisher #12-550-15). Slides were then stored in -80°C until use. *In situ* hybridization (ISH) was performed via either a modified hybridization chain reaction (HCR) version 3 protocol[2,7,8] or a modified RNAscope protocol[9].

For the HCR method[7,8,10], buffers, hairpin amplifiers and probes against transcripts for mouse genes *S100b, Calca, Scn10a, Mrgprd, Chrna7, Fxyd2*, *mCherry*, *TdTomato, eYFP*, *Piezo2*, or *Tubb3* were purchased from Molecular Instruments. Slides were fixed in 4% PFA/1XPBS for 15 minutes on ice, then washed 3X with 1X PBS. Following fixation, sections were acetylated with 0.3% v/v acetic anhydride and 0.1M triethanolamine in Milli-Q water and washed 3X in 1X PBS before undergoing dehydration through an ethanol series (50%, 70%, 100%, fresh 100%; 5 minutes each) then rinsed 3X in 2X saline-sodium citrate (SSC). Next, slides were prehybridized with hybridization buffer using a Coverwell chamber (Grace Biolabs) at 37°C for 10 min in a humidified hybridization oven. This was followed by incubation with a working hybridization solution (prepared by adding probes with adapters B1-B6 at a concentration of 4 nM per probe to pre-warmed hybridization buffer at 37°C), and incubated for 1-3 days in a humidified hybridization oven at 37°C. Following hybridization, slides were washed in 100% wash buffer for 5 min at 37°C then underwent a wash solution series (75% wash buffer in 5x SSC containing 0.1% Tween 20 (SSCT), 50% wash buffer in 5x SSCT, 25% wash buffer in 5x SSCT, 100% 5x SSCT for 10 min at 37°C) with a final wash in 100% 5x SSCT for 5 min at room temperature (RT). Next, slides were pre-incubated in amplification buffer for 30 min followed by overnight incubation in a working amplification solution, at RT in a humidified chamber protected from light. The working amplification solution consisting of amplification buffer containing amplifier hairpins for adapters B1-B6 conjugated to fluorophores was freshly prepared according to manufacturer recommendations, ensuring the selected associated fluorophores (Alexa488, Alexa561, Alexa594, Alexa647, Alexa750) had no overlap with any endogenous fluorescence present in the sample for each experiment. Finally, slides were washed 2x in 5X SSCT for 15 minutes with gentle agitation then mounted in Imaging Buffer (3 U/ml pyranose oxidase, 0.8% D-glucose, 2X SSC, 10 mM Tric HCl pH 7.4, 400 U RNAse inhibitor), coverslipped, and sealed for imaging using Cytobond (Scigene).

For RNAscope of retrograde labeled TG, all reagents and probes against transcripts for mouse genes *S100b*, *Smr2,* or *Bmpr1b* were purchased from ACD bio. RNAscope was performed based on modified published protocols[9] and following ACD Bio manufacturer guidelines. Briefly, slides were removed from -80°C storage and fixed for 1 hour with freshly made 4% PFA in 1X PBS precooled to 4°C on ice. Next, slides were rinsed twice with 1X PBS to remove excess fixative, then covered in 1X PBS, coverslipped, and imaged using confocal microscopy for retrograde labeling of CTB-AF647. We chose to screen for CTB at this time as we noticed considerable loss of CTB-AF647 signal following later Protease IV treatment. Tissue sections were protected from damage using spacers created from strips of electrical tape (200μm thick) that were adhered to the perimeter of coverslips prior to mounting. Following confocal imaging for CTB, coverslips were gently removed and slides were briefly rinsed in 1X PBS before undergoing dehydration through an ethanol series (50%, 70%, 100%, fresh 100%; 5 minutes each). After dehydration, slides were removed from 100% ethanol and allowed to air dry for 5 minutes at RT. A hydrophobic barrier was drawn around each section using an ImmEdge hydrophobic barrier pen (Vectorlabs). Slides were then rehydrated in 2X SSC and washed 2X in 2X SSC, then incubated with Protease IV for 10 minutes, followed by two washes in 2X SSC. Next, slides were incubated in a mix of RNAscope probes for 2 hours in a humidified hybridization oven at 40°C. Slides were then washed 2X in Wash Buffer for 2 min each at RT, followed by adding and washing Amp1, Amp2, and Amp3 then developing HRP C1 and C2 signals according to manufacturer instructions. Finally, slides were mounted using Prolong Gold Antifade Mountant with DAPI (Thermofisher, P36935), coverslipped, and imaged using a confocal microscope (Olympus FV3000, Evident Scientific, Inc.). Image alignment was performed using ImageJ prior to scoring for CTB and ISH overlap.

**In vivo epifluorescence $Ca^{2+}$ imaging of the trigeminal ganglion**

In vivo $Ca^{2+}$ imaging of trigeminal neurons was performed in anesthetized mice using previously described methods with slight modifications[11]. Briefly, mice were deeply anesthetized via isoflurane administration (4% for induction and 1.5-2% for maintenance using a SomnoSuite® Low-Flow Anesthesia System (Kent Scientific) administered through a secured nose cone. Ophthalmic ointment (Fisher Scientific Catalog #NC0490117) was applied to the eyes to prevent drying. Body temperature was maintained using a hand warmer. Whiskers were trimmed to ensure that unintentional deflection responses of trigeminal neurons did not occur during imaging. Mice were head-fixed using a custom stereotaxic apparatus that stabilized the skull while enabling access to the oral cavity. Optical access of the trigeminal ganglion surface was achieved via bilateral hemispherectomy as described previously[11]. Following trigeminal ganglion exposure and establishment of hemostasis, a spring-loaded wire was placed in between the maxillary and mandibular incisors to open the mouth via applying gentle tension vertically, followed by positioning of insulated retractors horizontally to expose the mandibular molars. $Ca^{2+}$ imaging was performed within 1 hr of trigeminal ganglion exposure using a Thorlabs custom-built epifluorescence microscope with a 4x, 0.16 NA Olympus objective. Two Ultra-High-Power LED Controllers (Prizmatix) were used to image GCaMP6f (480 nm) and RFP (561 nm) and were filtered using a GFP filter sets (Thorlabs) and tdTomato filter sets for RFP (Thorlabs). Image acquisition was performed using a pco.panda

4.2 bi scientific CMOS camera at 5 Hz. For all experiments in which no post-hoc in situ hybridization was performed, a piezoelectric actuator (NanoScan NPC-D-6111, Prior) was used to achieve highly precise vertical linear movement of the Olympus objective in order to capture a deeper field of view (300 µm) of the surface of the ganglion. To verify Piezo2-cKO in live imaging experiments, gentle stroking of the exterior mouse cheek with the grain of the hairs via a paintbrush was performed similar to previous reports[2,12]. Care was taken to ensure each stroke was consistent in gentle applied force resulting in slight deflection of hair. Cheek pinch was performed using forceps to pinch the external cheek skin.

**Tooth stimulation**

***Electrical stimulation*** was used to identify intradental neurons using a pulse generator (Koolertron DDS Signal Generator). For typical experiments: An insulated wire cathode was placed in the nearby lingual gingiva (see Figure S1). The tip of an insulated wire anode (+) was held on the molar occlusal surface. Following anode placement, a square waveform set to 2-4V, 0.6 Hz, 1-2V offset, and 12% duty pulse wave was delivered to a single molar. The anode was moved to the adjacent molar without changing the cathode position. Following the stimulation of both molars, the cathode position was changed. Anode stimulation proceeded as described. This process was repeated for 3 independent rounds with different cathode positions. Intradental neurons were defined as those cells that responded to at least 2 of 3 rounds of electrical stimulation. Cold application was applied to m1 and/or m2 molars as indicated via application of Hygenic® Endo-Ice® Pulp Vitality Refrigerant Spray (Coltene) that was sprayed on a small spherical cotton pellet (1 mm diameter) for 5 seconds then immediately applied. ***Enamel cutting*** was performed by application of a ¼ round carbide dental bur (Munce discovery burs, CJM Engineering) driven by a cordless micromotor drill

(35,000 RPM) to the occlusal surface of individual molars in a posterior to anterior sweeping motion. Friction stimulus was delivered using the same approach using a rounded “fluteless” carbide dental bur that was smoothed using sandpaper to generate diminutive damage to the molar occlusal surface. Care was taken to ensure each motion and the downward force was consistent between friction and cutting trials in series in the same mouse. In experiments, a single molar was the target of enamel cutting, but the relative size of the bur to the tooth sometimes resulted in hitting the surface of the adjacent molar. Degree of damage to the enamel occlusal surface depended on whether the bur was standard or fluteless (see Figure S5A). ***Vibration*** was delivered using a custom applicator that held a dental file placed in contact with the surface of a single tooth as previously reported[11]. Briefly, sinusoidal waveforms at 50, 75, 100, 150, 200 Hz lasting 3 sec for each were generated using a Digidata 1550 (Molecular Devices) to control a solenoid (Solid Drive SD1sm, Induction Dynamics) and were amplified by a 70W subwoofer plate amplifier (SA70, Dayton Audio). ***Direct forces*** were applied to the occlusal surface of individual molars via von Frey filaments (0.4g and 2g, Ugo Basile) or a dental file (Dentsply) with an applied force that was measured at 20g. For pulp/dentin exposure experiments, occlusal cavitations exposing the dentin and/or pulp were made just before or during live imaging depending on the experimental condition. Pulp/dentin exposure was performed based on our previously published protocol for retrograde labeling of intradental neurons[7] and as described above. We defined pulp exposure when drilling depth resulted in presentation of stereotypical visual cues including pink to red coloration and fluid emanating from the pulp bed. Conversely, dentin drilling was achieved via drilling through the enamel while avoiding exposure depths that gave rise to pulp exposure visual cues. Depth of drilling was carefully assessed for each tooth under high magnification with a stereoscope.

***Pulp temperature*** was measured during enamel cutting (including friction). Mice were anesthetized and access to the mandibular molars was achieved using protocols previously described for live imaging. A tunnel was drilled into the mesial side of the m1 molar using a needle carbide dental bur (Dentsply) driven by a cordless micromotor drill operating at 35,000 RPM. Drilling extended into the inner pulp chamber while keeping the occlusal tooth structure intact. A T-Type thermocouple (5SC-TT-T-40-36, Omega Thermocouples) was then implanted into the dental pulp. The thermocouple was sealed with Flow-It ALC B2 (Pentron) and light-cured to secure the preparation. Temperature readings were captured using a temperature reader from Sable Systems International (TC-2000 Type T Thermocouple Meter), capable of measuring temperatures from -100°C to +125°C with a resolution of 0.01°C. Data collected by the thermocouples were converted for computer analysis using a LABTRAX 4-Channel Data Acquisition system from World Precision Instruments (WPI Data Acquisition). Data analysis and recording was performed with LabScribe V4 software (IWORX). The standard temperature recording protocol involved a baseline measurement for 30 seconds, followed by enamel cutting applied for 20 seconds (between 30 and 50 seconds into recording), and continued recording up to 2 minutes. This protocol enabled reproducible evaluation of thermal dynamics of the inner tooth and pulp in response to enamel cutting.

### Analysis

*Motion Correction of In Vivo $Ca^{2+}$ Response Time-Lapse Image Data*

Inter-frame movement correction was performed using an in-house MATLAB code (Tomer Stern). The algorithm initially assumes translation alignment for a rough approximation, followed by a precise affine alignment fine tuning. Both steps utilize mean squares as the similarity metric and employ regular step gradient descent for optimization.

*Fluorescence Dynamics*

Manual regions of interest (ROIs) were drawn around responding neurons using the freehand tool in ImageJ. Through a custom Matlab script[2], normalized GCaMP6f fluorescence ($\Delta F/F_0$) was calculated for each ROI. To correct for fluctuations in background signal, a neuropil region local to the manually drawn somatic ROI was created to enable spatial averaging of the $Ca^{2+}$ response then subtraction from the somatic average. This process was repeated for every frame and corrects for all contaminant signals (i.e., underlying, out of focus, and neighboring cell fluorescence) throughout the experiment.

*Background Subtraction*

Images were background subtracted using ImageJ's Subtract Background Feature. First, a new image was generated in which the rolling ball pixel radius was set between 10-30 pixels, and the subsequent output image was subtracted from the initial raw image. All pseudocoloring was also performed using ImageJ's lookup table (LUT) feature. Additionally, other video timelapses were $\Delta F$ background subtracted. Here, the first ten frames were averaged using ImageJ's Z Project feature. This image was then subtracted from the initial video using the Image Calculator feature.

*AUC*

Area under the curve measurements were made using GraphPad Prism. In particular, the measurements for AUC were taken as *total* area under the curve. There was no user preference for peak intensity or peak width. Simply, the area under the curve was set to be the area under the normalized intensity values during time of stimulation (frames 40-120).

*Proportions*

Proportions were calculated on GraphPad Prism as a function of the number of responses of all voltage responders. Responders were identified using known time of stimulus application (typically around frame 40).

*Somal diameter*

Intradental neuron somal diameter was calculated using Feret's diameter for each ROI (ImageJ).

**Post hoc ISH and alignment and background subtraction**

*Whole-mount ISH of trigeminal ganglion*

Following live imaging, TG in the skulls were immediately washed with PBS to remove residual blood, followed by brief fixation in ice cold 4% PBS/1X PBS. TG were then dissected and processed for whole mount ISH using HCR as previously described[2]. Briefly, ganglia were immersed in 4% PFA/1X PBS on ice for 90 min, rinsed 3 times in 1X PBS, adhered dorsal surface exposed to a small plastic strip (10 x 2mm) to aid handling, then rinsed once more in 1X PBS. ISH was performed via a modified protocol using hybridization chain reaction version 3[2,8]. Pre-hybridization: 30 min, 35°C in HCR hybridization buffer (Molecular Instruments); Hybridization: 48-72 hr 35° combining the following probes: *d2eGFP* (detects GCaMP expression)*, mCherry, S100b, Calca, Scn10a, Mrgprd*. Following hybridization, ganglia were washed: 1) 2 x 15 min in wash buffer (Molecular Instruments) at 39°C, 2) 4 x 30 min in 30% formamide/2xSSC at 39°C 3) 2x in 2x SSC at RT (briefly), 4) 1 x 30 min in amplification buffer (Molecular Instruments) at RT rotating in dark, then amplified overnight rotating in dark in amplification buffer (Molecular Instruments) containing amplifier hairpins for adapters B2 - B6 conjugated to AF750, AF488, AF546, AF594, or AF647 prepared as directed by the manufacturer. Following amplification, ganglia were washed 4 x 30 min in 5xSSC/0.1% Tween rotating in dark, then embedded in 2% low-melt agarose filled halfway in a CoverWell imaging chamber (Grace Biolabs). Once agar solidified, the remainder of the chamber was filled with Imaging Buffer (3U/mL pyranose oxidase, 0.8% D-glucose, 2xSSC, 10mM Tris-HCL ph 7.4) containing 10µl RNase inhibitor (New England Biolabs), and ganglia were imaged immediately using an Olympus FV3000 confocal microscope.

*Alignment of Whole-Mount ISH, $Ca^{2+}$ Response, and In Vivo RFP Image Data*

For aligning the three datasets (in vivo mCherry, ISH, and $Ca^{2+}$ response), the in vivo mCherry image was designated as the fixed source, while the other two datasets were considered as moving targets. The mCherry reference image was taken prior to the beginning of $Ca^{2+}$ imaging (GCaMP6f) in which there is minimal movement of the ganglion. The alignment process involved two main steps:

1. Intra-dataset Alignment: Each dataset consisting of multiple slices was internally aligned. The $Ca^{2+}$ response slices were aligned as detailed in “Motion Correction of In vivo $Ca^{2+}$ Response Time-Lapse Image Data.” For ISH, channels 1, 2, and 4 were aligned to channel 3 using translation alignment, with mean squares as the similarity metric and regular step gradient descent for optimization.
2. Inter-dataset Alignment:
    a. ISH to mCherry Alignment: Initially, 8-10 pairs of control points (i.e. guideposts) were manually marked between the second ISH channel and the mCherry image. These points were used to fit a second-order polynomial transformation

for a preliminary alignment. Subsequently, a non-rigid alignment was performed using the Demons algorithm with iterations set at [5,1,1,1] across pyramid levels [4,3,2,1], respectively. Prior to this, intensity calibration was done by rescaling the 0.1 and 99.9 percentile values of each image to a 0-1 range. Lastly, both inferred polynomial and non-rigid alignment were also applied to ISH channels 1, 3, and 4.

b. $Ca^{2+}$ Response to mCherry Alignment: This involved aligning a maximum projection (along the time axis) of the $Ca^{2+}$ response time-lapse to the mCherry data using translation alignment. Mattes Mutual Information was used as the metric, with the One-Plus-One evolutionary optimizer. Similar to the ISH alignment, this step was preceded by percentile-based intensity calibration. The resulting translation alignment was then applied to each frame in the time-lapse series.

**Electrical and optogenetic stimulation and EMG measurements**

*Fiber optic and muscle electrode implantation, optogenetic stimulation, and EMG recording*

Four weeks post-injection of AAV6/2-hEF1a-iCre into the right-side molars of Ai32(RCL-ChR2(H134R)/EYFP) mice, a fiber optic was implanted on top of the right TG. After preemptive administration with 5mg/kg of analgesic carprofen, mice were anesthetized with isoflurane (4 - 5% for induction, 1.5% for maintenance) before being secured in the stereotactic frame (David Kopf Instruments). Body temperature was maintained at 35 – 37 °C using a feedback-controlled heating pad (Physitemp, TCAT-2LV). The following coordinates were used for implantation of the fiber optic: (TG) +2.4 mm from the midline; -3.6 mm posterior to the bregma, -4.7 mm ventral from the bone surface. The fiber optic (200 μm core, RWD) was affixed to the skull using dental cement (C&B Metabond, Parkell).

For the placement of electromyographic activity (EMG) electrodes, a 4-pin head-mount connected to wire electrodes was affixed to the skull with dental cement. Subsequently, all wire electrodes were positioned under the face skin and placed on the surface of each muscle. Incisions were made in the ventral part of the mandible to expose the right anterior digastric, and the right cheek to expose the masseter muscle, respectively. Two stainless steel wire electrodes (36 AWG size, Cat: 36744MHW, Phoenix wire) were inserted 2 mm apart into each muscle and then sutured with a 7-0 polypropylene surgical suture (Prolene TM, Ethicon). One week later, EMG signals of the anterior digastric and masseter muscle were recorded from awake animals by laser pulse administration (15 or 100 ms, 1 sec) via a fiber connected to a 473 nm laser. The EMG signals were recorded using an amplifier (Quad Bio Amp; AD instruments), filtered digitally (band-pass filter at 100-3000 Hz), and integrated (time constant of 0.1 s). All the equipment were connected to PowerLab 8/35 (AD instruments) for data acquisition. Following completion of experiments, animals were euthanized, perfused, and TG and jaws were dissected,screened for ChR2-YFP expression, flash frozen on dry ice in OCT, sectioned into 20 micron slices using a cryostat, and mounted on Superfrost Plus Slides (Fisher #12-550-15) for ISH.

*IAN stimulation*

Stimulating the inferior alveolar nerve (IAN) was performed as previously reported[13,14]. Adult wildtype mice were anesthetized with urethane (1000 mg/kg), with a second dose (600 mg/kg) after 30 minutes; supplementary doses (100 mg/kg) were administered if required. EMG electrodes were implanted as described above. The IAN was reached intraorally and a monopolar tungsten electrode (impedance 0.1-0.5 MΩ; Microprobes) was inserted into the right mandibular canal to access the IAN, with a reference electrode placed on the peripheral skin close to the stimulation site. IAN was electrically stimulated (pulse width, 100 μs; frequency, 1 Hz; intensity, 1 mA) using a stimulus isolator (FE-180; AD instruments).

*Data processing and analysis*

EMG data was analyzed using Labchart 8 software (AD Instruments). The reflex latency was determined by calculating the time from the onset of the optogenetic or electrical stimulus to the onset of the reflex. The reflex duration was measured from the onset to the end of the reflex, while amplitude was defined as the peak-to-peak amplitude of the elicited EMG trace.

**Behavioral analysis of intradental neuron activation using chemogenetics**

*Scn10a*-cre; CAG-LSL-Gq-DREADD KI mice and WT controls were habituated individually in a Plexiglas acrylic chamber (3.78” inner diameter x 6” height) equipped with a custom built controllable rotating stage. A camera was positioned horizontally facing the stage to capture mouse behaviors and facial changes (Logitech 920x, 1080P, 30 fps). Mice were placed in the testing chamber directly from their home cage and video recorded to capture natural behaviors. The stage was gently rotated as needed to ensure we were obtaining video footage with the face directed forward the camera. Frames where the animal was facing away or during rotation were removed from videos. Habituation took place for at least three sessions. Following habituation, each animal underwent a total of 4 experimental paradigms on separate days. The same cohort of mice was assayed for all conditions for each genotype. For baseline recordings, animals were placed directly from their home cage and recorded. This enabled us to categorize the major baseline animal behaviors in our chamber paradigm: **locomotion** (movement involving turning or repositioning within the chamber while on all four paws, **rearing** (transitioning from sitting/standing position with all paws on the ground to standing on hindpaws, or vice versa), **grooming** (body grooming consisting of licking the body below the neck, or face grooming consisting of using the paws to groom the face, head, mouth, and/or ears), and **resting** (mouse either remains in place with minor movements or is completely motionless, with ears upright and facing forward). To evaluate behavioral responses to global peripheral *Scn10a*+ neuron activation, chemogenetic activation of Gq-DREADD was achieved using clozapine N-oxide (CNO) (Bioteche, #4936) suspended in 1X PBS. CNO was injected intraperitoneally at a dose of 0.1 mg/kg, using 31-gauge BD insulin syringes on gently restrained mice. Following injection, mice were returned to their home cages for a 25-minute acclimation period before being placed in the recording chamber and filmed enabling capture and analysis of 10 minutes of recording. *Scn10a*-cre; CAG-LSL-Gq-DREADD KI mice exhibited pain-related behaviors which were used to categorize **pain** (postural hunching, orofacial grimace consisting of ears pulled apart and back from baseline position and/or orbital tightening). We then applied LabGym[15], a recently developed automated tool for identifying and quantifying user-defined behaviors, for subsequent analysis. First, we selected 50 frames that were generated from video recordings using LabGym and annotated the mice in these frames with Roboflow (Roboflow: Computer vision tools for developers and enterprises). We used the annotated frames to train a LabGym Detector to detect the mice in all video recordings. Next, we used the trained Detector to generate behavior examples from video recordings and sort them into different behavior categories. The sorted behavior examples were then used to train a LabGym Categorizer for behavior classification. Finally the trained Detector and Categorizer were used to automatically identify and quantify the identified behaviors (locomotion, rearing, grooming, resting, and pain). Experimenters were blinded to conditions and genotypes within each of the analyzed recordings. To assess behavior in response to selective chemogenetic activation of intradental neurons, mice were briefly anesthetized via isoflurane administration (4% for induction and 1.5-2% for maintenance using a SomnoSuite® Low-Flow Anesthesia System (Kent Scientific) administered through a secured nose cone. Body temperature was maintained using a hand warmer. Access to the mandibular molars was gently achieved via a custom device designed to separate the maxillary and mandibular incisors to open the mouth vertically, followed by oral insertion of surgical retractors horizontally to expose the molars. Shallow occlusal cavitations to unilateral mandibular first and second molars to expose superficial dentin were prepared with a ¼ round carbide dental bur attached to a micromotor drill (variable RPM) followed by application of temporary fillings (Flow-It ALC B2; Pentron) then light cured to seal the preparation for an 8-hour recovery period. Following recovery, mice were again briefly anesthetized using the described procedure to access the mandibular molars, fillings were removed, CNO (0.01 mg/kg solution, 1.5 µL) was directly applied to exposed dentin until fully absorbed, fillings were replaced and light cured, and animals were allowed to wake and acclimate in their home cage for 25 minutes prior to behavior recording. To rule out potential off target systemic behavioral effects of CNO tooth application, behavior was also examined in response to i.p. injection of an equivalent dose of CNO (0.01 mg/kg) in mice prior to performing the tooth CNO experiment. Data analysis and statistics were performed using LabGym.

**Optogenetic stimulation of jaw opening reflex**

Mice (*Scn10a*-Cre; Ai32, 8-12 weeks) were anesthetized via isoflurane administration (4% for induction and 0.5-1.5% for maintenance) using a SomnoSuite® Low-Flow Anesthesia System (Kent Scientific) administered through a secured nose cone. Mice were held at a level of anesthesia to achieve stable breathing marked by no jaw movement at baseline. Recordings were performed via TTL trigger (Doric, OTPG8) for the synchronization of laser stimulation with frame acquisition using a camera (FLIR, Blackfly S camera) aimed at the opening of the oral cavity. An optical fiber (500um, NA 0.63, 5 mm length, Goldstone Scientific) was directed at 90º to target oral cavity tissues to deliver blue (470 +/- 6 nm, average intensity 564.8 $mW/cm^2$) or green light (545 +/- 6 nm, average intensity 548.8 $mW/cm^2$) generated by LED light sources (UHT-P-470-SR or UHT-P-545-SR, Goldstone Scientific). Pulses were delivered in alternating trains of green and blue light (10 pulses, 0.5 Hz, 1 sec). Intradental HTMRs were selectively targeted by directing the optical fiber at the maxillary M1. To ensure the jaw opening was specific to activation of *Scn10a*+ intradental HTMRs, the stimulation paradigm was repeated for two adjacent tissues, the hard palate and buccal vestibule. The camera was configured to record in secondary trigger mode through a custom Python script adapted from the following repository: https://github.com/neurojak/pySpinCapture. The field of view was illuminated using an LED array (850 nm, CMIR-110, CM Vision) and the lens was equipped with an IR filter (Edmund Optics, Stock #21-697) to avoid capturing light from the optical fiber. Video was acquired at 170Hz, and the trigger source for the alignment of video with light stimulation was sampled at 10kHz. A DeepLabCut network was trained to detect the position of the top and bottom incisors of the mouse to measure the amplitude of the mandibular deflections. The Euclidean distance between incisors was calculated and aligned to the laser light trace with sub-millisecond precision in Python.

**Quantification and statistical analysis**

Statistical analyses were performed using GraphPad prism. Statistical methods, error bars, and sample sizes were described in the figure legends. Broadly, ANOVA with post hoc analysis was performed for multi-group comparisons, whereas t test (paired or unpaired as indicated) was used to compare two groups. Statistical significance was defined as P values <0.5.

To validate transduction efficiency of Ai95D mice injected with AAV-Cre at P0, TG sections underwent ISH for *GCaMP* and *Tubb3*. In ImageJ, ROIs for individual neurons were drawn based on *Tubb3* expression, followed by manual scoring for the presence of *GCaMP*, with *GCaMP*+ positive cells defined based on visible presence of fluorescent signal above background throughout the ROI. Transduction efficiency was calculated as the percent of GCaMP+ cells/*Tubb3*. Somal diameter for GCaMP+ versus GCaMP- cells was also calculated using Feret's diameter for each ROI (ImageJ).

To validate knockout efficiency in *Piezo2* cKO mice, TG sections from Piezo2-cKO; Ai95D or control Ai95D mice injected with AAV-Cre at P0 underwent ISH for *GCaMP*, *Piezo2,* and *Tubb3*. Individual neurons from imaged sections were analyzed for the presence of *GCaMP* and *Piezo2*. In ImageJ, ROIs for individual neurons were drawn based on *Tubb3* expression, followed by manual scoring for the presence of *GCaMP*, with *GCaMP*+ positive cells (indicating Cre recombination) defined based on visible presence of fluorescent signal above background throughout the ROI. Next, ROIs taken from both *Piezo2-cKO* and control Ai95D mice were blindly shuffled using a custom MatLab application and manually evaluated for *Piezo2* expression. ROIs were scored as positive for *Piezo2* based on the presence of diffuse fluorescent puncta throughout the cytosol.

*Quantification of immune cell fluorescent intensity*

Fluorescence quantification of immune cells was performed using a MATLAB-based image processing workflow. Images were first pre-processed by excluding any bright areas outside the region of interest—the pulp and surrounding dentin of the tooth. The DAPI channel, due to its brightness and clear delineation, was used to define ROIs. Thresholding was applied to identify these regions, followed by morphological operations to clean up noise by removing small objects and filling holes. Once ROIs were identified, the immune cell fluorescence signal was normalized by subtracting the mean intensity of non-ROI background areas, ensuring consistent signal comparisons across images. For enhanced accuracy, background fluorescence was subtracted. This involved thresholding the background channel to identify significant areas of noise and subtracting these from the immune cell channel. Fluorescence intensity was quantified by calculating the average brightness within the ROIs, achieved by dividing the total fluorescence intensity of the ROIs by their total area. This approach enabled precise quantification while accounting for variations in background intensity and ensuring consistency across different images. Visual checks were implemented throughout the process to ensure accuracy in thresholding.

*Specificity/efficiency calculation*

Cell segmentation in microscopy images of the trigeminal ganglion was performed with the aid of Cellpose, a neural network for biological segmentation tasks. The *cyto* pre-trained model was tuned to our dataset using the Cellpose human-in-the-loop functionality for significant gains in segmentation accuracy[16]. Cellpose masks were manually filtered and adjusted to remove any background artifacts or to fix segmentation borders. The masks were then visually inspected across fluorescence channels before calculating labeling efficiency and specificity.

The efficiency metric reflects the proportion of labeled neurons within an imaging field, while the specificity metric reflects a fluorophore's fidelity to a molecular marker.

#### Visualization of *Scn10a*+ neuron central projections

The animal was perfused with 1X PBS (phosphate-buffered saline) followed by 4% paraformaldehyde (PFA) to fix the neural tissue, ensuring preservation of the brain's structural integrity. After perfusion, the brain was carefully dissected out while keeping the brainstem intact, a crucial step for any subsequent analyses involving brainstem structures. Once dissected, the brain was immersed in 4% PFA for post-fixation overnight, allowing complete tissue fixation. This step is vital for maintaining cellular details and preventing degradation. After fixation, the brain was transferred to a solution of 30% sucrose in 1X PBS, which functions as a cryoprotectant. The brain remains in this solution until it sank, indicating sufficient infiltration (usually about 2-3 days). For long-term preservation, the brain was embedded in an OCT (optimal cutting temperature) compound within a mold, labeled with the sample name, date, and experiment details, and then stored at -80°C until sectioning and further analysis. For sectioning, we set a thickness of 50 µm for each section and collected the serial sections onto gelatin-coated slides. The slides were stored at -80 until they were ready for imaging and visualization of YFP using confocal microscopy (Olympus FV3000, Evident Scientific, Inc.).

### Appendix

#### Table 1: Statistics for 4G Pain Duration.

Statistics for pain duration using One-way ANOVA with Tukey's correction.

**Pain Duration**

| Tukey's multiple comparisons test | Mean Diff. | 95.00% CI of diff. | Below threshold? | Summary | Adjusted P Value |
|---|---|---|---|---|---|
| Baseline KI vs. 0.01 mg/kg CNO (i.p.) KI | -82.86 | -174.1 to 8.410 | No | ns | 0.1013 |
| Baseline KI vs. 0.1 mg/kg CNO (i.p.) KI | -527.7 | -618.9 to -436.4 | Yes | **** | <0.0001 |
| Baseline KI vs. 0.01 mg/kg CNO (tooth) KI | -472 | -563.3 to -380.7 | Yes | **** | <0.0001 |
| Baseline KI vs. Baseline WT | 10 | -81.27 to 101.3 | No | ns | >0.9999 |
| Baseline KI vs. 0.01 mg/kg CNO (i.p.) WT | -12.5 | -103.8 to 78.78 | No | ns | 0.9999 |
| Baseline KI vs. 0.1 mg/kg CNO (i.p.) WT | -9.345 | -100.6 to 81.93 | No | ns | >0.9999 |
| Baseline KI vs. 0.01 mg/kg CNO (tooth) WT | -95.17 | -186.4 to -3.893 | Yes | * | 0.0352 |
| 0.01 mg/kg CNO (i.p.) KI vs. 0.1 mg/kg CNO (i.p.) KI | -444.8 | -536.1 to -353.5 | Yes | **** | <0.0001 |
| 0.01 mg/kg CNO (i.p.) KI vs. 0.01 mg/kg CNO (tooth) KI | -389.2 | -480.4 to -297.9 | Yes | **** | <0.0001 |
| 0.01 mg/kg CNO (i.p.) KI vs. Baseline WT | 92.86 | 1.592 to 184.1 | Yes | * | 0.0434 |
| 0.01 mg/kg CNO (i.p.) KI vs. 0.01 mg/kg CNO (i.p.) WT | 70.37 | -20.90 to 161.6 | No | ns | 0.2488 |
| 0.01 mg/kg CNO (i.p.) KI vs. 0.1 mg/kg CNO (i.p.) WT | 73.52 | -17.75 to 164.8 | No | ns | 0.2022 |
| 0.01 mg/kg CNO (i.p.) KI vs. 0.01 mg/kg CNO (tooth) WT | -12.3 | -103.6 to 78.97 | No | ns | 0.9999 |
| 0.1 mg/kg CNO (i.p.) KI vs. 0.01 mg/kg CNO (tooth) KI | 55.63 | -35.64 to 146.9 | No | ns | 0.5442 |
| 0.1 mg/kg CNO (i.p.) KI vs. Baseline WT | 537.7 | 446.4 to 628.9 | Yes | **** | <0.0001 |
| 0.1 mg/kg CNO (i.p.) KI vs. 0.01 mg/kg CNO (i.p.) WT | 515.2 | 423.9 to 606.4 | Yes | **** | <0.0001 |
| 0.1 mg/kg CNO (i.p.) KI vs. 0.1 mg/kg CNO (i.p.) WT | 518.3 | 427.0 to 609.6 | Yes | **** | <0.0001 |
| 0.1 mg/kg CNO (i.p.) KI vs. 0.01 mg/kg CNO (tooth) WT | 432.5 | 341.2 to 523.8 | Yes | **** | <0.0001 |
| 0.01 mg/kg CNO (tooth) KI vs. Baseline WT | 482 | 390.7 to 573.3 | Yes | **** | <0.0001 |
| 0.01 mg/kg CNO (tooth) KI vs. 0.01 mg/kg CNO (i.p.) WT | 459.5 | 368.3 to 550.8 | Yes | **** | <0.0001 |
| 0.01 mg/kg CNO (tooth) KI vs. 0.1 mg/kg CNO (i.p.) WT | 462.7 | 371.4 to 553.9 | Yes | **** | <0.0001 |
| 0.01 mg/kg CNO (tooth) KI vs. 0.01 mg/kg CNO (tooth) WT | 376.9 | 285.6 to 468.1 | Yes | **** | <0.0001 |
| Baseline WT vs. 0.01 mg/kg CNO (i.p.) WT | -22.5 | -113.8 to 68.78 | No | ns | 0.9937 |
| Baseline WT vs. 0.1 mg/kg CNO (i.p.) WT | -19.35 | -110.6 to 71.93 | No | ns | 0.9975 |
| Baseline WT vs. 0.01 mg/kg CNO (tooth) WT | -105.2 | -196.4 to -13.89 | Yes | * | 0.0134 |
| 0.01 mg/kg CNO (i.p.) WT vs. 0.1 mg/kg CNO (i.p.) WT | 3.15 | -88.12 to 94.42 | No | ns | >0.9999 |
| 0.01 mg/kg CNO (i.p.) WT vs. 0.01 mg/kg CNO (tooth) WT | -82.67 | -173.9 to 8.602 | No | ns | 0.1029 |
| 0.1 mg/kg CNO (i.p.) WT vs. 0.01 mg/kg CNO (tooth) WT | -85.82 | -177.1 to 5.452 | No | ns | 0.0797 |

**Table 2. Statistics for S11A Resting Duration.**

Statistics for resting duration using One-way ANOVA with Tukey’s correction.

**Resting Duration**

| Tukey's multiple comparisons test | Mean Diff. | 95.00% CI of diff. | Below threshold? | Summary | Adjusted P Value |
|---|---|---|---|---|---|
| Baseline KI vs. 0.01 mg/kg CNO (i.p.) KI | -124.2 | -239.3 to -9.014 | Yes | * | 0.026 |
| Baseline KI vs. 0.1 mg/kg CNO (i.p.) KI | 123.6 | 8.437 to 238.8 | Yes | * | 0.0271 |
| Baseline KI vs. 0.01 mg/kg CNO (tooth) KI | 92.46 | -22.70 to 207.6 | No | ns | 0.2055 |
| Baseline KI vs. Baseline WT | 24.46 | -90.71 to 139.6 | No | ns | 0.9975 |
| Baseline KI vs. 0.01 mg/kg CNO (i.p.) WT | 7.858 | -107.3 to 123.0 | No | ns | >0.9999 |
| Baseline KI vs. 0.1 mg/kg CNO (i.p.) WT | -21.76 | -136.9 to 93.40 | No | ns | 0.9988 |
| Baseline KI vs. 0.01 mg/kg CNO (tooth) WT | -105.2 | -220.3 to 10.01 | No | ns | 0.0975 |
| 0.01 mg/kg CNO (i.p.) KI vs. 0.1 mg/kg CNO (i.p.) KI | 247.8 | 132.6 to 362.9 | Yes | **** | <0.0001 |
| 0.01 mg/kg CNO (i.p.) KI vs. 0.01 mg/kg CNO (tooth) KI | 216.6 | 101.5 to 331.8 | Yes | **** | <0.0001 |
| 0.01 mg/kg CNO (i.p.) KI vs. Baseline WT | 148.6 | 33.47 to 263.8 | Yes | ** | 0.0036 |
| 0.01 mg/kg CNO (i.p.) KI vs. 0.01 mg/kg CNO (i.p.) WT | 132 | 16.87 to 247.2 | Yes | * | 0.0142 |
| 0.01 mg/kg CNO (i.p.) KI vs. 0.1 mg/kg CNO (i.p.) WT | 102.4 | -12.75 to 217.6 | No | ns | 0.1157 |
| 0.01 mg/kg CNO (i.p.) KI vs. 0.01 mg/kg CNO (tooth) WT | 19.02 | -96.14 to 134.2 | No | ns | 0.9995 |
| 0.1 mg/kg CNO (i.p.) KI vs. 0.01 mg/kg CNO (tooth) KI | -31.14 | -146.3 to 84.02 | No | ns | 0.9891 |
| 0.1 mg/kg CNO (i.p.) KI vs. Baseline WT | -99.15 | -214.3 to 16.02 | No | ns | 0.1409 |
| 0.1 mg/kg CNO (i.p.) KI vs. 0.01 mg/kg CNO (i.p.) WT | -115.7 | -230.9 to -0.5793 | Yes | * | 0.048 |
| 0.1 mg/kg CNO (i.p.) KI vs. 0.1 mg/kg CNO (i.p.) WT | -145.4 | -260.5 to -30.20 | Yes | ** | 0.0047 |
| 0.1 mg/kg CNO (i.p.) KI vs. 0.01 mg/kg CNO (tooth) WT | -228.8 | -343.9 to -113.6 | Yes | **** | <0.0001 |
| 0.01 mg/kg CNO (tooth) KI vs. Baseline WT | -68.01 | -183.2 to 47.16 | No | ns | 0.5838 |
| 0.01 mg/kg CNO (tooth) KI vs. 0.01 mg/kg CNO (i.p.) WT | -84.6 | -199.8 to 30.56 | No | ns | 0.3052 |
| 0.01 mg/kg CNO (tooth) KI vs. 0.1 mg/kg CNO (i.p.) WT | -114.2 | -229.4 to 0.9432 | No | ns | 0.0534 |
| 0.01 mg/kg CNO (tooth) KI vs. 0.01 mg/kg CNO (tooth) WT | -197.6 | -312.8 to -82.45 | Yes | **** | <0.0001 |
| Baseline WT vs. 0.01 mg/kg CNO (i.p.) WT | -16.6 | -131.8 to 98.57 | No | ns | 0.9998 |
| Baseline WT vs. 0.1 mg/kg CNO (i.p.) WT | -46.22 | -161.4 to 68.95 | No | ns | 0.9083 |
| Baseline WT vs. 0.01 mg/kg CNO (tooth) WT | -129.6 | -244.8 to -14.45 | Yes | * | 0.0171 |
| 0.01 mg/kg CNO (i.p.) WT vs. 0.1 mg/kg CNO (i.p.) WT | -29.62 | -144.8 to 85.55 | No | ns | 0.9919 |
| 0.01 mg/kg CNO (i.p.) WT vs. 0.01 mg/kg CNO (tooth) WT | -113 | -228.2 to 2.151 | No | ns | 0.058 |
| 0.1 mg/kg CNO (i.p.) WT vs. 0.01 mg/kg CNO (tooth) WT | -83.4 | -198.6 to 31.77 | No | ns | 0.3228 |

### Table 3. Statistics for S11B Grooming Duration.

Statistics for grooming duration using One-way ANOVA with Tukey's correction.

**Grooming Duration**

| Tukey's multiple comparisons test | 95.00% CI of diff. | Below threshold? | Summary | Adjusted P Value |
|---|---|---|---|---|
| Baseline KI vs. 0.01 mg/kg CNO (i.p.) KI | -94.97 to 166.9 | No | ns | 0.988 |
| Baseline KI vs. 0.1 mg/kg CNO (i.p.) KI | 16.24 to 278.1 | Yes | * | 0.0174 |
| Baseline KI vs. 0.01 mg/kg CNO (tooth) KI | -0.1064 to 261.8 | No | ns | 0.0503 |
| Baseline KI vs. Baseline WT | -144.3 to 117.6 | No | ns | >0.9999 |
| Baseline KI vs. 0.01 mg/kg CNO (i.p.) WT | -140.9 to 121.0 | No | ns | >0.9999 |
| Baseline KI vs. 0.1 mg/kg CNO (i.p.) WT | -157.8 to 104.1 | No | ns | 0.998 |
| Baseline KI vs. 0.01 mg/kg CNO (tooth) WT | -83.48 to 178.4 | No | ns | 0.9445 |
| 0.01 mg/kg CNO (i.p.) KI vs. 0.1 mg/kg CNO (i.p.) KI | -19.74 to 242.2 | No | ns | 0.1524 |
| 0.01 mg/kg CNO (i.p.) KI vs. 0.01 mg/kg CNO (tooth) KI | -36.09 to 225.8 | No | ns | 0.3223 |
| 0.01 mg/kg CNO (i.p.) KI vs. Baseline WT | -180.3 to 81.63 | No | ns | 0.9327 |
| 0.01 mg/kg CNO (i.p.) KI vs. 0.01 mg/kg CNO (i.p.) WT | -176.9 to 84.99 | No | ns | 0.9531 |
| 0.01 mg/kg CNO (i.p.) KI vs. 0.1 mg/kg CNO (i.p.) WT | -193.8 to 68.12 | No | ns | 0.7985 |
| 0.01 mg/kg CNO (i.p.) KI vs. 0.01 mg/kg CNO (tooth) WT | -119.5 to 142.5 | No | ns | >0.9999 |
| 0.1 mg/kg CNO (i.p.) KI vs. 0.01 mg/kg CNO (tooth) KI | -147.3 to 114.6 | No | ns | >0.9999 |
| 0.1 mg/kg CNO (i.p.) KI vs. Baseline WT | -291.5 to -29.58 | Yes | ** | 0.0067 |
| 0.1 mg/kg CNO (i.p.) KI vs. 0.01 mg/kg CNO (i.p.) WT | -288.1 to -26.22 | Yes | ** | 0.0086 |
| 0.1 mg/kg CNO (i.p.) KI vs. 0.1 mg/kg CNO (i.p.) WT | -305.0 to -43.09 | Yes | ** | 0.0024 |
| 0.1 mg/kg CNO (i.p.) KI vs. 0.01 mg/kg CNO (tooth) WT | -230.7 to 31.24 | No | ns | 0.2629 |
| 0.01 mg/kg CNO (tooth) KI vs. Baseline WT | -275.1 to -13.24 | Yes | * | 0.0213 |
| 0.01 mg/kg CNO (tooth) KI vs. 0.01 mg/kg CNO (i.p.) WT | -271.8 to -9.877 | Yes | * | 0.0266 |
| 0.01 mg/kg CNO (tooth) KI vs. 0.1 mg/kg CNO (i.p.) WT | -288.7 to -26.75 | Yes | ** | 0.0083 |
| 0.01 mg/kg CNO (tooth) KI vs. 0.01 mg/kg CNO (tooth) WT | -214.3 to 47.58 | No | ns | 0.4883 |
| Baseline WT vs. 0.01 mg/kg CNO (i.p.) WT | -127.6 to 134.3 | No | ns | >0.9999 |
| Baseline WT vs. 0.1 mg/kg CNO (i.p.) WT | -144.5 to 117.4 | No | ns | >0.9999 |
| Baseline WT vs. 0.01 mg/kg CNO (tooth) WT | -70.13 to 191.8 | No | ns | 0.8236 |
| 0.01 mg/kg CNO (i.p.) WT vs. 0.1 mg/kg CNO (i.p.) WT | -147.8 to 114.1 | No | ns | >0.9999 |
| 0.01 mg/kg CNO (i.p.) WT vs. 0.01 mg/kg CNO (tooth) WT | -73.49 to 188.4 | No | ns | 0.8618 |
| 0.1 mg/kg CNO (i.p.) WT vs. 0.01 mg/kg CNO (tooth) WT | -56.62 to 205.3 | No | ns | 0.6312 |

**Table 4. Statistics for S11C Locomotion Duration.**

Statistics for locomotion duration using One-way ANOVA with Tukey's correction.

**Locomotion Duration**

| Tukey's multiple comparisons test | Mean Diff. | 95.00% CI of diff. | Below threshold? | Summary | Adjusted P Value |
|---|---|---|---|---|---|
| Baseline KI vs. 0.01 mg/kg CNO (i.p.) KI | 70.65 | 21.49 to 119.8 | Yes | *** | 0.0008 |
| Baseline KI vs. 0.1 mg/kg CNO (i.p.) KI | 129.4 | 78.48 to 180.2 | Yes | **** | <0.0001 |
| Baseline KI vs. 0.01 mg/kg CNO (tooth) KI | 123.2 | 74.06 to 172.4 | Yes | **** | <0.0001 |
| Baseline KI vs. Baseline WT | -1.689 | -50.85 to 47.47 | No | ns | >0.9999 |
| Baseline KI vs. 0.01 mg/kg CNO (i.p.) WT | 38.44 | -10.72 to 87.60 | No | ns | 0.2329 |
| Baseline KI vs. 0.1 mg/kg CNO (i.p.) WT | 30.66 | -18.50 to 79.82 | No | ns | 0.5141 |
| Baseline KI vs. 0.01 mg/kg CNO (tooth) WT | 68.23 | 19.07 to 117.4 | Yes | ** | 0.0013 |
| 0.01 mg/kg CNO (i.p.) KI vs. 0.1 mg/kg CNO (i.p.) KI | 58.71 | 7.829 to 109.6 | Yes | * | 0.0133 |
| 0.01 mg/kg CNO (i.p.) KI vs. 0.01 mg/kg CNO (tooth) KI | 52.57 | 3.412 to 101.7 | Yes | * | 0.0281 |
| 0.01 mg/kg CNO (i.p.) KI vs. Baseline WT | -72.34 | -121.5 to -23.18 | Yes | *** | 0.0006 |
| 0.01 mg/kg CNO (i.p.) KI vs. 0.01 mg/kg CNO (i.p.) WT | -32.21 | -81.37 to 16.95 | No | ns | 0.4502 |
| 0.01 mg/kg CNO (i.p.) KI vs. 0.1 mg/kg CNO (i.p.) WT | -39.99 | -89.15 to 9.171 | No | ns | 0.192 |
| 0.01 mg/kg CNO (i.p.) KI vs. 0.01 mg/kg CNO (tooth) WT | -2.415 | -51.57 to 46.74 | No | ns | >0.9999 |
| 0.1 mg/kg CNO (i.p.) KI vs. 0.01 mg/kg CNO (tooth) KI | -6.143 | -57.03 to 44.74 | No | ns | >0.9999 |
| 0.1 mg/kg CNO (i.p.) KI vs. Baseline WT | -131 | -181.9 to -80.17 | Yes | **** | <0.0001 |
| 0.1 mg/kg CNO (i.p.) KI vs. 0.01 mg/kg CNO (i.p.) WT | -90.92 | -141.8 to -40.04 | Yes | **** | <0.0001 |
| 0.1 mg/kg CNO (i.p.) KI vs. 0.1 mg/kg CNO (i.p.) WT | -98.7 | -149.6 to -47.82 | Yes | **** | <0.0001 |
| 0.1 mg/kg CNO (i.p.) KI vs. 0.01 mg/kg CNO (tooth) WT | -61.13 | -112.0 to -10.24 | Yes | ** | 0.0085 |
| 0.01 mg/kg CNO (tooth) KI vs. Baseline WT | -124.9 | -174.1 to -75.75 | Yes | **** | <0.0001 |
| 0.01 mg/kg CNO (tooth) KI vs. 0.01 mg/kg CNO (i.p.) WT | -84.78 | -133.9 to -35.62 | Yes | **** | <0.0001 |
| 0.01 mg/kg CNO (tooth) KI vs. 0.1 mg/kg CNO (i.p.) WT | -92.56 | -141.7 to -43.40 | Yes | **** | <0.0001 |
| 0.01 mg/kg CNO (tooth) KI vs. 0.01 mg/kg CNO (tooth) WT | -54.99 | -104.1 to -5.827 | Yes | * | 0.0183 |
| Baseline WT vs. 0.01 mg/kg CNO (i.p.) WT | 40.13 | -9.031 to 89.29 | No | ns | 0.1886 |
| Baseline WT vs. 0.1 mg/kg CNO (i.p.) WT | 32.35 | -16.81 to 81.51 | No | ns | 0.4445 |
| Baseline WT vs. 0.01 mg/kg CNO (tooth) WT | 69.92 | 20.76 to 119.1 | Yes | *** | 0.0009 |
| 0.01 mg/kg CNO (i.p.) WT vs. 0.1 mg/kg CNO (i.p.) WT | -7.779 | -56.94 to 41.38 | No | ns | 0.9996 |
| 0.01 mg/kg CNO (i.p.) WT vs. 0.01 mg/kg CNO (tooth) WT | 29.79 | -19.36 to 78.95 | No | ns | 0.5506 |
| 0.1 mg/kg CNO (i.p.) WT vs. 0.01 mg/kg CNO (tooth) WT | 37.57 | -11.59 to 86.73 | No | ns | 0.2582 |

**Table 5. Statistics for S11D Normalized Locomotion Distance.**

Statistics for normalized locomotion distance using One-way ANOVA with Tukey’s correction.

**Normalized Locomotion Distance**

| Tukey's multiple comparisons test | Mean Diff. | 95.00% CI of diff. | Below threshold? | Summary | Adjusted P Value |
|---|---|---|---|---|---|
| Baseline KI vs. 0.01 mg/kg CNO (i.p.) KI | 33.25 | 12.87 to 53.62 | Yes | **** | <0.0001 |
| Baseline KI vs. 0.1 mg/kg CNO (i.p.) KI | 51.93 | 30.84 to 73.02 | Yes | **** | <0.0001 |
| Baseline KI vs. 0.01 mg/kg CNO (tooth) KI | 50.37 | 29.99 to 70.74 | Yes | **** | <0.0001 |
| Baseline KI vs. Baseline WT | -1.181 | -21.56 to 19.19 | No | ns | >0.9999 |
| Baseline KI vs. 0.01 mg/kg CNO (i.p.) WT | 15.92 | -4.456 to 36.29 | No | ns | 0.2338 |
| Baseline KI vs. 0.1 mg/kg CNO (i.p.) WT | 13.39 | -6.990 to 33.76 | No | ns | 0.4468 |
| Baseline KI vs. 0.01 mg/kg CNO (tooth) WT | 27.33 | 6.952 to 47.70 | Yes | ** | 0.0022 |
| 0.01 mg/kg CNO (i.p.) KI vs. 0.1 mg/kg CNO (i.p.) KI | 18.68 | -2.407 to 39.77 | No | ns | 0.1185 |
| 0.01 mg/kg CNO (i.p.) KI vs. 0.01 mg/kg CNO (tooth) KI | 17.12 | -3.254 to 37.50 | No | ns | 0.1615 |
| 0.01 mg/kg CNO (i.p.) KI vs. Baseline WT | -34.43 | -54.80 to -14.05 | Yes | **** | <0.0001 |
| 0.01 mg/kg CNO (i.p.) KI vs. 0.01 mg/kg CNO (i.p.) WT | -17.33 | -37.70 to 3.049 | No | ns | 0.1511 |
| 0.01 mg/kg CNO (i.p.) KI vs. 0.1 mg/kg CNO (i.p.) WT | -19.86 | -40.24 to 0.5151 | No | ns | 0.0611 |
| 0.01 mg/kg CNO (i.p.) KI vs. 0.01 mg/kg CNO (tooth) WT | -5.918 | -26.29 to 14.46 | No | ns | 0.9833 |
| 0.1 mg/kg CNO (i.p.) KI vs. 0.01 mg/kg CNO (tooth) KI | -1.562 | -22.65 to 19.53 | No | ns | >0.9999 |
| 0.1 mg/kg CNO (i.p.) KI vs. Baseline WT | -53.11 | -74.20 to -32.02 | Yes | **** | <0.0001 |
| 0.1 mg/kg CNO (i.p.) KI vs. 0.01 mg/kg CNO (i.p.) WT | -36.01 | -57.10 to -14.92 | Yes | **** | <0.0001 |
| 0.1 mg/kg CNO (i.p.) KI vs. 0.1 mg/kg CNO (i.p.) WT | -38.54 | -59.63 to -17.45 | Yes | **** | <0.0001 |
| 0.1 mg/kg CNO (i.p.) KI vs. 0.01 mg/kg CNO (tooth) WT | -24.6 | -45.69 to -3.510 | Yes | * | 0.0118 |
| 0.01 mg/kg CNO (tooth) KI vs. Baseline WT | -51.55 | -71.92 to -31.17 | Yes | **** | <0.0001 |
| 0.01 mg/kg CNO (tooth) KI vs. 0.01 mg/kg CNO (i.p.) WT | -34.45 | -54.82 to -14.07 | Yes | **** | <0.0001 |
| 0.01 mg/kg CNO (tooth) KI vs. 0.1 mg/kg CNO (i.p.) WT | -36.98 | -57.36 to -16.61 | Yes | **** | <0.0001 |
| 0.01 mg/kg CNO (tooth) KI vs. 0.01 mg/kg CNO (tooth) WT | -23.04 | -43.41 to -2.664 | Yes | * | 0.0164 |
| Baseline WT vs. 0.01 mg/kg CNO (i.p.) WT | 17.1 | -3.275 to 37.48 | No | ns | 0.1626 |
| Baseline WT vs. 0.1 mg/kg CNO (i.p.) WT | 14.57 | -5.809 to 34.94 | No | ns | 0.3381 |
| Baseline WT vs. 0.01 mg/kg CNO (tooth) WT | 28.51 | 8.134 to 48.88 | Yes | ** | 0.0012 |
| 0.01 mg/kg CNO (i.p.) WT vs. 0.1 mg/kg CNO (i.p.) WT | -2.534 | -22.91 to 17.84 | No | ns | >0.9999 |
| 0.01 mg/kg CNO (i.p.) WT vs. 0.01 mg/kg CNO (tooth) WT | 11.41 | -8.966 to 31.78 | No | ns | 0.6462 |
| 0.1 mg/kg CNO (i.p.) WT vs. 0.01 mg/kg CNO (tooth) WT | 13.94 | -6.433 to 34.32 | No | ns | 0.3938 |

**Table 6. Statistics for S11E Normalized Locomotion Speed Mean.**

Statistics for normalized locomotion speed mean using One-way ANOVA with Tukey's correction.

**Normalized Locomotion Speed Mean**

| Tukey's multiple comparisons test | Mean Diff. | 95.00% CI of diff. | Below threshold | Summary | Adjusted P Value |
|---|---|---|---|---|---|
| Baseline KI vs. 0.01 mg/kg CNO (i.p.) KI | 0.0725 | -0.003765 to 0.1488 | No | ns | 0.0736 |
| Baseline KI vs. 0.1 mg/kg CNO (i.p.) KI | 0.1114 | 0.03249 to 0.1904 | Yes | ** | 0.0011 |
| Baseline KI vs. 0.01 mg/kg CNO (tooth) KI | 0.1025 | 0.02623 to 0.1788 | Yes | ** | 0.0021 |
| Baseline KI vs. Baseline WT | -0.00875 | -0.08502 to 0.06752 | No | ns | >0.9999 |
| Baseline KI vs. 0.01 mg/kg CNO (i.p.) WT | -0.0025 | -0.07877 to 0.07377 | No | ns | >0.9999 |
| Baseline KI vs. 0.1 mg/kg CNO (i.p.) WT | 0.0075 | -0.06877 to 0.08377 | No | ns | >0.9999 |
| Baseline KI vs. 0.01 mg/kg CNO (tooth) WT | 0.0075 | -0.06877 to 0.08377 | No | ns | >0.9999 |
| 0.01 mg/kg CNO (i.p.) KI vs. 0.1 mg/kg CNO (i.p.) KI | 0.03893 | -0.04001 to 0.1179 | No | ns | 0.7749 |
| 0.01 mg/kg CNO (i.p.) KI vs. 0.01 mg/kg CNO (tooth) KI | 0.03 | -0.04627 to 0.1063 | No | ns | 0.9163 |
| 0.01 mg/kg CNO (i.p.) KI vs. Baseline WT | -0.08125 | -0.1575 to -0.004985 | Yes | * | 0.0291 |
| 0.01 mg/kg CNO (i.p.) KI vs. 0.01 mg/kg CNO (i.p.) WT | -0.075 | -0.1513 to 0.001265 | No | ns | 0.0571 |
| 0.01 mg/kg CNO (i.p.) KI vs. 0.1 mg/kg CNO (i.p.) WT | -0.065 | -0.1413 to 0.01127 | No | ns | 0.1492 |
| 0.01 mg/kg CNO (i.p.) KI vs. 0.01 mg/kg CNO (tooth) WT | -0.065 | -0.1413 to 0.01127 | No | ns | 0.1492 |
| 0.1 mg/kg CNO (i.p.) KI vs. 0.01 mg/kg CNO (tooth) KI | -0.008929 | -0.08787 to 0.07001 | No | ns | >0.9999 |
| 0.1 mg/kg CNO (i.p.) KI vs. Baseline WT | -0.1202 | -0.1991 to -0.04124 | Yes | *** | 0.0003 |
| 0.1 mg/kg CNO (i.p.) KI vs. 0.01 mg/kg CNO (i.p.) WT | -0.1139 | -0.1929 to -0.03499 | Yes | *** | 0.0008 |
| 0.1 mg/kg CNO (i.p.) KI vs. 0.1 mg/kg CNO (i.p.) WT | -0.1039 | -0.1829 to -0.02499 | Yes | ** | 0.0028 |
| 0.1 mg/kg CNO (i.p.) KI vs. 0.01 mg/kg CNO (tooth) WT | -0.1039 | -0.1829 to -0.02499 | Yes | ** | 0.0028 |
| 0.01 mg/kg CNO (tooth) KI vs. Baseline WT | -0.1113 | -0.1875 to -0.03498 | Yes | *** | 0.0006 |
| 0.01 mg/kg CNO (tooth) KI vs. 0.01 mg/kg CNO (i.p.) WT | -0.105 | -0.1813 to -0.02873 | Yes | ** | 0.0015 |
| 0.01 mg/kg CNO (tooth) KI vs. 0.1 mg/kg CNO (i.p.) WT | -0.095 | -0.1713 to -0.01873 | Yes | ** | 0.0056 |
| 0.01 mg/kg CNO (tooth) KI vs. 0.01 mg/kg CNO (tooth) WT | -0.095 | -0.1713 to -0.01873 | Yes | ** | 0.0056 |
| Baseline WT vs. 0.01 mg/kg CNO (i.p.) WT | 0.00625 | -0.07002 to 0.08252 | No | ns | >0.9999 |
| Baseline WT vs. 0.1 mg/kg CNO (i.p.) WT | 0.01625 | -0.06002 to 0.09252 | No | ns | 0.9974 |
| Baseline WT vs. 0.01 mg/kg CNO (tooth) WT | 0.01625 | -0.06002 to 0.09252 | No | ns | 0.9974 |
| 0.01 mg/kg CNO (i.p.) WT vs. 0.1 mg/kg CNO (i.p.) WT | 0.01 | -0.06627 to 0.08627 | No | ns | 0.9999 |
| 0.01 mg/kg CNO (i.p.) WT vs. 0.01 mg/kg CNO (tooth) WT | 0.01 | -0.06627 to 0.08627 | No | ns | 0.9999 |
| 0.1 mg/kg CNO (i.p.) WT vs. 0.01 mg/kg CNO (tooth) WT | 0 | -0.07627 to 0.07627 | No | ns | >0.9999 |

### SUPPLEMENTAL REFERENCES:


1. Madisen, L. *et al.* Transgenic mice for intersectional targeting of neural sensors and effectors with high specificity and performance. *Neuron* **85**, 942–958 (2015).

2. von Buchholtz, L. J. *et al.* Decoding Cellular Mechanisms for Mechanosensory Discrimination. *Neuron* **109**, 285-298.e5 (2021).

3. Stirling, C. L. *et al.* Nociceptor-specific gene deletion using heterozygous NaV1.8-Cre recombinase mice. *PAIN* **113**, 27 (2005).

4. Daigle, T. L. *et al.* A Suite of Transgenic Driver and Reporter Mouse Lines with Enhanced Brain-Cell-Type Targeting and Functionality. *Cell* **174**, 465-480.e22 (2018).

5. Servin-Vences, M. R. *et al.* PIEZO2 in somatosensory neurons controls gastrointestinal transit. *Cell* **186**, 3386-3399.e15 (2023).

6. Emrick, J. J., von Buchholtz, L. J. & Ryba, N. J. P. Transcriptomic Classification of Neurons Innervating Teeth. *J Dent Res* **99**, 1478–1485 (2020).

7. von Buchholtz, L. J., Lam, R. M., Emrick, J. J., Chesler, A. T. & Ryba, N. J. P. Assigning transcriptomic class in the trigeminal ganglion using multiplex in situ hybridization and machine learning. *Pain* **161**, 2212–2224 (2020).

8. Ghitani, N. *et al.* Specialized Mechanosensory Nociceptors Mediating Rapid Responses to Hair Pull. *Neuron* **95**, 944-954.e4 (2017).

9. Choi, H. M. T. *et al.* Third-generation in situ hybridization chain reaction: multiplexed, quantitative, sensitive, versatile, robust. *Development* **145**, dev165753 (2018).

10. Stringer, C., Wang, T., Michaelos, M. & Pachitariu, M. Cellpose: a generalist algorithm for cellular segmentation. *Nat Methods* **18**, 100–106 (2021).

11. Pachitariu, M. & Stringer, C. Cellpose 2.0: how to train your own model. *Nat Methods* **19**, 1634–1641 (2022).

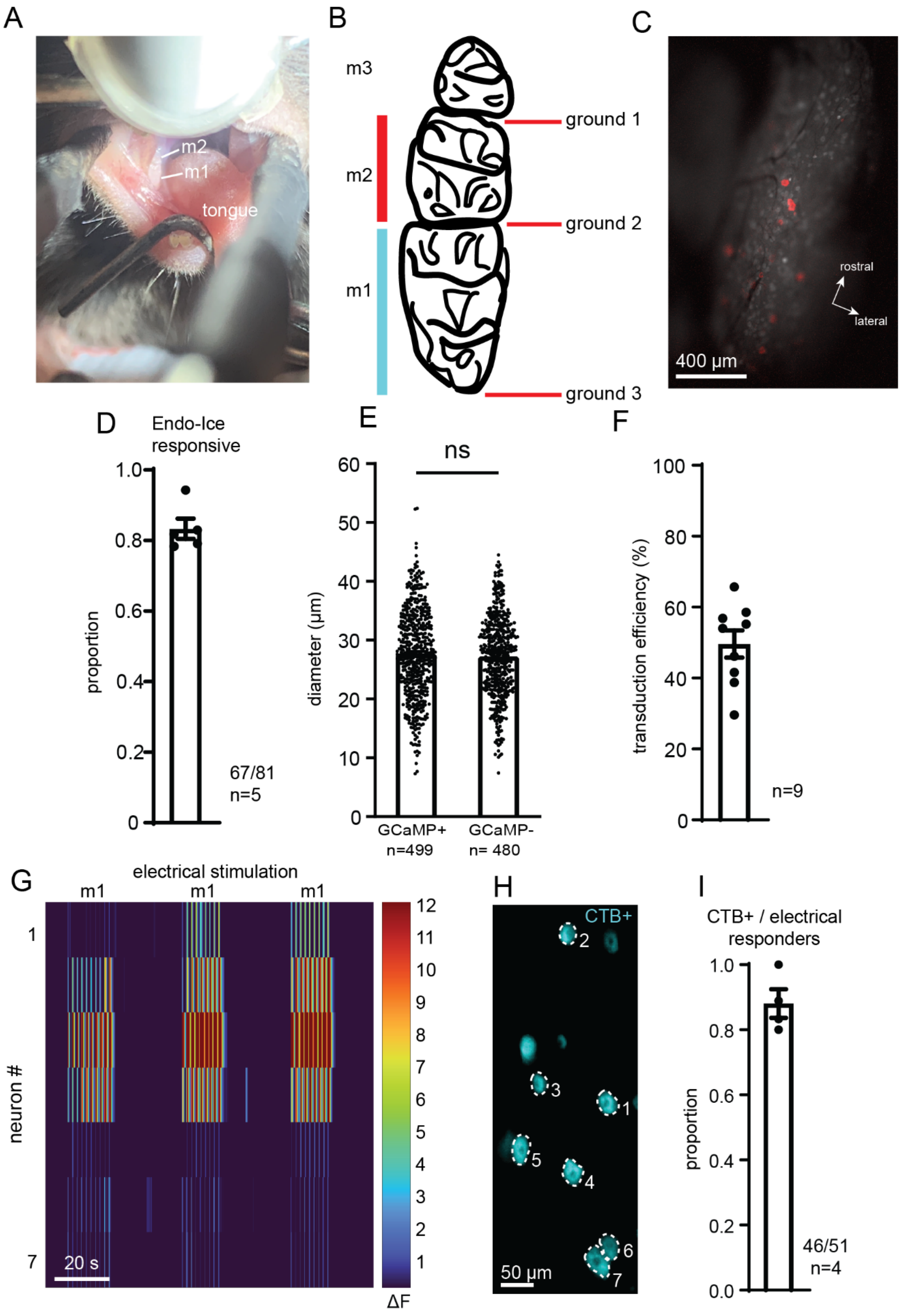
A
m2
m1
tongue
B
m3
m2
m1
ground 1
ground 2
ground 3
C
rostral
lateral
400 µm
D
Endo-Ice
responsive
proportion
1.0
0.8
0.6
0.4
0.2
0
67/81
n=5
E
ns
diameter (µm)
60
50
40
30
20
10
0
GCaMP+
n=499
GCaMP-
n= 480
F
transduction efficiency (%)
100
80
60
40
20
0
n=9
G
electrical stimulation
m1
m1
m1
neuron #
1
7
20 s
12
11
10
9
8
7
6
5
4
3
2
1
ΔF
H
CTB+
2
3
1
5
4
6
7
50 µm
I
CTB+ / electrical
responders
proportion
1.0
0.8
0.6
0.4
0.2
0.0
46/51
n=4

**Figure S1. Additional data on functional imaging of intradental neurons. Related to Figure 1.**

**(A)** Snapshot image showing the oral cavity field of view as viewed by operator in a head fixed mouse during in vivo imaging. Molar 1 (m1), molar 2 (m2), and the protruded tongue are annotated.

**(B)** Illustrative schematic showing locations of a fine wire cathode (-) placement during successive rounds of molar voltage stimulation. To identify intradental trigeminal neurons innervating a single molar, a fine wire anode (+) was held on the occlusal surface of the molar, while the cathode was positioned in the lingual gingiva. Delivery of voltage pulses to a single molar was repeated for a total of 3 times while varying the location of the cathode to each of the three ground positions shown.

**(C)** Epifluorescent image of a trigeminal ganglion in a AAV9-Cre injected Ai95D mouse demonstrating the localization of neurons responding to voltage pulses (red). In vivo visualization of the trigeminal ganglion surface is achieved via a cranial window preparation and a modified stereotax to fix the head (see Methods). Trigeminal neurons responding to molar stimulation are located in the caudal, mandibular division of the ganglion. Scale bar: 500μm.

**(D)** Bar graph depicting the proportion of intradental neurons that respond to Endo-Ice stimulation. Plotted individual data points represent the proportion of cold responsive/intradental neurons identified in each experiment. Bar shows the mean, and error bars indicate the SEM. n = 5 mice, comprising 81 total cells.

**(E)** Bar graph showing the mean diameter of GCaMP+ versus GCaMP- trigeminal neurons from adult Ai95(RCL-GCaMP6f)-D (Ai95D) mice injected postnatally (P0-P3) with AAV9-Cre.

The expression of GCaMP was amplified with ISH and quantified in nonsequential sections from 6 TG taken from 3 mice. Plotted individual data points represent single somal diameters. Bar shows the mean value, and error bars indicate the SEM. No significant differences were observed with $p>0.05$ using One-way ANOVA with Tukey's correction.

**(F)** Bar graph showing the transduction efficiency of GCaMP from adult Ai95(RCL-GCaMP6f)-D (Ai95D) mice injected postnatally (P0-P3) with AAV9-Cre. The expression of GCaMP was examined by ISH. Plotted individual data points represent single somal diameters. Bar shows the mean value, and error bars indicate the SEM. Quantifications are from 9 TG sections taken from 3 mice.

**(G-I)** Retrograde labeling marks intradental neurons that are activated by electrical stimulation of the molars. Retrograde tracer CTB-AF647 was applied to small cavitations in a single mandibular molar (m1) 16 hr prior to functional imaging in a AAV9-Cre injected Ai95D mouse. (G) Example heatmap showing calcium responses of CTB+ neurons to voltage pulses. (H) Epifluorescent image of corresponding CTB-647 retrograde-labeled intradental neurons shown in G. Scale bar, 50μm. (I) Bar graph showing the proportion of TG neurons responding to ≥2 voltage pulses that are CTB+. Bar shows the mean and error bars indicate the SEM. n = 4 mice. CTB, cholera toxin B-subunit.

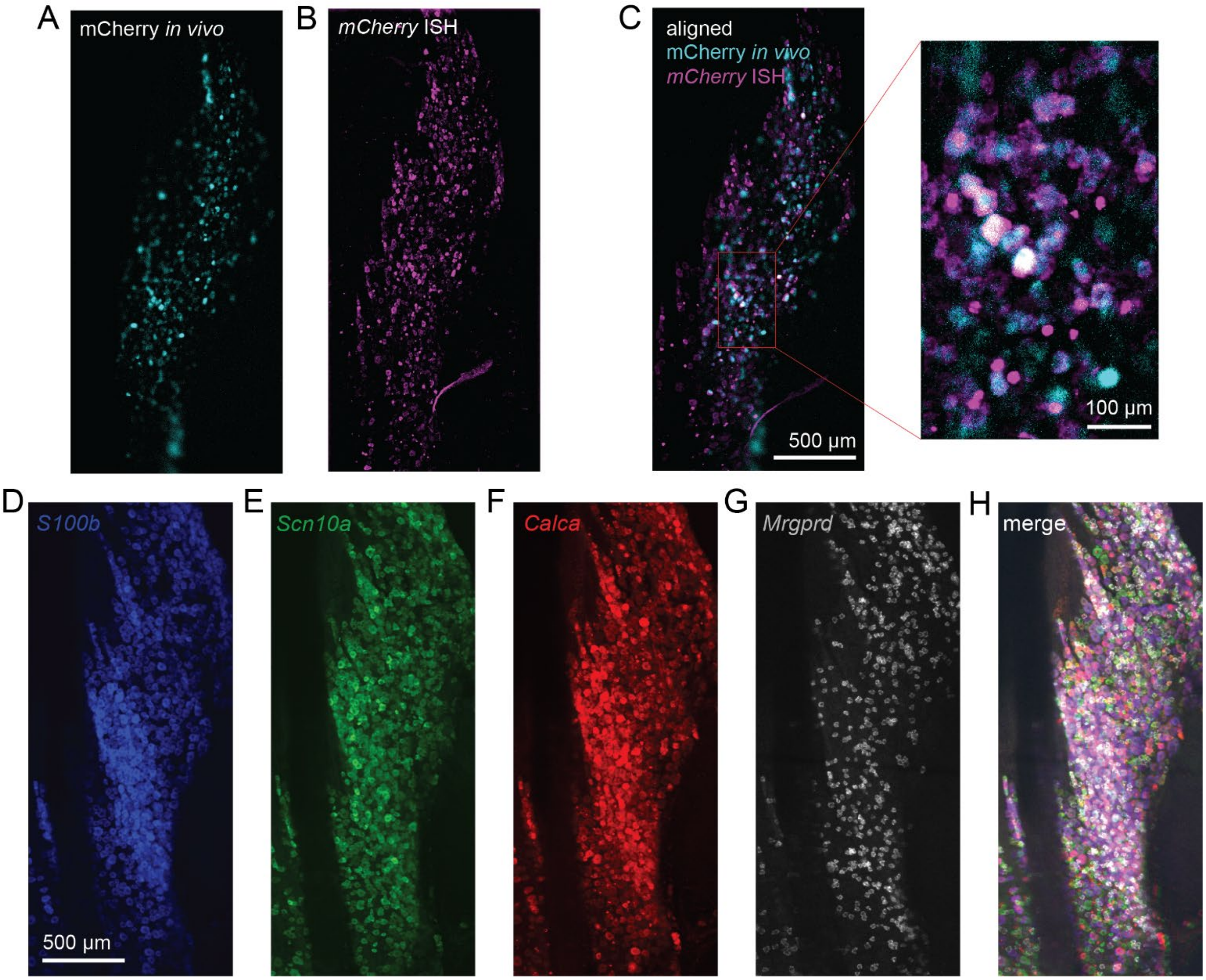


**Figure S2. Molecular characterization of functionally responding intradental neurons using post hoc whole mount in situ hybridization (ISH). Related to Figure 1.**

**(A-H)** Example images of alignment of functional images with ISH images (A) of the dorsal view of the mCherry-positive neurons in the TG (left panel, cyan); (B) image of the dorsal view of the excised ganglion after whole-mount ISH against mCherry(magenta); (C) images were aligned using a combination of Demon's algorithm and Bigwarp (see Methods) for image registration via MatLab (right panel). Inset of the field demonstrates overlap (white). TGs were from Ai95(RCL-GCaMP6f)-D (Ai95D) mice injected postnatally (P0-P3) with AAV9-hSyn1-mCherry-2A-iCre, enabling mCherry expression to serve as guideposts to align functional imaging with ISH images. Scale bar, 500µm. (D-H) Example images showing multiplexed whole-mount ISH staining following alignment to in vivo GCaMP6f fluorescence. Probes: s100 calcium-binding protein B (*S100b*, blue), alpha-Nav1.8 (*Scn10a*, green), calcitonin gene-related peptide (CGRP; *Calca*, red), mas-related G protein-coupled receptor member D (*Mrgprd*, white). Scale bar, 500µm (left).

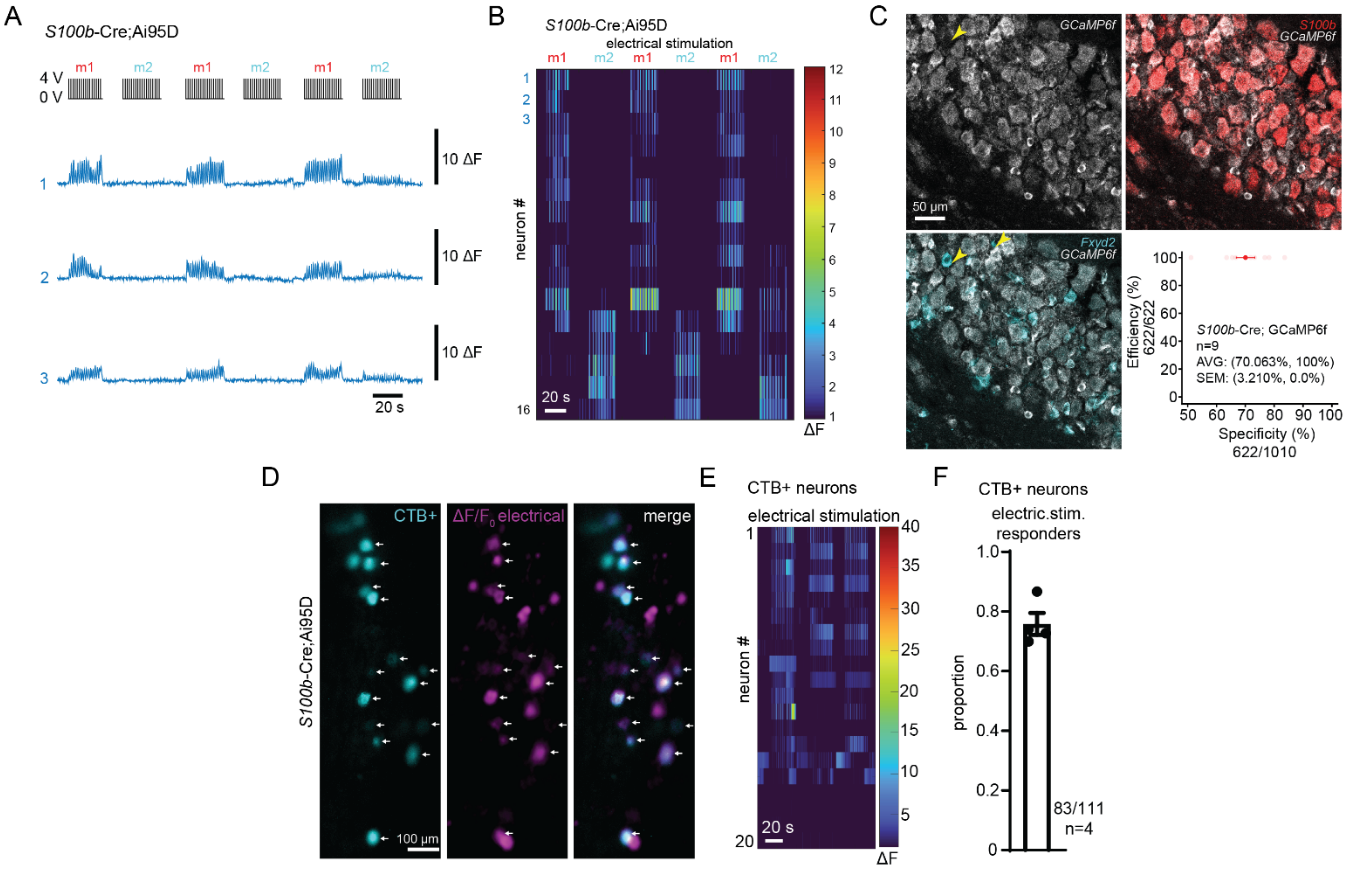


**Figure S3. Validation and functional imaging of *S100b*-Cre driver line. Related to Figure 1.**

**(A-B)** Example traces (A) and related heatmap (B) showing electrical responses in *S100b*-Cre; Ai95D transgenic mice. Bars to the right of traces indicate 10 ΔF.

(**C**) Validation of the *S100b*-Cre driver line. Representative ISH images for a single section of the trigeminal ganglion taken from *S100b*-Cre; Ai95D mice. Probes (left to right): *GCaMP6f*, s100 calcium-binding protein B (*S100b*), FXYD domain-containing ion transport regulator 2 (*Fxyd2*). Yellow arrows indicate *Fxyd2*(+)/*GCaMP6f*(-)/*S100b*(-) cells (left and right panels). Scale bar: 50μm. Line graph depicts summary of the specificity and efficiency of *GCaMP6f* expression in our transgenic *S100b*-Cre; Ai95D mice. Specificity refers to the percentage of *GCaMP6f*+ cells that are S100b+. Efficiency refers to the percentage of *S100b*+ cells that express *GCaMP6f*+. n = 9 mice.

**(D-F)** Data obtained from *S100b*-Cre; Ai95D mice. Retrograde labeling marks intradental neurons that are activated by electrical stimulation of the molars. Retrograde tracer CTB-AF555 was applied to small cavitations in mandibular molars (m1/m2) 16 hr prior to functional imaging. (D) Sample images depicting overlap of CTB labeling with $Ca^{2+}$ response (ΔF) of neurons responding to applied voltage pulses. Scale bar, 50μm. Left panel: CTB labeling; Middle panel: single frame (ΔF) from recording during response to voltage pulse; Right panel: Merge. (E) Example heatmap showing responses of combined m1/m2 CTB+ neurons to voltage pulses. (F) Bar graph showing the proportion of CTB+ TG neurons that respond to ≥2 voltage pulses. Plotted individual data points represent the calculated proportion of CTB+ cells that respond to ≥2 voltage pulses for each ganglion. Bar shows the mean and error bars indicate the SEM. n = 4 mice. CTB, cholera toxin B-subunit.

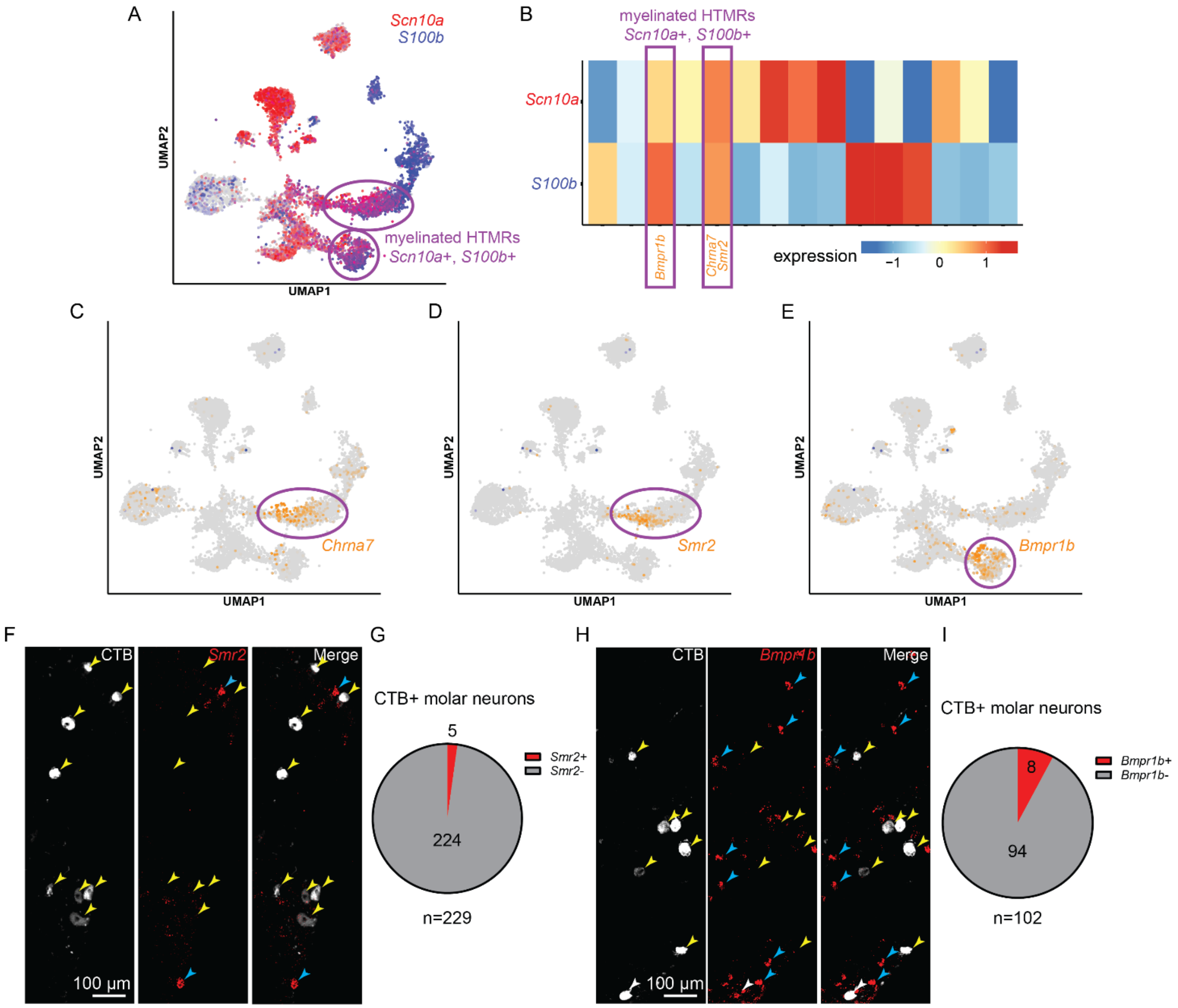


**Figure S4. Additional data on markers to differentiate HTMRs and transcriptional expression of intradental neurons. Related to Figure 1.**

**(A-E)** Gene expression data from integrated datasets and plots generated from painseq.shinyapps.io (Bhuiyan, A. S., Xu M., et al. doi: 10.1126/sciadv.adj9173).

**(A-B)** UMAP projection and heatmap representing gene expression patterns of *Scn10a* (red) and *S100b* (blue). **(A)** Dots represent cells/nuclei. Clusters represent distinct classes of trigeminal sensory neurons from the atlas. Co-expression of *Scn10a* and *S100b* (purple) is found primarily within two clusters that represent myelinated HTMRs (purple ovals). **(B)** Heatmap representing dataset from **(A)** demonstrating Scn10a and S100b normalized expression across distinct classes of trigeminal sensory neurons from the atlas. Orange and red indicate normalized expression >0. Two classes feature co-expression of Scn10a and S100b >0 (purple rectangles) and represent myelinated HTMRs that are also defined by enriched expression of *Bmpr1b*, *Chrna7*, *Smr2*.

**(C-E)** UMAP projection representing gene expression patterns of **(C)** *Chrna7***, (D)** *Smr2***,** or **(E)** *Bmpr1b*. Co-expression of *Scn10a* and *S100b* from **(A)** is found primarily within two clusters that represent myelinated HTMRs (purple ovals).

(**F-G**) Representative image from retrograde-labeling of intradental neurons using CTB-AF647 tracer followed by ISH. Retrograde tracer CTB-AF647 was applied to small cavitations in maxillary and mandibular molars (m1/m2) 16 hr prior to tissue harvest. The expression of the candidate marker gene *Smr2* was examined by ISH. (F) Left panel: CTB labeling; Middle panel: *Smr2* expression; Right panel: Merge. Scale bar: 100 µm. Yellow arrow heads indicate CTB-AF647+ intradental neurons. Blue arrowheads indicate cells that are only positive for *Smr2*. (G) Pie chart representing counts of CTB-labeled intradental neurons that were positive or negative for *Smr2*. n=3 mice, comprising 229 retrograde-labeled intradental neurons.

(**H-I**) Representative image from retrograde-labeling of intradental neurons using CTB-AF647 tracer followed by ISH. Retrograde tracer CTB-AF647 was applied to small cavitations in maxillary and mandibular molars (m1/m2) 16 hr prior to tissue harvest. The expression of the candidate marker gene *Bmpr1b* was examined by ISH. (H) Left panel: CTB labeling; Middle panel: *Bmpr1b* expression; Right panel: Merge. Scale bar: 100 µm. Yellow arrow heads indicate CTB-AF647+ intradental neurons. Blue arrowheads indicate cells that are only positive for *Bmpr1b*. White arrow head indicates retrograde-labeled intradental neuron that was co-positive for *Bmpr1b* (I) Pie chart representing counts of CTB-labeled intradental neurons that were positive or negative for *Bmpr1b*. n=3 mice, comprising 102 retrograde-labeled intradental neurons.

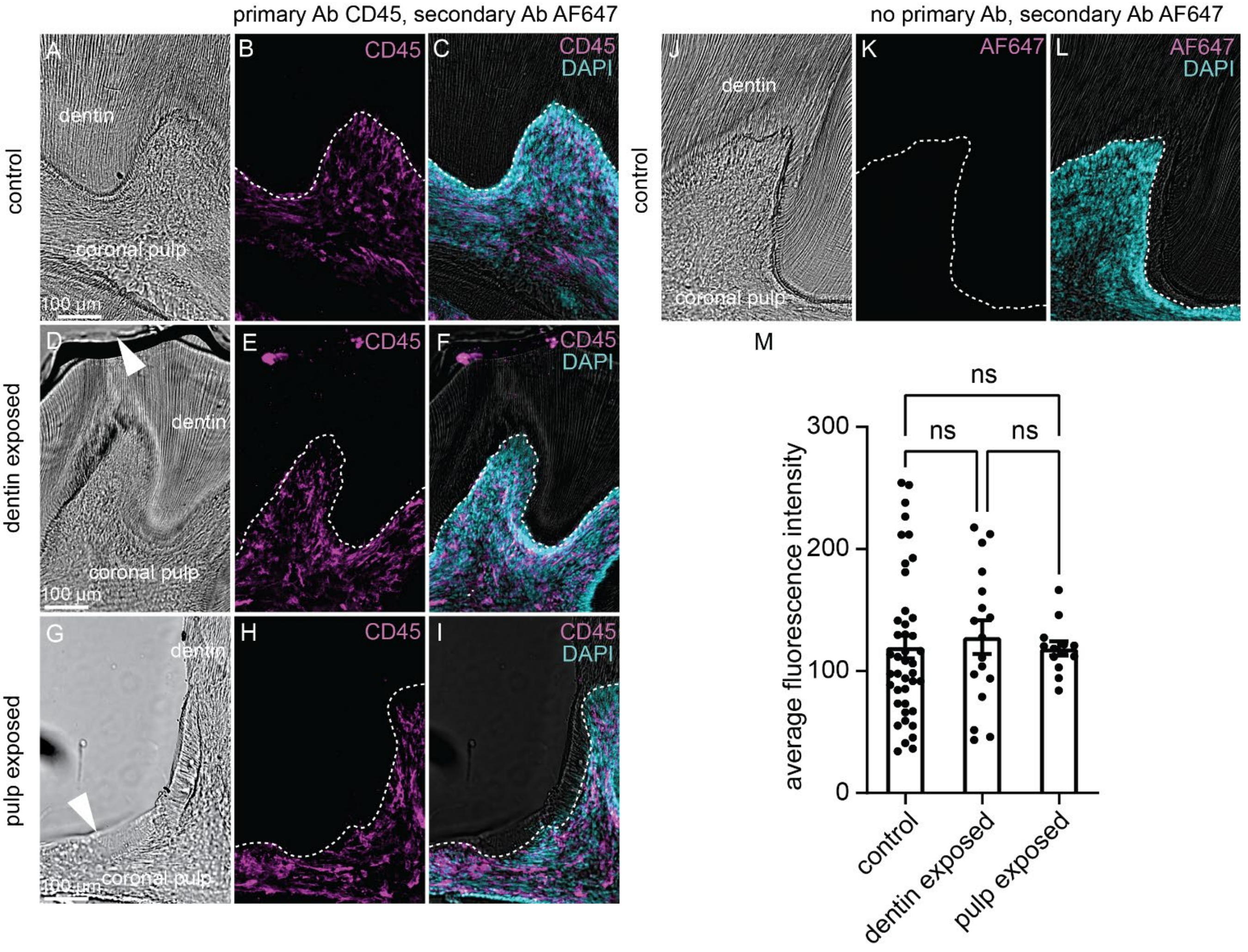


**Figure S5. Acute dentin/pulp exposure does not alter pulpal immune cells. Related to Figure 2.**

**(A-I)** Representative images of staining for immunoreactivity to CD45 in control, dentin exposed, or pulp exposed molars. Enamel, a fully mineralized structure, has been removed completely by demineralization. (A) Brightfield image of control m1 molar cross section, highlighting anatomy of the coronal pulp versus dentin boundary. (B) Immunostaining for CD45. (C) Merged image of (B) overlaid with DAPI staining labeling nuclei of pulp cells and brightfield image. Dashed lines in B-C indicate the pulp-dentin border. This experiment was repeated in n = 3 mice. Scale bar: 100μm (D) Brightfield image of m1 molar cross section following dentin exposure, highlighting anatomy of the coronal pulp versus dentin boundary. White arrowhead indicates depth of dentin exposure. (E) Immunostaining for CD45. (F) Merged image of (E) overlaid with DAPI staining labeling nuclei of pulp cells and brightfield image. Dashed lines in E-F indicate the pulp-dentin border. This experiment was repeated in n = 3 mice. Scale bar: 100μm. (G) Brightfield image of m1 molar cross section following pulp exposure. White arrowhead indicates exposed pulp. (H) Immunostaining for CD45. (I) Merged image of (H) overlaid with DAPI staining labeling nuclei of pulp cells and brightfield image. Dashed lines in H-I indicate the pulp-dentin border. This experiment was repeated in n = 3 mice. Scale bar: 100μm

(**J-L**) Representative image of immunostaining omitting primary antibody for CD45. (J) Brightfield image of m1 molar cross section. (K) Immunostaining shows no AF647+ signal in the absence of the primary antibody. (L) Merged image of (K) overlaid with DAPI-stained cell nuclei in the pulp and brightfield image. Dashed line in K-L indicates pulp-dentin border. This experiment was repeated in n = 3 mice. Scale bar: 100μm.

(**M**) Bar graph depicting the quantified mean fluorescent intensity of CD45 within the pulp across all conditions. Plotted individual data points represent the average intensity for individual tooth pulps. Bar shows the mean, and error bars indicate the SEM. n = 3 mice. No significant differences were observed with $p>0.05$ using One-way ANOVA with Tukey's correction.

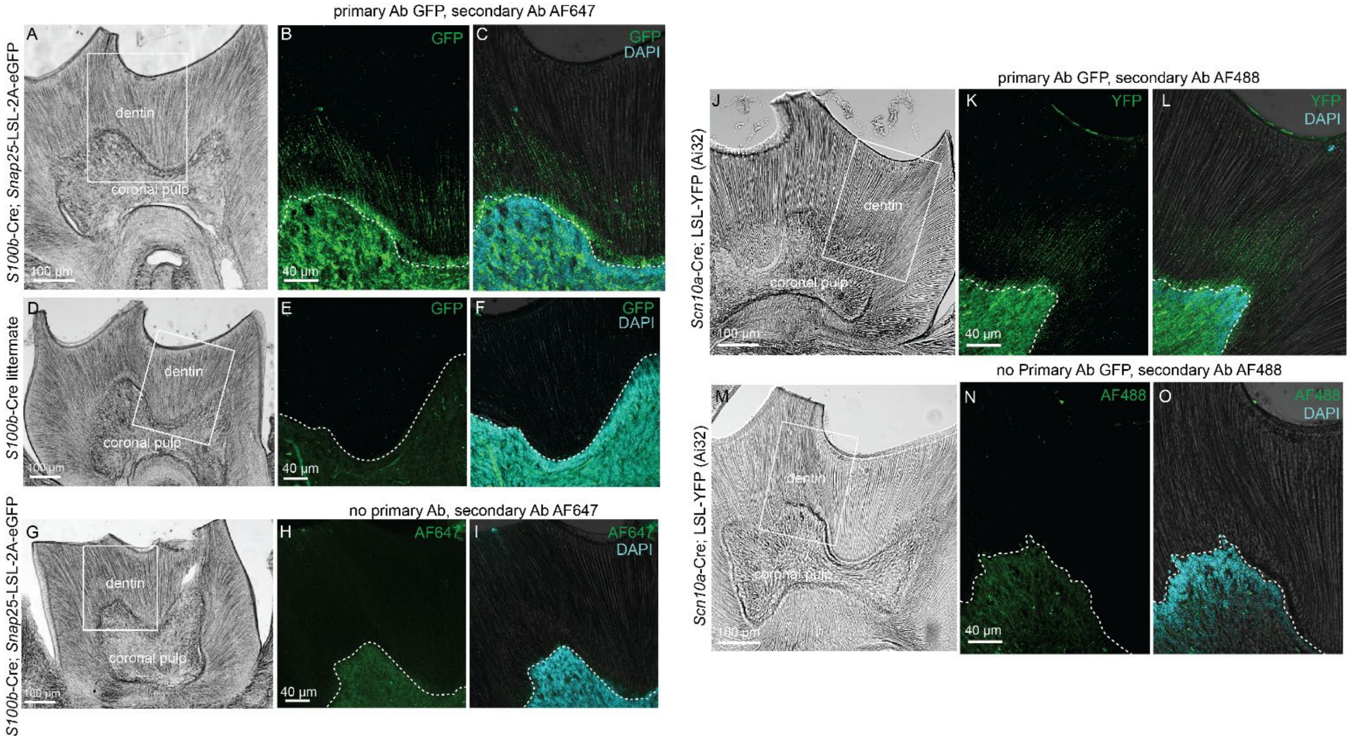


**Figure S6. *S100b*-Cre and *Scn10a*-Cre driver lines label terminal endings in pulp and inner dentin. Related to Figure 2.**

**(A-C)** Representative images of immunostained neuronal terminals in demineralized molars from *S100b*-Cre; *Snap25*-LSL-2A-eGFP mice. Enamel, a fully mineralized structure, has been removed completely by demineralization. (A) Brightfield image of molar cross section, highlighting anatomy of the coronal pulp versus dentin boundary. (B) Magnified image of inset from (A). Immunostaining for GFP is localized to *S100b*-positive terminal endings that radiate from the pulp and extend into the dentin. (C) Merged image of (B) overlaid with DAPI staining labeling nuclei of pulp cells and brightfield image. Dashed lines in B-C indicate the pulp-dentin border. This experiment was repeated in n = 3 mice. Scale bars: 100μm (A), 40μm (B,C).

**(D-F)** Representative image of *S100b*-Cre; *Snap25*-LSL-2A-eGFP(-) littermate controls showing no GFP+ signal. Enamel, a fully mineralized structure, has been removed completely by demineralization. (D) Brightfield image of molar cross section. (E-F) Magnified image of inset from (D). (F) Merged image of (E) overlaid with DAPI staining labeling nuclei of pulp cells and brightfield image showing immunostaining for GFP is absent. Dashed line in E-F indicates pulp-dentin border. This experiment was repeated in n = 3 mice. Scale bars: 100μm (D), 40μm (E, F)

**(G-I)** Representative image of immunostaining omitting primary antibody for *S100b*-Cre; *Snap25*-LSL-2A-eGFP mice. Enamel, a fully mineralized structure, has been removed completely by demineralization. (G) Brightfield image of molar cross section. (H) Magnified image of inset from (G). (I) Immunostaining shows no GFP+ signal, overlaid with DAPI-stained cell nuclei in the pulp and brightfield image. Dashed line in H-I indicates pulp-dentin border. This experiment was repeated in n = 3 mice. Scale bars: 100μm (G), 40μm (H, I)

**(J-L)** Representative images of immunostained neuronal terminals in demineralized molars from *Scn10a*-Cre; *Ai32* mice. Enamel, a fully mineralized structure, has been removed completely by demineralization. (J) Brightfield image of molar cross section, highlighting anatomy of the coronal pulp versus dentin boundary. (K) Magnified image of inset from (J). Immunostaining for YFP is localized to *Scn10a*-positive terminal endings that radiate from the pulp and extend into the dentin. (L) Merged image of (K) overlaid with DAPI staining labeling nuclei of pulp cells and brightfield image. Dashed lines in K-L indicate the pulp-dentin border. This experiment was repeated in n = 4 mice. Scale bars: 100μm (J), 40μm (K, L).

**(M-O)** Representative image of immunostaining omitting primary antibody for *Scn10a*-Cre; *Ai32* mice. Enamel, a fully mineralized structure, has been removed completely by demineralization. (M) Brightfield image of molar cross section. (N) Magnified image of inset from (M). (N) Immunostaining shows no YFP+ signal, overlaid with DAPI-stained cell nuclei in the pulp and brightfield image. Dashed line in N-O indicates pulp-dentin border. This experiment was repeated in n = 4 mice. Scale bars: 100μm (M), 40μm (N, O).

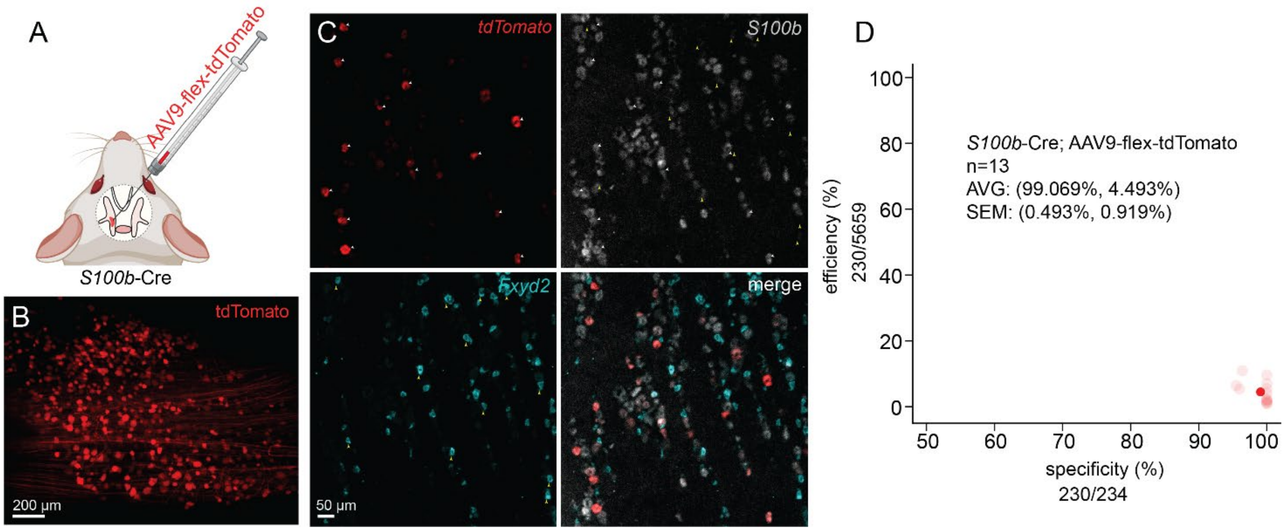

**Figure S7. Validation of the *S100b*-Cre driver line in conjunction with AAV9-flex-tdT direct trigeminal injections. Related to Figure 2.**

**(A)** Schematic of methodology to sparsely label trigeminal neurons and intradental neuron endings. AAV9-flex-tdTomato was orbitally injected into the trigeminal ganglion to induce tdTomato labeling in neurons of *S100b*-Cre mice.

**(B)** Example whole mount image of the TG validating approach shown in (A) depicting tdTomato+ positive somas in an *S100b*-Cre mouse following orbital delivery of AAV9-flex-tdT. Scale bar: 200 µm.

**(C)** Representative ISH images for a single section of the trigeminal ganglion taken from a S100b-Cre mouse following orbital injection of AAV9-flex-tdTomato. Probes (left to right): *tdTomato*, s100 calcium-binding protein B (*S100b*), FXYD domain-containing ion transport regulator 2 (*Fxyd2*). Merge: bottom right panel. Green arrow heads indicate particular *Scn10a*+ cells. White arrow heads highlight soma co-positive for *S100b* and *tdTomato*. Yellow arrowheads indicate cells that are only positive for *Fxyd2*. Scale bar: 50 µm.

**(D)** Summary of the specificity and efficiency of TG *tdTomato* expression following AAV9-flex-tdTomato orbital injection in *S100b*-Cre mice. Specificity refers to the percentage of *tdTomato*+ cells that are *S100b*+. Efficiency refers to the percentage of *S100b*+ cells that express *tdTomato*+. n = 13 mice.

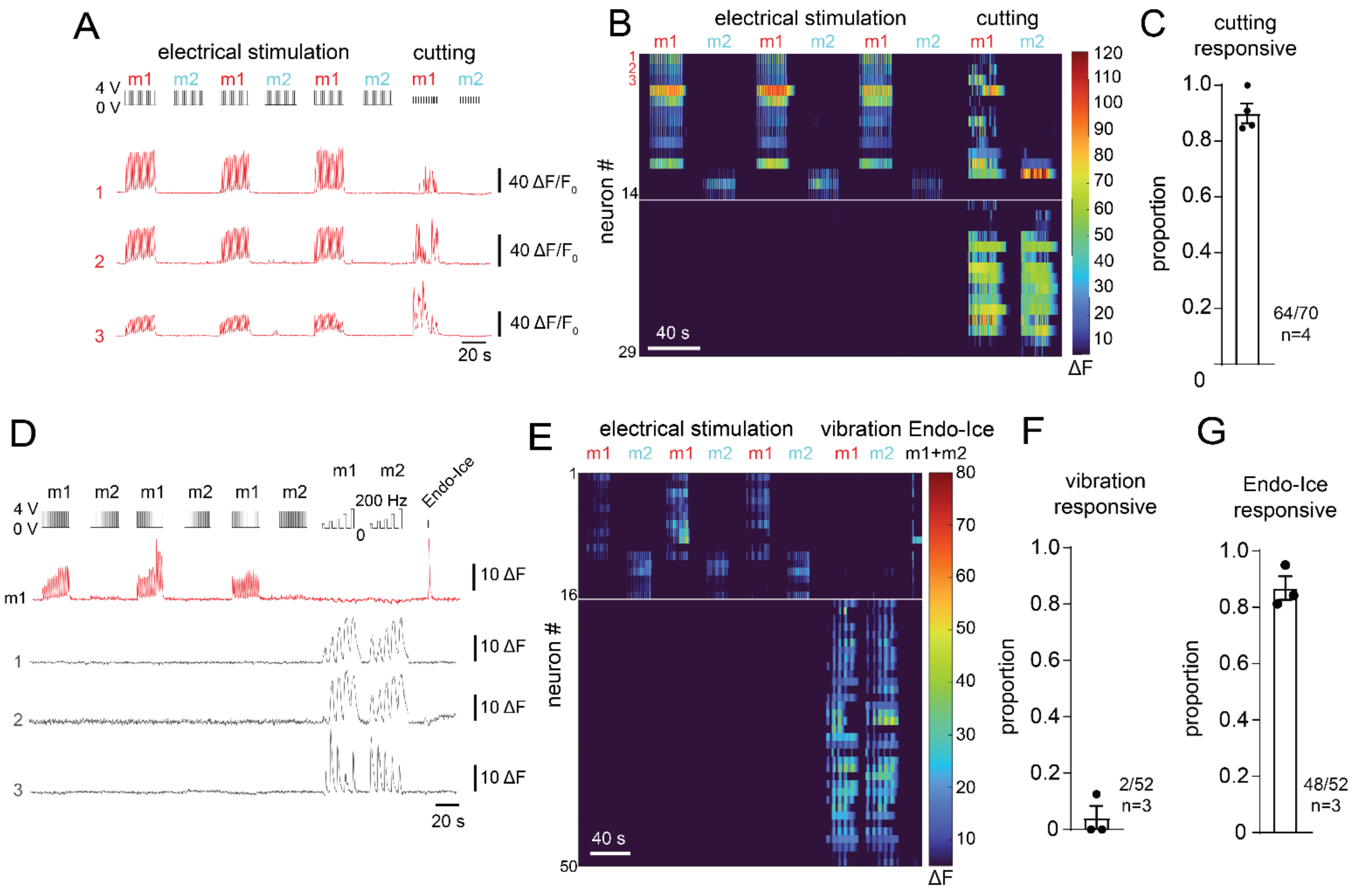


**Figure S8. Intradental neuron respond to enamel damage but not vibration of intact tooth. Related to Figure 3.**

**(A-C)** Intradental HTMRs encode cutting of tooth enamel. (A) Example traces showing intradental neuron response to enamel cutting applied to individual molars. Stimuli used are indicated above the traces. Bars to the right of traces indicate 40 ΔF. (B) Example heatmap containing traces shown in (A). (C) Bar graph showing proportion of cutting responsive intradental neurons identified by electrical stimulation. Plotted individual data points represent the proportion for each trial. Bar shows the mean and error bars indicate the SEM. n = 4 mice, comprising 70 total cells. Data for A-C were obtained from Ai95(RCL-GCaMP6f)-D (Ai95D) mice injected postnatally (P0) with AAV9-Cre.

**(D-G)** Intradental neurons do not respond to vibration applied to the intact tooth. (D) Example traces showing intradental neurons response to surface vibration of the intact tooth. Gray traces (cells 1-3) show that molar surface vibration induces transient responses in select non-intradental innervating neurons. Stimuli used are indicated above the traces. Bars to the right of traces indicate 10 ΔF. (E) Example heatmap related to traces shown in (H). Vibration responders were only observed in non-intradental innervating neurons (did not respond to electrical or cold stimulation of molars). Neurons were grouped as intradental neurons (cells 1-16) or non-intradental vibration responders (cells 17-50). Stimuli used are indicated above the heatmap. (F) Bar graph showing proportion of vibration responses/intradental neurons. Plotted individual data points represent the calculated proportion of intradental neurons that also respond to vibration. Bar shows the mean and error bars indicate the SEM. n = 3 mice, comprising 52 total intradental neurons. (G) Bar graph depicting the proportion of intradental neurons that respond to Endo-Ice stimulation. Plotted individual data points represent the proportion of cold responsive/intradental neurons identified in a given experiment. Bar shows the mean, and error bars indicate the SEM. n = 3 mice, comprising 52 total cells. Data were obtained from Ai95(RCL-GCaMP6f)-D (Ai95D) mice injected postnatally (P0-P3) with AAV-Cre.

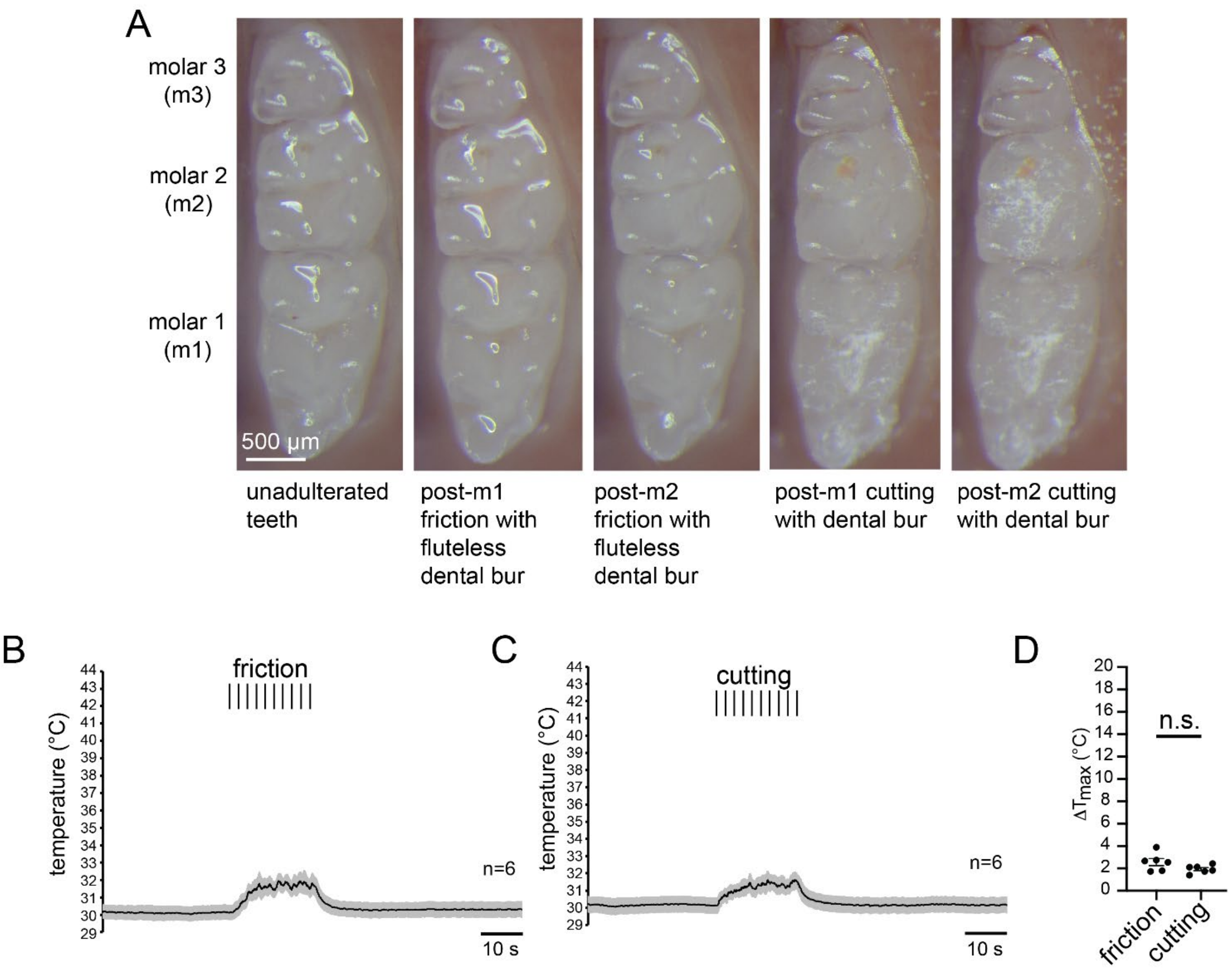


**Figure S9. Additional data on cutting and friction stimuli. Related to Figure 3.**

**(A)** Snapshot images showing occlusal surface structure of murine molars before and after friction and then cutting. Application of friction to m1 or m2 via a rounded "fluteless" carbide dental bur generates diminutive damage to the molar occlusal surface. Enamel cutting of m1 or m2 via a ¼ carbide dental bur produces shallow damage to the enamel occlusal surface as indicated by white flecks. Scale bar: 500 µm.

**(B-D)** Graphs depicting friction (B) and cutting (C) stimulation produce minimal temperature increases (<3ºC) in tooth pulp temperature as measured with a thermocouple implanted into the pulp bed while leaving the molar occlusal surface intact (see Methods for details). Traces show the average, shading indicates the SEM. n = 6 measurements per condition. (D) Bar graph showing maximal change in temperature in the tooth pulp in response to stimuli. Plotted individual data points represent the calculated ΔTmax (°C) for each measurement. Bar shows the mean and error bars indicate the SEM. n = 6 (friction), n = 6 (cutting). p>0.05 friction vs. drill, Paired t-test.

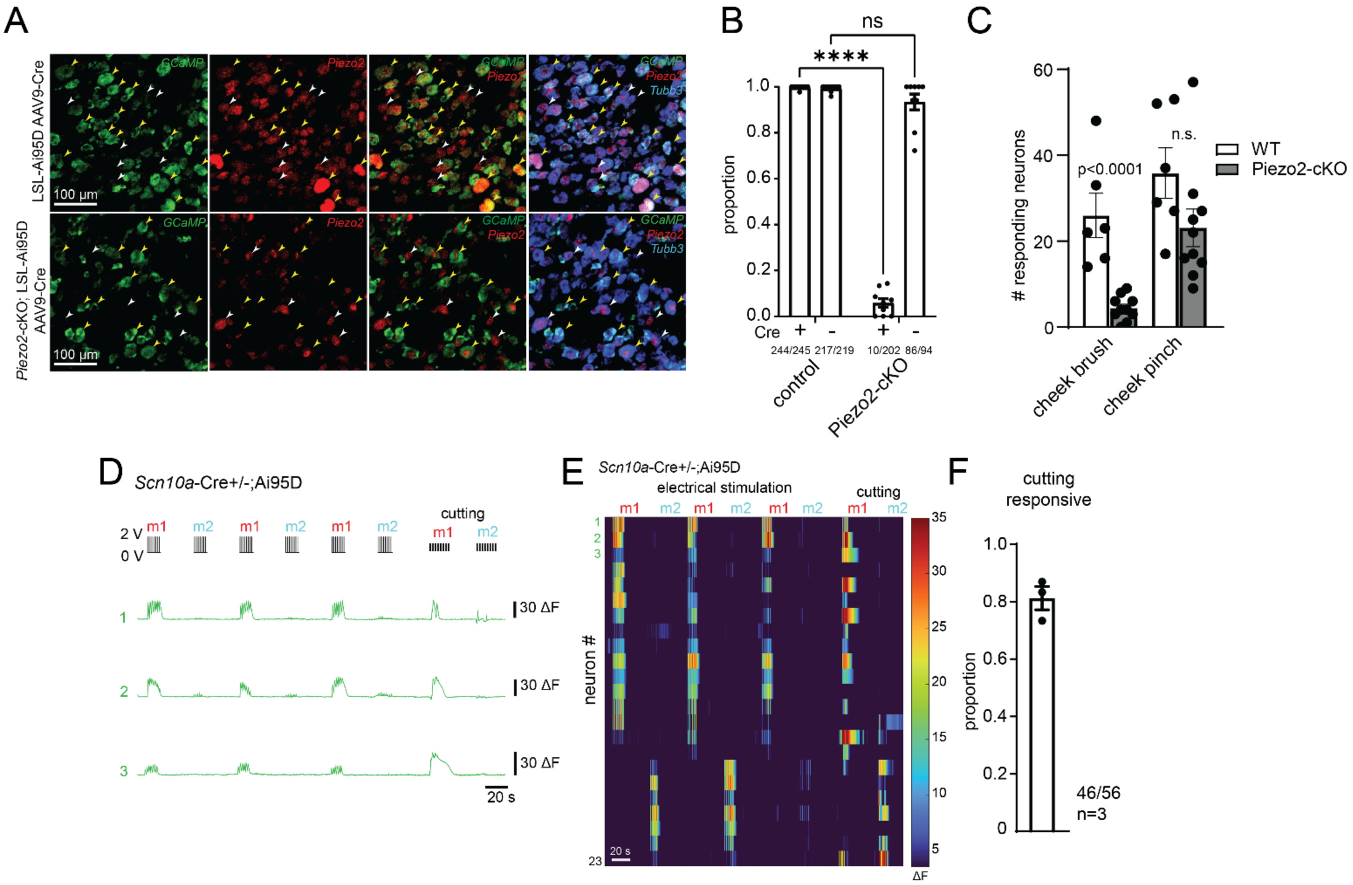


**Figure S10. Additional data related to *Piezo2*-cKO and *Scn10a*-Cre strains. Related to Figure 3.**

**(A)** Representative ISH images for a single section of the trigeminal ganglion taken from Ai95D (top) or *Piezo2*-cKO; Ai95D mice (bottom) injected postnatally (P0-P3) with AAV9-Cre. Probes (left to right): *Piezo2* (red), *GCaMP* (green), *Tubb3* (cyan). Yellow arrow heads indicate *GCaMP*+ neurons, white arrow heads indicate *GCaMP*- neurons. In *Piezo2*-cKO; Ai95D mice white arrow heads correspond to neurons with cytoplasmic *Piezo2* without cKO. Scale bar: 100 μm.

**(B)** Bar graph showing the proportion of diffuse cytoplasmic *Piezo2* in TG neurons in Ai95D (top) or *Piezo2*-cKO; Ai95D mice. Cre expression was inferred by expression of GCaMP ISH signal. Plotted individual data points represent proportions scored within each section. Bar shows the mean and error bars indicate the SEM. n = 3 mice per group, with data taken from at least 3 sections averaged per animal. ****$p<0.0001$, ns = $p>0.05$, Two-way ANOVA with Šídák's multiple comparisons test.

**(C)** Functional validation of AAV9 Cre mediated *Piezo2*-cKO. Bar graph showing in vivo functional imaging TG responses to cheek stimulation in WT versus *Piezo2*-cKO mice. Plotted individual data points represent responding neurons to either cheek brush or pinch. Bar shows the mean, and error bars indicate the SEM. n = 6 (WT), n = 6 (*Piezo2*-cKO). $p<0.0001$, n.s. = $p>0.05$, Paired t-test

**(D-F)** *Scn10a*-Cre heterozygote enables functional identification of intradental HTMRs that respond to cutting

(D) Example traces showing cutting responses of intradental HTMRs in *Scn10a*-Cre+/-;Ai95D mice. Stimuli used are indicated above the traces. Bars to the right of traces indicate 30 ΔF. (E) Example heatmap containing traces shown in (D). Stimuli used are indicated above the heatmap. (F) Bar graph showing proportion of cutting responsive intradental neurons. Plotted individual data points represent the proportion for each trial. Bar shows the mean and error bars indicate the SEM. n = 3 mice, comprising 56 total cells.

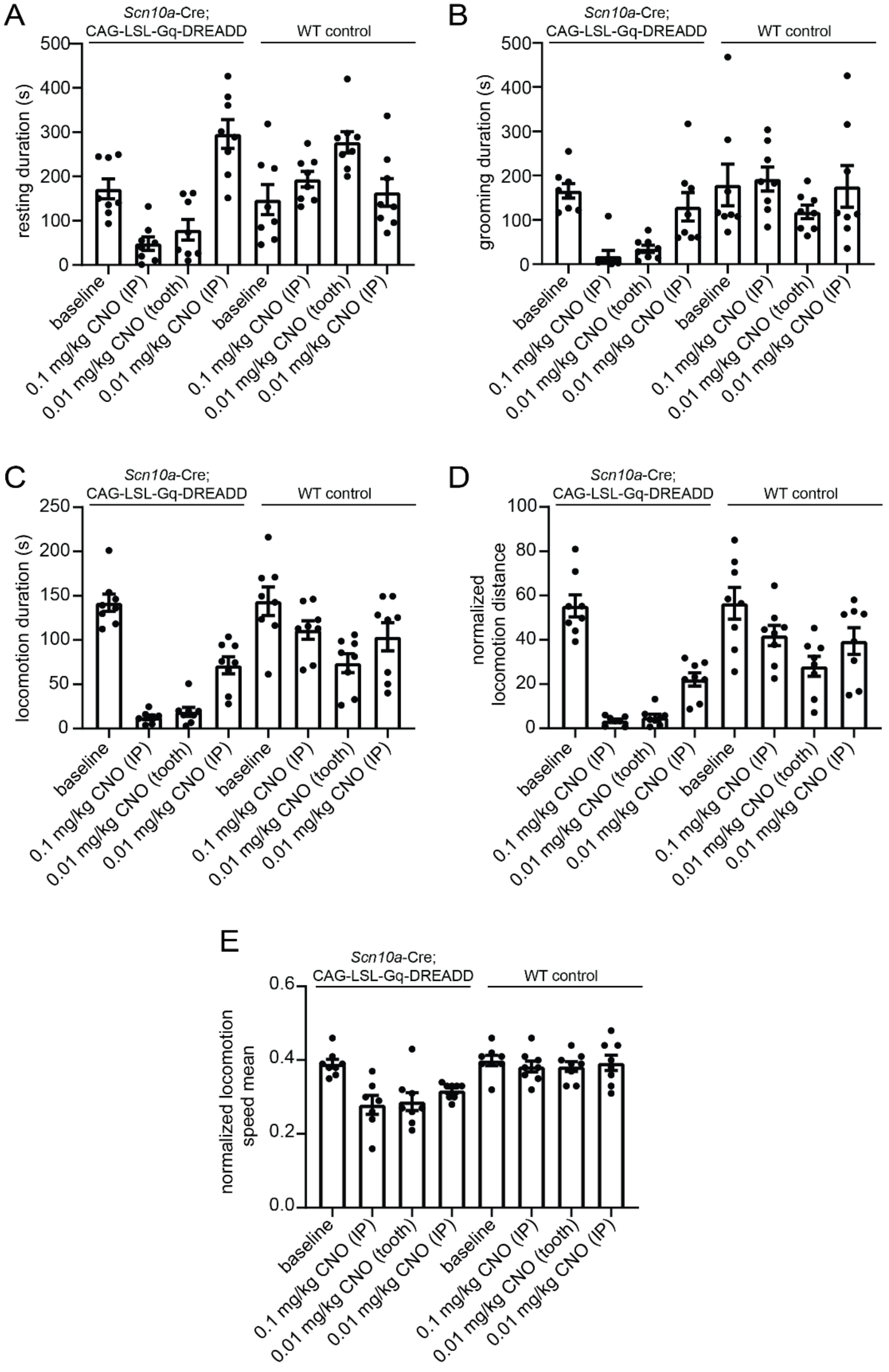


**Figure S11. Quantification of resting, grooming, and locomotion behavior from *Scn10a*-Cre;CAG-LSL-Gq-DREADD and WT control mice. Related to Figure 4.**

**(A)** Bar graph depicting resting duration across conditions. Plotted individual data points represent cumulative values for individual animals. Bar shows the mean, and error bars indicate the SEM (see Table 2 for statistics).

**(B)** Bar graph depicting grooming duration across conditions. Plotted individual data points represent cumulative values for individual animals. Bar shows the mean, and error bars indicate the SEM (see Table 3 for statistics).

**(C)** Bar graph depicting locomotion duration across conditions. Plotted individual data points represent cumulative values for individual animals. Bar shows the mean, and error bars indicate the SEM (see Table 4 for statistics).

**(D)** Bar graph depicting normalized locomotion distance across conditions. Plotted individual data points represent cumulative values for individual animals. Bar shows the mean, and error bars indicate the SEM (see Table 5 for statistics).

**(E)** Bar graph depicting normalized locomotion speed mean across conditions. Plotted individual data points represent the mean for individual animals. Bar shows the mean, and error bars indicate the SEM (see Table 6 for statistics).

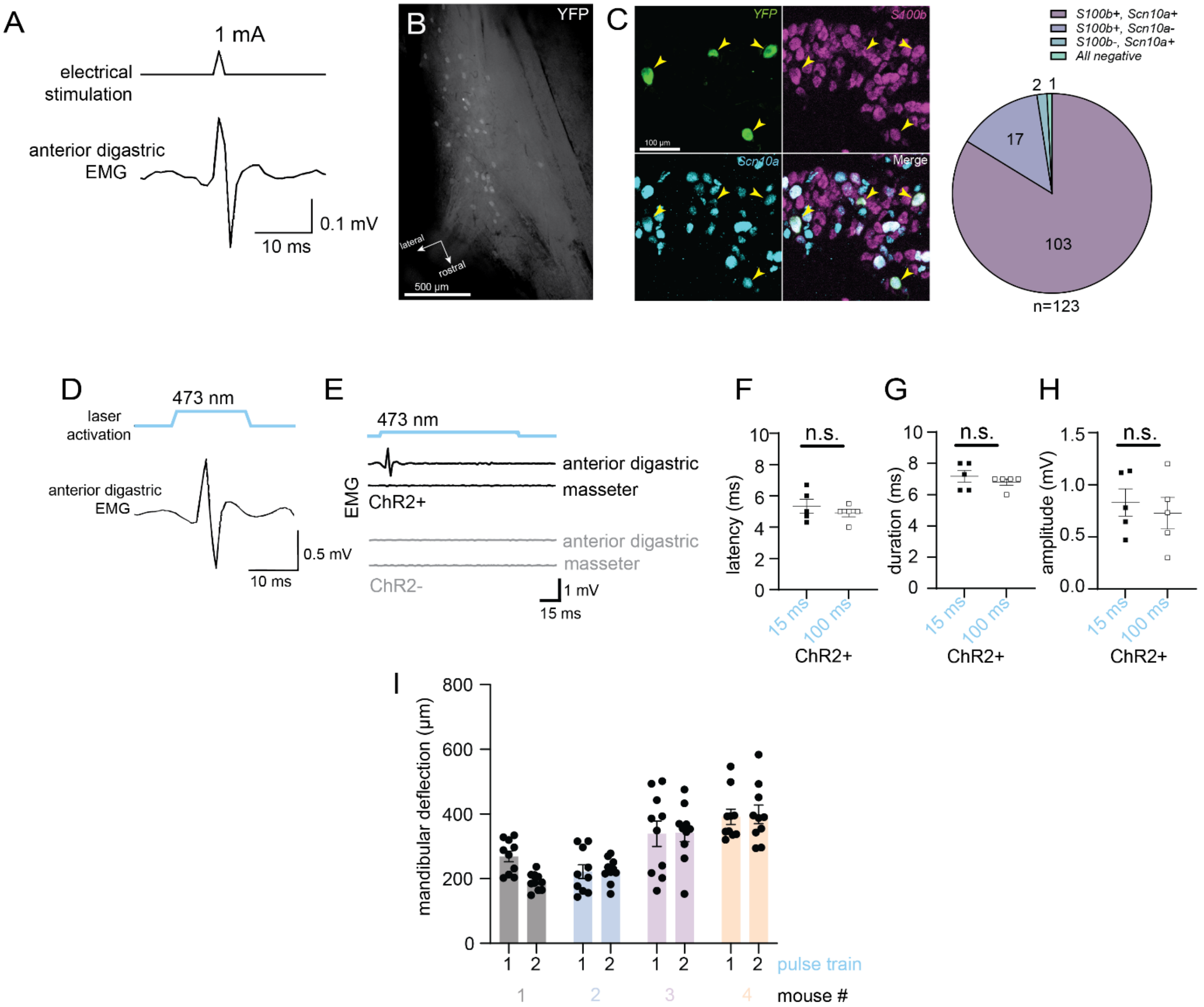


**Figure S12. Electromyography and jaw deflection elicited by activation of intradental neurons. Related to Figure 5.**

**(A)** Example trace of anterior digastric muscle EMG elicited by electrical stimulation of inferior alveolar nerve (1 mA, 0.1 ms, 1 Hz electrical stimulation). Experiment was repeated in n = 3 mice.

**(B)** Example image of the dorsal view of the excised ganglion showing ChR2-eYFP positive intradental neurons in the TG induced by AAV-Cre injection into the ipsilateral molars. Scale bar: 500 μm.

**(C)** Left: Representative ISH images for a single section of the trigeminal ganglion showing labeled YFP+ intradental neurons taken from an Ai32 (Cre-dependent ChR2-YFP) mouse receiving AAV-Cre tooth injections. Probes: *YFP* (top left), *Scn10a* (top right), *S100b* (bottom left), merge (bottom right). Yellow arrow heads indicate *YFP*+ intradental neurons. Scale bar: 100 μm. Right: Pie chart representing counts of *YFP*+ intradental neurons expressing *Scn10a* and/or *S100b*. n = 4 mice, comprising 123 total *YFP*+ neurons.

**(D)** Example trace of anterior digastric muscle EMG elicited by optogenetic activation of ipsilateral intradental neurons (15 ms, 1 Hz, 43 nm laser activation). Experiment was repeated in n = 3 mice.

**(E)** Optogenetic activation (100 ms duration) induces anterior digastric muscle activity when channelrhodopsin-2 is expressed in intradental neurons, with no activity observed in the masseter muscle. Control ChR2- group shows no changes in muscle activity for either the anterior digastric or masseter muscles. Experiment was repeated in n = 5 mice.

(**F-H**) Graph showing the (F) latency, (G) duration, and (H) amplitude of the digastric muscle activity elicited 15 ms or 100 ms optogenetic activation depicting average +/- SD for each condition. (F) Latency: 15 ms: 5.2 +/- 1.3 ms vs. 100 ms: 4.8 +/- 0.8 ms. (G) Duration: 15 ms: 6.6 +/- 0.6 ms vs. 100 ms: 6.7 +/- 0.6 ms. (H) Amplitude: 15 ms: 0.97 +/- 0.3 mV vs. 100 ms: 0.93 +/- 0.2 mV. For all conditions n = 5 mice and $p>0.05$, Unpaired t-Test.

**(I)** Bar graph depicting the amplitude of mandibular deflections in response to a blue-light pulse trains (10 pulses, 470nm, 1s), from data shown in Figure 4F, G. Plotted data points represent the measurements from individual mandibular deflections during pulse trains. Bar shows the mean and error bars indicate the SEM. n = 4 mice, comprising 160 measurements. $p>0.05$, Unpaired t-Test.

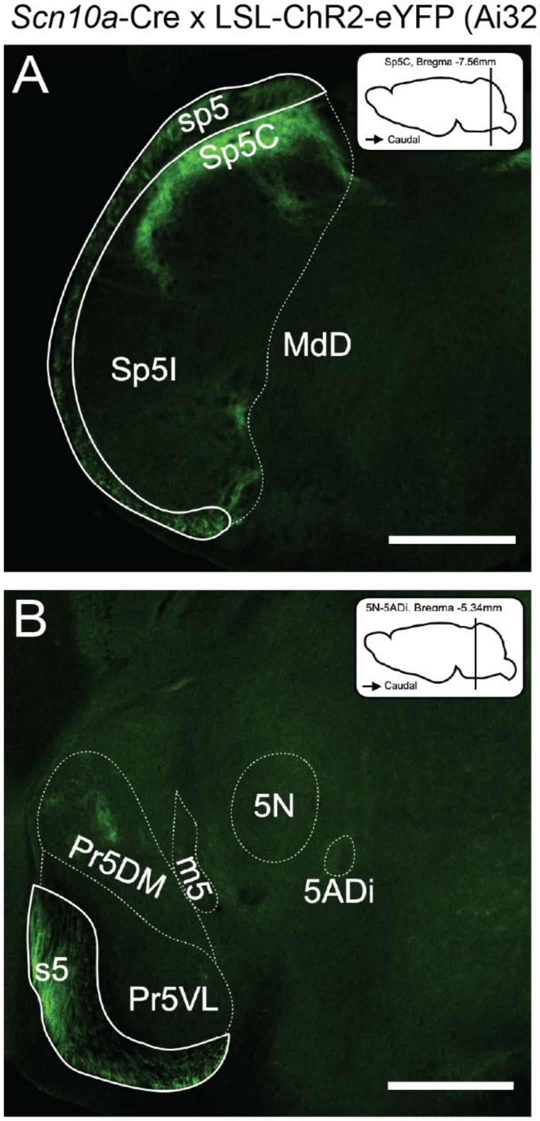


**Figure S13.** ***Scn10a*+ projections directly target the spinal trigeminal nucleus, not the trigeminal motor nuclei. Related to Figure 5.**

Panels A and B demonstrate coronal sections aligned with the corresponding planes indicated by a cartoon brain diagram based on the Allen Mouse Brain Atlas. The sections are positioned at bregma -7.56 mm (A) and bregma -5.34 mm (B). Green fluorescence marks *Scn10a*+ nerve fibers. n = 3 mice were utilized for anatomical mapping and confirmation of intradental nerve fiber projections.

**(A)** Image of coronal section showing the caudal part of the spinal trigeminal nucleus (Sp5C). *Scn10a*+ innervation corresponds to eYFP expression.

**(B)** Image of coronal section of the motor trigeminal nucleus (5N) or the anterior digastric part of the motor trigeminal nucleus (5ADi). *Scn10a*+ innervation corresponds to eYFP expression.

Abbreviations (Allen Brain Atlas): Spinal trigeminal tract (sp5); Spinal trigeminal nucleus, caudal part (Sp5C); Spinal trigeminal nucleus, interpolar part (Sp5I); Medullary reticular nucleus, dorsal part (MdD); Sensory root of trigeminal nerve (s5); Principal sensory trigeminal nucleus, dorsomedial part (Pr5DM); Principal sensory trigeminal nucleus, ventrolateral part (Pr5VL); Motor root of the trigeminal nerve (m5); Motor trigeminal nucleus (5N); Motor trigeminal nucleus, anterior digastric part (5ADi). Scale bar = 500 μm.
